# Bet hedging is not sufficient to explain germination patterns of a winter annual plant

**DOI:** 10.1101/2022.09.15.508102

**Authors:** Gregor-Fausto Siegmund, David A. Moeller, Vincent M. Eckhart, Monica A. Geber

## Abstract

Bet hedging consists of life history strategies that buffer against environmental variability by trading off immediate and long-term fitness. Delayed germination in annual plants is a classic example of bet hedging, and is often invoked to explain low germination fractions. We examined whether bet hedging explains low and variable germination fractions among 20 populations of the winter annual plant *Clarkia xantiana* ssp. *xantiana* that experience substantial variation in reproductive success among years. Leveraging 15 years of demographic monitoring and 3 years of field germination experiments, we assessed the fitness consequences of seed banks and compared optimal germination fractions from a density-independent bet-hedging model to observed germination fractions. We did not find consistent evidence of bet hedging or the expected trade-off between arithmetic and geometric mean fitness, though delayed germination increased long-term fitness in 7 of 20 populations. Optimal germination fractions were 2 to 5 times higher than observed germination fractions, and among-population variation in germination fractions was not correlated with risks across the life cycle. Our comprehensive test suggests that bet hedging is insufficient to explain the observed germination patterns. Understanding variation in germination strategies will likely require integrating bet hedging with complementary forces shaping the evolution of delayed germination.

## Introduction

Organisms across the tree of life exhibit life-history strategies that allow persistence in the face of environmental uncertainty. For annual plants, interannual variation in reproductive success driven by environmental variation can favor the evolution of delayed germination that establishes soil seed banks. Seed banks not only buffer plant populations against environmental change and stochasticity (Eager et al. 2014; Paniw et al. 2017), but also increase effective population size (Nunney 2002; Waples 2006), and maintain genetic diversity (McCue and Holtsford 1998). Theory thus suggests that seed banks have key ecological and evolutionary consequences (Evans and Dennehy 2005).

Evolutionary ecologists have long interpreted delayed germination, caused by persistent or variable seed dormancy, as a bet hedging strategy (Bulmer 1984; Cohen 1966; Ellner 1985*a*,*b*; Philippi and Seger 1989; Simons 2011). Bet hedging increases geometric mean fitness by reducing variability in reproductive success, even if it decreases the arithmetic mean fitness (Seger and Brockman 1987). At the level of individuals, this trade-off between fitness mean and variance is the product of a single genotype that expresses phenotypic variance (Philippi and Seger 1989; Seger and Brockman 1987). Relative to a genotype without dormant seeds, a bet hedging genotype with dormant seeds may have lower fitness in years when all seedlings successfully set seed because only a fraction of the bet hedging genotype’s seeds contribute to next year’s population. However, geometric mean fitness is multiplicative and sensitive to variability in reproductive success across years. A seed bank prevents the bet hedging genotype’s extinction if there is any chance of complete reproductive failure. Genotypes without delayed germination would be lost. The value of delayed germination depends on risk throughout the life cycle. High seed mortality in the seed bank makes it risky for seeds to remain in the soil and selects against delayed germination (Brown and Venable 1986; Cohen 1966; Donohue et al. 2010). Germinating and setting seed can also be risky if seeds experience high mortality after seed set, but before there is an opportunity to germinate or enter the seed bank (Brown and Venable 1991). Ultimately, the individual-level advantage of bet hedging translates to the population-level by increasing long-term population growth rates and persistence.

Some empirical studies have demonstrated that delayed germination, relative to a strategy with complete germination, meets the criteria for bet hedging (Clauss 1999; Evans et al. 2007; Gremer and Venable 2014; Kalisz and McPeek 1993). Specifically, these studies identify the following population-level patterns: (1) reduced arithmetic mean fitness but (2) lower variance in fitness (Clauss 1999), (3) higher long-term stochastic population growth rate (Kalisz and McPeek 1993), or all three at once (Evans et al. 2007; Gremer and Venable 2014). Some degree of delayed germination should be favored whenever there is a nonzero probability of complete reproductive failure (Cohen 1966). If delayed germination functions as a bet hedging strategy that maximizes geometric mean fitness, the optimal germination fraction is expected to evolve in response to levels of seed mortality and temporal variability in reproductive success (Cohen 1966; Franch-Gras et al. 2017; Pinceel et al. 2021). Strong tests of whether germination fractions are adaptive bet hedging strategies thus compare observed and optimal germination fractions, taking into account the complete life history (Childs et al. 2010; Simons 2011). For example, interspecific comparisons of species in a community of Sonoran Desert winter annual plants demonstrated that lower germination fractions were adaptive for species with low seed mortality and with high variability in reproductive success (Gremer and Venable 2014).

Species exhibit substantial variation in germination fractions among populations (e.g., Fernández-Pascual et al. 2013; Gremer et al. 2020; Torres-Martınez et al. 2017), and explaining intraspecific variation in germination strategies remains an open area of inquiry. Studies that have searched for relationships between germination fractions and measures of environmental variability among populations have failed to show that intraspecific variation in germination fractions reflects adaptive bet hedging (e.g., Clauss and Venable 2000; Philippi 1993*b*). However, we are not aware of studies that focus on the relationship between delayed germination and temporal variability in reproductive success among populations. Directly linking germination fractions to fitness consequences, rather than to proxies for environmental variability, may be particularly important for population comparisons because populations are typically distributed across complex environmental gradients that encompass multiple abiotic and biotic variables (Buckley and Puy 2022). Estimates of temporal variability in reproductive success are thus crucial to test whether delayed germination functions as bet hedging in populations, and to assess whether differences in germination fractions among populations are adaptive (Simons 2011).

Populations of the winter annual, *Clarkia xantiana* ssp. *xantiana*, are distributed across a complex landscape in the southern Sierra Nevada (Fig. 1A; Eckhart et al. 2011; Gould et al. 2014). Although early research using seed sowing experiments suggested that the species lacked a seed bank (Lewis 1962), multiple lines of evidence now demonstrate the presence and importance of a seed bank in *C. x.* ssp. *xantiana*. In field experiments, seeds can germinate at least up to three years after being buried in bags (Eckhart et al. 2011) or pots (M. A. Geber, *unpublished data*). Fifteen years of field surveys suggest that the seed bank allows some populations to persist exclusively as seeds for as long as four consecutive years (Fig. 1B). Seeds can also remain viable for up to 11 years when buried in bags 30 cm below the soil surface (D. A. Moeller, *unpublished data*). *Clarkia xantiana* ssp. *xantiana* seeds lack morphological adaptations for dispersal (Knies et al. 2004), and the species’ small-scale spatial distribution is consistent with dispersal limitation (Kramer et al. 2011). We thus expect a limited role for dispersal to complement delayed germination under temporal variability (Venable and Brown 1988).

**Figure 1:**
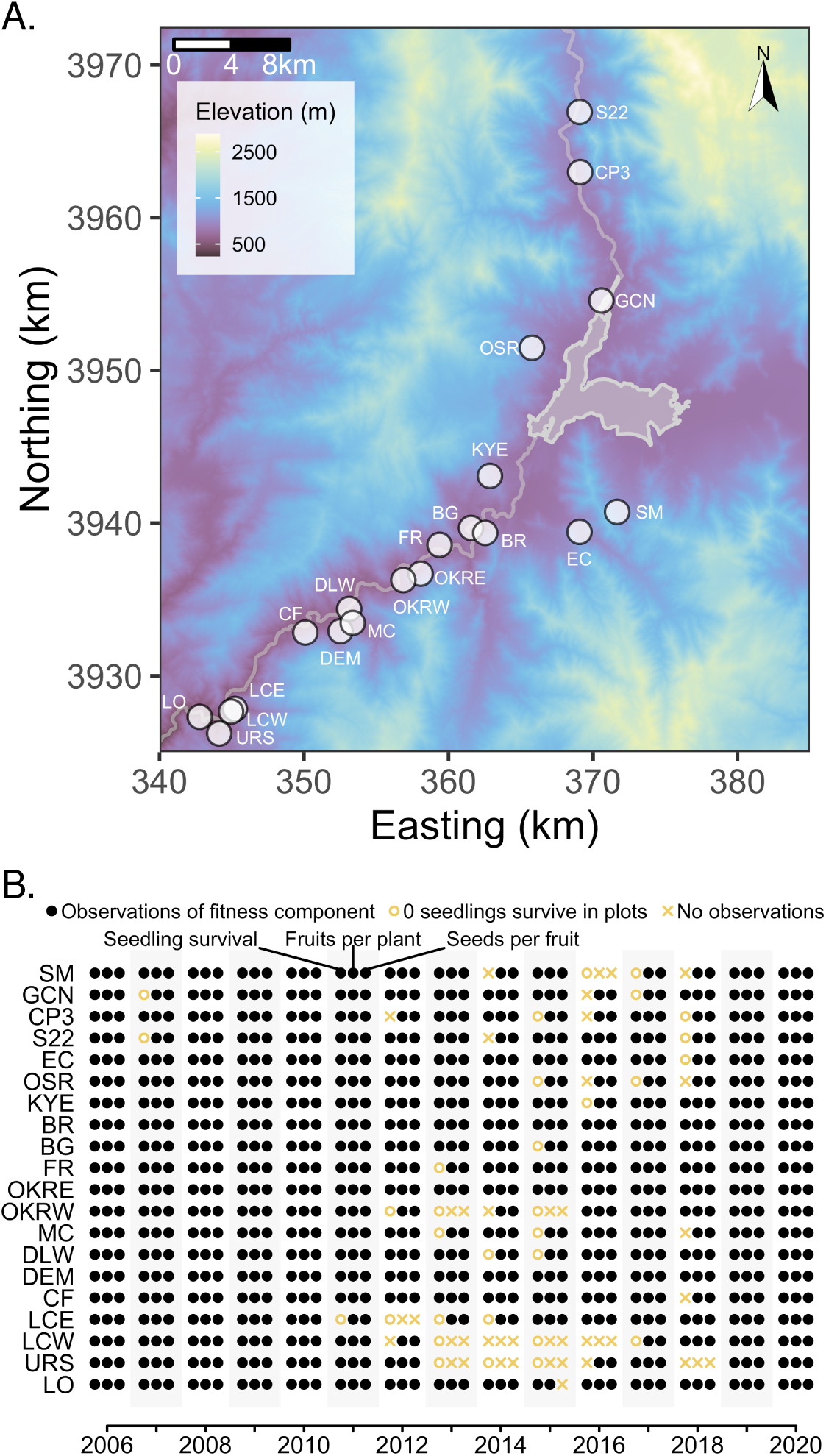
Map of the populations, and summary of aboveground observations of demography. (A) Elevation map of study populations. (B) Graphical summary of 15 years of aboveground observations at study populations. Open circles indicate that no seedlings survived in permanent plots; Xs indicate that no seedlings or plants were observed in surveys. Populations are arrayed from west (bottom) to east (top).

Delayed germination is ubiquitous in *C. x.* ssp. *xantiana* and, quantitatively, the strength of delayed germination varies among populations (Eckhart et al. 2011). Specifically, Eckhart et al. (2011) found that all 20 populations they studied exhibited delayed germination and that the germination fractions of first-year seeds increased from west to east, roughly doubling from around 10% in western populations to over 20% in eastern populations. We ask whether delayed germination in these populations functions as bet hedging and whether the germination fractions in each population are adaptive, given the temporal variability in reproductive success in each population. Complex spatial and temporal variation in abiotic (e.g., soil texture: Eckhart et al. 2010; temperature and precipitation: Eckhart et al. 2011) and biotic (e.g., pollinators: Moeller 2004; mammalian herbivores: Benning et al. 2019; insect herbivores: D. A. Moeller, *unpublished data*) variables affect individual and population performance. We do not focus on the relationship between delayed germination and any particular environmental variable, but instead leverage a long-term demographic study to directly quantify temporal variability in reproductive success.

Here, we tested whether observed germination fractions and life-history patterns in *Clarkia xantiana* ssp. *xantiana* are consistent with predictions made by bet hedging models. To connect germination and its fitness consequences, we combined 15 years of observations on reproductive success and three years of seed burial experiments from 20 populations to address the following questions. (1) Does delayed germination and the formation of a seed bank meet the criteria for bet hedging? Specifically, for each population, we tested whether delayed germination decreases arithmetic mean fitness, reduces the variability in fitness, and increases the long-term stochastic population growth rate. Next, we tested whether the observed germination fractions are likely to be adaptive. (2) For each population, does the optimal germination fraction predicted by bet hedging models match observed germination fraction? We found that life-history patterns are not entirely consistent with bet hedging expectations. We thus examined the relationship between germination fraction and risk, both by seeds before germination, and by seedlings after germination. Under bet hedging, we expected a negative correlation between germination fraction and risk, so we specifically asked the following questions: (3) Is there a negative correlation between germination fraction and seed survival across populations? (4) Is there a negative correlation between germination fraction and variability in per capita reproductive success across populations?

## Methods

### Clarkia xantiana ssp. xantiana life history

*Clarkia xantiana* (A. Gray) ssp. *xantiana* (Onagraceae) is a winter annual that germinates with late fall and winter rains, and sets seeds during the summer drought, in California’s Mediterranean climate. In our study region, the Kern River Canyon and Valley (Kern and Tulare Counties, California, U.S.A.), germination happens from November through March. Seedlings grow in winter and spring, and surviving plants flower in late spring and early summer, late April into mid-June. Pollinated fruits set seed in the early summer, June to July, and fruits subsequently dry out and gradually split open. Most seeds appear to be shed from fruits within three to four months after seed set, but can remain on the plant for more than a year. Seeds are small (*<* 1 mm in width) and have no structures to aid in aerial or other dispersal. After seed set in June/July, these new seeds survive to the subsequent winter before germinating or entering the soil seed bank.

We represent the *C. x.* ssp. *xantiana* life history in terms of transitions from October of year *t* to October of year *t* + 1. Transitions are the product of seed survival and germination, and aboveground seedling survival to fruiting, fruit production, and seeds per fruit. For this study, we assume that new seeds and seeds in the soil seed bank have different survival rates, but we assume that germination rates are the same regardless of seed age. We also assume that all plants experience the same vital rates upon germination; in other words, vital rates are not a function of plant size. We describe population growth rate by the following equation:

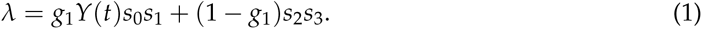

Germination is given by *g*_1_. Per capita reproductive success in year *t*, *Y*(*t*) is the product of seedling survival to fruiting, fruits per plant, and seeds per fruit. Seed survival from seed set in June/July to the first October is *s*_0_. Seed survival from the first October to germination in January/February is *s*_1_. For new seeds, seed survival from June/July to January/February is thus *s*_0_*s*_1_. Survival of ungerminated seeds from January/February to the next October is *s*_2_. Seed survival from October to the second germination opportunity the following January/February is *s*_3_. For seeds in the seed bank, seed survival from January/February in one year to the next is thus *s*_2_*s*_3_. All parameters are summarized in Table 1.

**Table 1:** Vital rate components of the structured population model.

| Component | Description | Data contributing to quantity |
| --- | --- | --- |
| **SEED SURVIVAL** |  |  |
| $s_0$ | Probability that a seed produced in July of year $t$ is intact and viable in October of year $t$ | Seed bag burial experiment, viability trials, seedling counts in permanent plots, fruiting plant counts in permanent plots, fruit per plant counts, seeds per fruit counts |
| $s_1$ | Probability that a seed survives from October of year $t$ to January of year $t + 1$, for seeds produced in year $t$ | Seed bag burial experiment, viability trials |
| $s_2$ | Probability that a seed survives from January of year $t + 1$ to October of year $t + 1$, for seeds produced in year $t$ | Seed bag burial experiment, viability trials |
| $s_3$ | Probability that a seed survives from October of year $t + 1$ to January of year $t + 2$, for seeds produced in year $t$ | Seed bag burial experiment, viability trials |
| **GERMINATION** |  |  |
| $g_1$ | Probability of germination for a seed that has survived to January of year $t + 1$, for seeds produced in year $t$ | Seed bag burial experiment, viability trials |
| **PER CAPITA REPRODUCTIVE SUCCESS** |  |  |
| $\sigma$ | Probability of seedling survival to fruiting, from a January/February census through reproduction in June/July | Seedling count in permanent plots, fruiting plant count in permanent plots |
| $F$ | Number of fruits per fruiting plant | Fruit counts on plants in permanent plots, fruit counts on plants in additional plots, seeds per fruit counts on plants in additional plots |
| $\phi$ | Number of seeds per fruit | Seeds per fruit counts on plants in additional plots |

### Creating the dataset

We used field surveys and experiments to assemble observations of above- and below-ground demography for 20 populations of *C. x.* ssp. *xantiana* across its range (Table 2, S1). A subset of the demographic data has been used to test hypotheses about geographic variation in population growth rate and species distributions (Eckhart et al. 2011; Pironon et al. 2018). Here, we used field surveys to collect data on seedling survival, fruit production, and seed set. We also conducted field experiments to observe emergence of seedlings and seeds remaining intact in the soil seed bank. We used the data from the surveys and experiments to fit statistical models for the demographic parameters that describe the life cycle (Equation 1). Ultimately, we used the parameter estimates from these statistical models to calculate per capita reproductive success, seed survival, and germination to test predictions of bet hedging models.

**Table 2:** Summary of observations and experiments

| Parameter data | Description | Time span |
| --- | --- | --- |
| SEED VITAL RATES | — | — |
| Seed survival and germination | Seed bag burial | 2005-2008 |
| Seed viability | Viability trials | 2005-2008 |
| SEEDLING SURVIVAL | — | — |
| Seedling survival to fruiting | Field surveys | 2006-2020 |
| FRUITS PER PLANT | — | — |
| Total fruit equivalents per plant | Field surveys | 2006-2012 |
| Undamaged and damaged fruits per plant | Field surveys | 2013-2020 |
| Total fruit equivalents per plant | Extra plots | 2006-2012 |
| Undamaged and damaged fruits per plant | Extra plots | 2013-2020 |
| SEEDS PER FRUIT | — | — |
| Seeds per undamaged fruit | Lab counts | 2006-2020 |
| Seeds per damaged fruit | Lab counts | 2013-2020 |

#### Field surveys for aboveground components of demography

We conducted field surveys of seedlings, fruiting plants, fruits per plant, and seeds per fruit at two spatial scales (Figure 2A; Eckhart et al. 2011). First, in October 2005, we established 30 1 *×* 0.5 m^2^ permanent plots at each of the 20 study populations. The permanent plots were arrayed across four to six transects per population, and each plot was 2.5 m apart along a transect. Permanent plots were used for annual surveys of seedlings, fruiting plants, and fruits per plant. Second, additional, haphazardly distributed 1 *×* 0.5 m^2^ plots were used each year to supplement counts of fruits per plant from permanent plots, and to identify plants for fruit collection. Finally, when we found no or few plants in the haphazardly distributed plots, we also searched the population for additional plants to count fruits per plant and from which to collect fruits. By collecting fruits from plants outside the permanent plots, we did not affect seed input into the permanent plots. To estimate the survival of seedlings to fruiting plants, we counted seedlings (*n_ijk_*) and fruiting plants (*y_ijk_*) in each permanent plot each year from 2006–2020. Seedlings and fruiting plants were counted in January/February and June, respectively, in plot *i*, year *j*, and population *k*.

**Figure 2:**
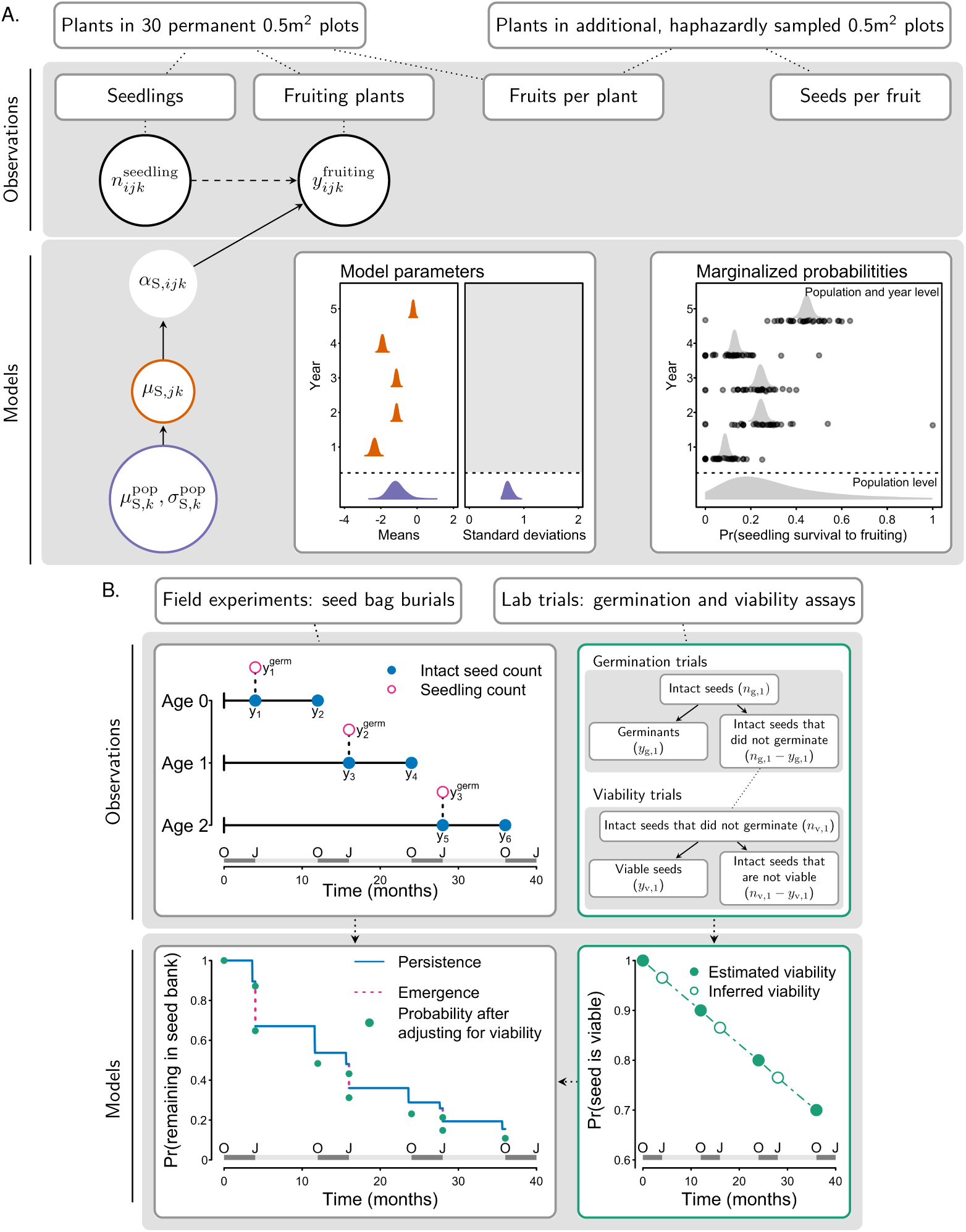
Graphical summary of the observations, models, and parameters used to estimate per capita reproductive success, germination, and seed survival. (A) A graphical representation of the relationship between the structure of observations and the data. A directed acyclic graph showing the model for seedling survival to fruiting, with colors corresponding to the simulated example in the plots showing the relationship between model parameters, marginalized probabilities, and data. The statistical model for seedling survival to fruiting is presented in Appendix S2.2.1. (B) A graphical representation of the field seed bag experiments and lab viability trials. In the panel showing the seed bag burial experiments, we show counts of germinants, 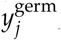, in year *j* of the experiment and counts of intact seeds, *y_m_*, at the *m*th observation. The statistical model for observations from the seed bag burial experiment is presented in Appendix S2.2.3. The statistical model for observations from the lab viability trials is presented in Appendix S2.2.4.

Of more than 8000 observations, there were fewer seedlings than fruiting plants in approximately 5% of observations; 50% of these had 1 fewer seedling than fruiting plant (Table S5). There are at least two possible sources of undercounts of seedlings. An observer might miss small seedlings that were present at the January/February seedling census, or additional seedlings emerged after the census. We assume that we did not under- or over-count fruiting plants because plants stand out from the background vegetation in June. To account for the undercount of seedlings, we recoded the data so that the count of seedlings was equal to the number of fruiting plants observed later in the season.

To determine the number of fruits per plant, we counted the number of fruits per plant on up to 15 plants in each of the permanent plots from 2007–2020, and on additional plants in the haphazardly distributed plots from 2006–2020 (Figure 2A). We combined counts from plants in permanent and haphazardly distributed plots, because the latter often sampled a broader distribution of plant sizes and combining them allowed us to better estimate fruit number per plant in years with relatively few plants in permanent plots.

From 2006–2012, we counted the number of undamaged fruits on a plant. We then took the damaged fruits on a plant and visually stacked them end to end to estimate how many additional undamaged fruits that was equivalent to (e.g., two half fruits corresponded to one undamaged fruit). We used this as our count (*y_ijk_^TFE^*) of total fruit equivalents on plant *i*, in year *j*, and in population *k*. From 2013–2020, we separately recorded the number of undamaged (*y_ijk_^UF^*) and damaged (*y_ijk_^DF^*) fruits on a plant.

From 2006–2020, we counted the number of seeds in one undamaged fruit (*y_ijk_^US^*) collected from each of 20-30 plants in or outside of the haphazardly distributed plots. Our counts corresponded to fruit *i*, in year *j*, and in population *k*. From 2013–2020, we also counted the number of seeds in one damaged fruit (*y_ijk_^DS^*) collected from each of 20-30 plants in or outside of the haphazardly distributed plots.

#### Field experiments for belowground components of demography

We conducted a field experiment to estimate seed persistence from fall (October) to winter (January/February), emergence in the winter, and seed persistence from winter to fall (Fig. 2B). At each population, we buried seeds in mesh bags in the fall before the onset of winter rains, counted intact seeds and seedlings in a subset of bags in the winter, and then retrieved those bags the following fall to count intact seeds and conduct a two-stage lab trial to assay viability of intact seeds. Seeds that were intact in the field may have been dormant (and viable) or dead but not decayed (and not viable). To estimate seed persistence and emergence, we used data from field experiments; these estimates do not account for loss of seed viability. To estimate seed survival and germination, we combined the field and lab experiments; these estimates account for loss of seed viability.

The experiment consisted of three rounds starting in October 2005, 2006, or 2007. For each round, we collected seeds at each population in summer before the round started. For each population, we pooled and distributed seeds across 5*×*5-cm nylon mesh bags (100 seeds/bag). In October, we returned the bags to the population at which the which seeds were collected, staked one bag near each permanent plot (Methods: Field surveys for aboveground demography) and covered the bags with soil.

In Round 1, we placed 30 bags at each population in October 2005. We unearthed a first set of 10 bags in January 2006 to count the number of intact seeds (*y*) and the number of seedlings (*y_g_*) (Age 0 in Fig. 2B). We returned the bags to the ground until October 2006, when we retrieved bags to the lab to count intact seeds (*y*) and test seed viability (see below). In the second year of Round 1, we counted intact seeds and seedlings in a second set of 10 bags unearthed in January 2007 (Age 1 in Fig. 2B). We again returned these bags to the ground until October 2007, when we retrieved these 10 bags to count intact seeds and test seed viability. In the third year of Round 1, a third set of 10 bags was unearthed in January 2008 to count intact seeds and seedlings (Age 2 in Fig. 2B), and brought to the lab in October 2008 for seed counts and viability tests.

The experiment was repeated in all populations two more times. Round 2 started in October 2006 with 20 bags per population, and 10 bags were dug up in the first and second year (2007 and 2008, respectively). Round 3 started in October 2007 with 10 bags per population, and 10 bags each were dug up after one year (2008). We thus made three sets of observations associated with age 0 seeds (brought to the lab after one year in the field), two sets of observations associated with age 1 seeds (brought to the lab after two years in the field), and one set of observations associated with age 2 seeds (brought to the lab after three years in the field).

In October of each experimental year, the seeds remaining intact in the subset of bags that were brought to the lab were counted and tested for viability in a two-stage trial (Fig. 2B). We placed up to 15 seeds from each bag on moist filter paper in a disposable cup; over a 10-day span, we counted and removed germinants every two days. Because we conducted two to three tests of 15 seeds each per bag, we summed the number of seeds tested (*n_g_*^viab^) and germinating (*y_g_*^viab^) to summarize the trials and successes.

After 10 days, up to 10 remaining ungerminated seeds per cup were sliced in half and individually placed into 96-well plates filled with a solution of tetrazolium chloride, which stains viable tissue red. We covered the plates with foil. Each 96-well plate contained seed from at least one bag per population of a given seed-age class. We counted viable seeds every two days for 10 days. For each bag, we summed the number of seeds tested (*n_v_*^viab^) and staining (*y_v_*^viab^) to summarize the trials and successes.

### Statistical models

We used observational and experimental data from 20 populations to fit statistical models for the demographic parameters that describe the life cycle (Fig. 2). We refer readers to Appendix S2 for a description of the statistical models, directed acyclic graphs, and for the mathematical expressions for the posterior proportional to the joint distribution for all the models.

#### Aboveground components of demography

We used a hierarchical, Bayesian approach to fit models to observations of seedling survival, fruits per plant, and seeds per fruit. As an example, we describe the structure of the model for seedling survival to fruiting, which is essentially a generalized linear mixed model with a binomial likelihood and a logit link (Fig. 2A). We use directed acyclic graphs (DAGs) to illustrate the relationship between the observations, the model, and parameters of interest. In the field, we counted seedlings (*n_ijk_*^seedlings^) and fruiting plants (*y_ijk_*^fruiting^) in plot *i*, year *j*, and population *k*. These quantities are outlined in black in the DAG and are shown as black points in the corresponding graphs. The model uses a binomial likelihood and relates the data to a probability of survival, *α*_S_. This parameter is logit-transformed and links the year-level distribution, outlined in orange, to the observations. Parameters for the year-level distribution are annual estimates of the mean, which are drawn from the population-level distribution, outlined in purple. We write the model using hierarchical centering to account for the structure of our observations and for computational efficiency (Evans et al. 2010; Ogle and Barber 2020), but it is equivalent to a random effects structure in which years are nested within populations. For each set of observations, we fit separate models to each population so that the resulting annual estimates were partially pooled towards the population-level mean.

The models for fruits per plant and seeds per fruit have a similar hierarchical structure but use Poisson likelihoods and a log link (Appendix S2.2.2). We separately modeled observations of total fruit equivalents per plant for 2006–2012 and total fruits per plant for 2013–2020. In years with observations of total fruits per plant, we used counts of undamaged and damaged fruits per plant to model the proportion of fruits that were damaged. We estimated seeds per undamaged fruit for 2006–2020, and combined those estimates with counts of seeds per damaged fruit to infer the proportion of seeds that were lost to herbivory for 2013–2020. To make the two sets of observations for fruits per plant compatible, we used the proportion of fruits per plant that were damaged and the proportion of seeds lost to herbivory on a damaged fruit to calculate total fruit equivalents per plant from 2013–2020.

We fit hierarchical, Bayesian models to our data for several reasons. First, hierarchical models perform well for making inferences about annual variation in demography (Metcalf et al. 2015). Second, the study period included substantial variation in sample size (Tables S2-S4, S6-S9), including years in which we did not observe plants in permanent plots even when they were present in the broader population (Fig. 1B). Hierarchical models for seedling survival introduce partial pooling, which allowed us to account for sampling variation in fitting the model rather than post-hoc. Third, our approach made it straightforward to quantify uncertainty associated with annual estimates of components of reproductive success. Fourth, estimating germination and seed survival from the seed bag experiment required combining three datasets (see below), a process that is a strength of Bayesian methods (Hobbs and Hooten 2015).

#### Belowground components of demography

Estimating seed survival and germination from the seed experiment required combining datasets. Here, we describe and graphically illustrate the model that we fit to observations from field experiments (Fig. 2B). The model we fit to the observational data jointly accounts for loss of seeds from the seed bank through mortality and germination (Siegmund and Geber 2022). Germination occurs once a year in the winter, and is estimated from the seeds that germinate each year. Mortality occurs throughout the year, and is estimated from the seeds that remain intact. In Fig. 2B, the model describes the stairstep shape of the curve in the lower left panel. In practice, we fit a survival function that is the product of discrete germination and mortality hazards (Klein and Moeschberger 2003).

Separately, we obtained viability of seeds using the two-stage lab trials. Each lab trial consisted of two binomial experiments that measured (1) germination of intact seeds and then (2) viability of seeds that did not germinate. We combined these estimates to infer viability in each population and year. The lab trials involved destructive sampling, and we only conducted them when bags were retrieved in October (filled points in lower right panel of Fig. 2B). We inferred the viability of intact seeds in January by assuming that seeds lost viability at a constant rate (exponential decay). Further, we interpolated between estimates by assuming that viability changed at a constant rate across years, and that all seeds were viable at the start of the experiment (open points in lower right panel of Fig. 2B).

Finally, because plants set seed in July but the field experiments with seed bags did not start until October, we did not have direct observations to inform estimates of *s*_0_, the probability of seed survival from seed set in July to four months later in October. To infer seed survival during this part of the life cycle, we combined data from the field surveys and seed bag experiments (Elderd and Miller 2016). We assumed that the seedlings emerging in permanent plots in 2008 were primarily from seeds produced in permanent plots in the previous two years, 2006 and 2007, that survived to and germinated in 2008. We ignored contributions from older seeds, assuming for simplicity that they make up a small proportion of seedlings. We used counts of fruiting plants in the permanent plots, and estimates of seed set per fruiting plant, to calculate the average seed set per transect in 2006 and 2007. We then linked seed set, and estimates of seed survival and germination from the seed bag burial experiment, to the average number of seedlings observed in permanent plots. Once we joined these observations, we inferred *s*_0_ as the proportion of seeds lost between seed set in July and October.

### Model statements, implementation, and fitting

We show the expressions for the posterior proportional to the joint distribution, and corresponding directed acyclic graphs, for all models in Appendix S2. Prior choice is described in Appendix S3, and Table S10 shows all parameters with associated priors. We prepared data for analysis using the tidyverse (Wickham and RStudio 2021) and tidybayes (Kay and Mastny 2020) packages in R version 3.6.2 (R Core Team 2019). We wrote, fit all models, and estimated posterior distributions using JAGS 4.10 with rjags (Plummer et al. 2019). We used the MCMCvis package to work with the model output, check chains for convergence, and recover posterior distributions (Youngflesh et al. 2021). We randomly generated initial conditions for all parameters with a prior by drawing from the corresponding probability distribution in R before passing the initial values to rjags. We ran three chains for 45,000 iterations. The first 10,000 iterations were for adaptation, the next 15,000 iterations were discarded as burn-in, and we sampled the following 15,000 iterations. We assessed convergence of the MCMC samples with visual inspection of trace plots, by calculating the Brooks-Gelman-Rubin diagnostic, R̂, and by calculating the Heidelberg-Welch diagnostic (Appendix S4).

### Computing vital rates

In the following sections, we describe how we used estimates from the statistical models to obtain the parameters that describe the *C. x.* ssp. *xantiana* life history. To calculate variation in per capita reproductive success for the study populations, we obtained annual estimates for seedling survival to fruiting, fruits per plant, and seeds per fruit from the field surveys. Because our goal was to compare patterns of seed bank dynamics among populations, we obtained population-level estimates for germination and seed survival from the seed bag burial experiment. The calculations summarized here are described in detail in Appendix S5.

#### Per capita reproductive success

We calculated annual per capita reproductive success as the number of seeds produced per seedling each year, on average (Gremer and Venable 2014; Venable 2007). In other words, it is the product of the annual mean probabilities of seedling survival to fruiting, fruits per plant, and seeds per fruit. We calculated the posterior mode of annual estimates for each of the vital rate parameters in each year (the orange distribution in Fig. 2A) and multiplied them to obtain the per capita reproductive success in that year. By using vital rates estimated for the same year to calculate per capita reproductive success, we kept the observed covariation among vital rates within years (i.e., kernel sampling with partial pooling, as described in Metcalf et al. 2015). Our estimates thus represented temporal variability in per capita reproductive success over the 15 year study period, and implicitly integrated over environmental variables (e.g., precipitation, temperature, pollination, herbivory) that influence survival, fruit production, and seed set.

To compute per capita reproductive success, we used 15 years of observations from each of 20 populations. Our observations throughout the study period included missing data that reflects natural variability in population size or the spatial distribution of plants at study populations (Fig. 1B). We accounted for missing data while calculating per capita reproductive success. In some years (n=3), we observed no seedlings in permanent plots or fruiting plants in permanent plots, additional haphazard plots or elsewhere across the population. In other years (n=9), we observed seedlings in permanent plots but no fruiting plants anywhere in the population. We assumed that this reflected a true absence of fruiting plants in that year and that there was no seed set in those years, so we set fruits per plant and seeds per fruit to zero. In one year at one population, we observed a single fruiting plant with three fruits, from which we did not collect seeds. For this estimate, we substituted the population average of seeds per fruit. Finally, there were years (n=13 years) when there were no plants in permanent plots but we found plants elsewhere throughout the population. We had no information about seedling survival in these years, and so used the population’s average for seedling survival to fruiting.

#### Belowground vital rates

Estimates from the seed bag burial experiment describe persistence, the probability that a seed remains intact in the seed bank, and emergence, the probability that an intact seed becomes a seedling. To estimate seed survival and germination, which account for loss of seed viability of intact seeds, we combined information from the seed bag burial experiment and the lab trials (Table S11). First, we estimated the probability that seeds persisted, or remained intact, in the seed bank (Fig. 2B). We combined estimates for persistence with the viability estimates to calculate seed survival, the probability that seeds remained intact and viable in the seed bank. Similarly, we combined estimates for emergence with viability to calculate germination, the probability that viable, intact seeds became seedlings. We used the seed survival (*s*_1_, *s*_2_, *s*_3_) and germination (*g*_1_) probabilities to test predictions from bet hedging theory. Because seed survival from seed set in July to October (*s*_0_) implicitly included loss of seed viability, we did not adjust these estimates.

### Analysis

#### Demographic test of bet hedging

We used estimates for the vital rate components to test whether delayed germination is an adaptive bet hedging trait in *C. x.* ssp. *xantiana*. The life history described by Equation 1 incorporates a seed bank. Specifically, populations form a seed bank by delaying germination (i.e. *g*_1_ *<* 1). Immediate germination (*g*_1_ = 1) eliminates the seed bank, in which case Equation 1 reduces to:

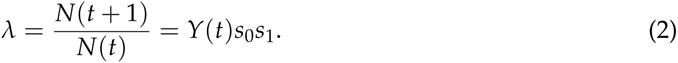

We calculated per capita reproductive success, *Y*(*t*), as the product of seedling survival to fruiting, fruits per plant, and seeds per fruit. We tested whether delayed germination (*g*_1_ *<* 1) functions as bet hedging by eliminating the seed bank (Equation 2). If delayed germination is consistent with bet hedging, we expected eliminating the seed bank to increase arithmetic mean fitness, increase the variability in fitness, and decrease the long-term stochastic population growth rate (Clauss 1999; Evans et al. 2007).

To calculate the arithmetic mean population growth rate, we used the average of annual population growth rates, *λ_a_* (Evans et al. 2007). We obtained values for the average population growth rate with the field estimates of germination as well as with the seed bank eliminated (*g*_1_ = 1). We used the posterior modes for annual estimates of per capita reproductive success, *Y*(*t*), and for population-level estimates of seed mortality and germination. We assumed that each estimate for per capita reproductive success was equally likely, calculated annual population growth rates with (Equation 1) and without (Equation 2) a seed bank, and computed *λ_a_* as the average.

To calculate temporal variability in population growth rate, we drew 1,000 samples from the 15 years of per capita reproductive success estimates with replacement. We paired these resampled years of estimates with the population-level values for germination and seed survival rates to calculate annual population growth rates. For both the case with and the case without a seed bank, we calculated the variance of the sequence of population growth rates.

To calculate the long-term stochastic population growth rate, we used the same sequence of population growth rates that we used to calculate temporal variability in fitness. We calculated the long-term stochastic population growth rate with the field estimates of germination, as well as with the seed bank eliminated (*g*_1_ = 1). We used the following equation to calculate the stochastic population growth rate:

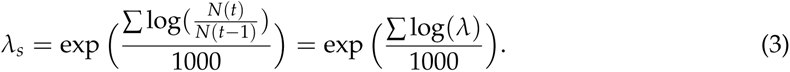

To examine the effect of uncertainty about parameter estimates on our results, we used the full posterior distribution for parameter estimates to calculate the arithmetic mean growth rate, temporal variability in population growth rate, and long-term stochastic population growth rate (Appendix S6.2). To assess how sensitive our results were to years with complete reproductive failure, we repeated our analysis with quasi-complete germination (*g*_1_ = 0.99) in Equation 1. By including a minimal seed bank, we evaluated support for the trade-off between arithmetic and geometric mean fitness in the case when populations do not go extinct with just one bad year.

#### Density-independent model for germination fractions

We calculated the optimal germination fraction for the observed variation in reproductive success and seed survival. For each population, we used a sequence of 100,000 resampled values for per capita reproductive success, *Y*(*t*), and the observed seed survival probabilities, *s*_0_, *s*_1_, *s*_2_, and *s*_3_, to calculate population growth rates at each germination fraction, *G*, along an evenly spaced grid of values from 0 and 1. Temporal variation was incorporated into the model by resampling per capita reproductive success, *Y*(*t*). The optimal germination fraction is the value of *G* that maximizes the geometric mean of the population growth rate. We found the optimal germination fraction *G* using a one-dimensional optimization algorithm (Brent 1973). To evaluate the relationship between the optimal and observed germination fractions, we calculated the Pearson correlation coefficient between the optimal *G* and the posterior mode of *g*_1_. To assess the influence of parameter uncertainty on optimal germination fractions, we examined how optimal *G* varied when we drew samples from the posterior distribution of each parameter in the population model (Appendix S6.4). For each sample, we found the optimal germination fraction, *G*, using a single optimization with 100,000 years of resampled values for *Y*(*t*).

#### Correlation between germination and seed survival

We tested whether observed germination, *g*_1_ was negatively correlated with seed survival, *s*_2_*s*_3_. Here, seed survival is the probability that seeds which do not germinate in January remain in the seed bank until the following January. We obtained the posterior distribution for the correlation between germination and seed survival by calculating the sample correlation of *g*_1_ and *s*_2_*s*_3_ at each iteration of the MCMC output.

#### Correlation between germination and variability in per capita reproductive success

We tested whether observed germination, *g*_1_ was negatively correlated with the temporal variability in per capita reproductive success for each population. We estimated variability by sampling the posterior distribution of reproductive success for each year and calculating the geometric standard deviation of per capita reproductive success as exp(SD (log (per capita reproductive success+0.5))). We obtained the sample correlation of germination and geometric standard deviation of per capita reproductive success at each iteration of the MCMC output.

## Results

The MCMC chains exhibited good convergence and mixing, which we assessed with the R̂ and Heidelberg-Welch diagnostics (Fig. S7, S8). We used parameter estimates from the statistical models to compute the vital rates for the population models, and we present graphical summaries of all vital rate parameters in Appendix S5.3.

### Demographic test of bet hedging

To determine whether delayed germination meets the criteria for bet hedging in each population, we compared the arithmetic mean population growth rate, variance in population growth rate, and long-term stochastic population growth rate with and without a seed bank. The arithmetic mean growth rate was greater without a seed bank than with a seed bank (Fig. 3A). The variance in population growth rates was also greater without a seed bank than with a seed bank (Fig. 3B, S17). However, the long-term stochastic population growth rate was not always higher with a seed bank (Fig. 3C); the stochastic population growth rate was greater with a seed bank in only 7 out of 20 populations. These results were robust to uncertainty in parameter estimates (Fig. S18). When we assumed quasi-complete germination and a minimal seed bank, the stochastic population growth rate was greater with a seed bank in 4 out of 20 populations (Fig. S19), qualitatively consistent with the results assuming complete germination and no seed bank.

**Figure 3:**
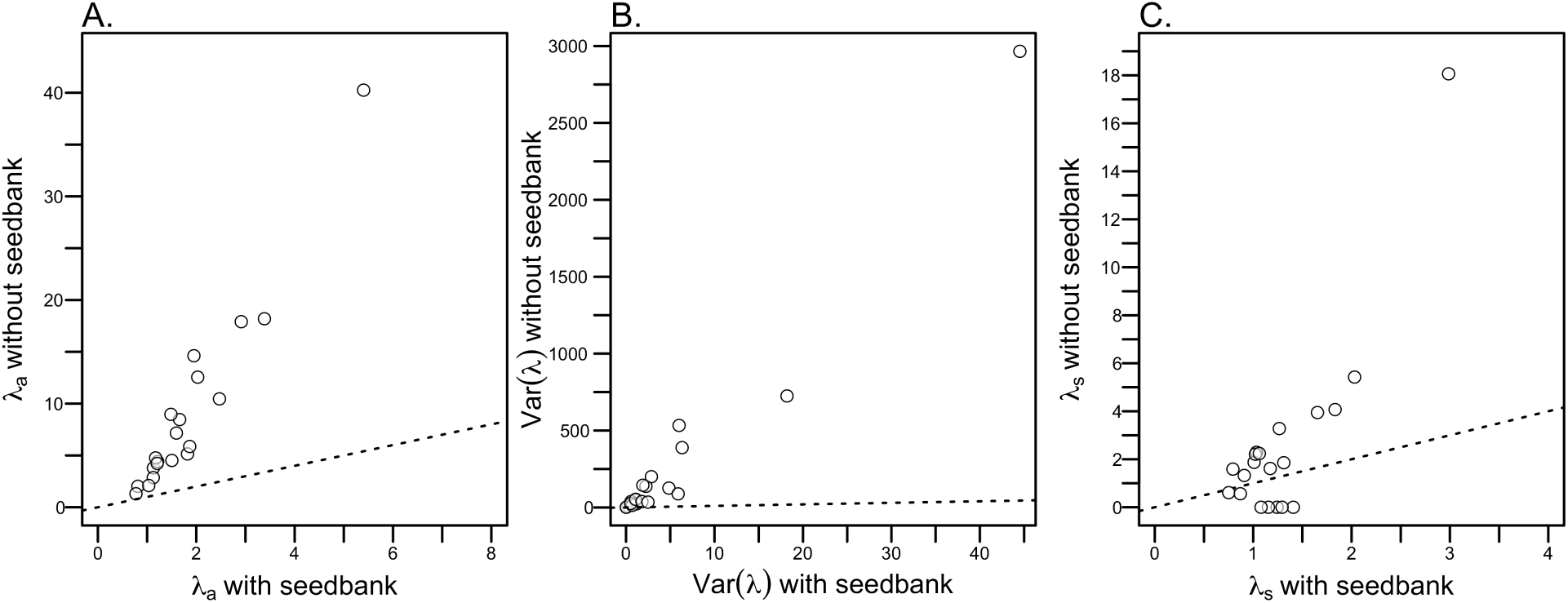
Test of the demographic patterns expected with bet hedging. (A) Plot of the arithmetic population growth rate without a seed bank against arithmetic population growth with a seed bank. (B) Plot of the variance in annual population growth rate without a seed bank against the variance in population growth rate with a seed bank. See Fig. S17 for a plot that zooms in on the lower left hand corner and shows the variability in the variance of population growth rate in that region of the plot. (C) Plot of the long-term stochastic population growth rate without a seed bank against the long-term stochastic growth rate without a seed bank. In all plots, the dotted line is the 1:1 line.

### Observed germination fractions are lower than predicted by a density-independent model

To evaluate the density-independent model, we compared observed germination to predicted germination optima (Fig. 4). Optimal germination fractions were less than one in 13 out of 20 populations (Fig. 4). Optimal and observed germination fractions were uncorrelated (Fig. 4; r=-0.158, p=0.507). Predictions from the density-independent model were higher, often by two- or three-fold, than observed germination fractions. Optimal germination fractions were robust to uncertainty in parameter estimates, but parameter uncertainty produced a wide range of optimal germination fractions for several populations (Fig. S20).

**Figure 4:**
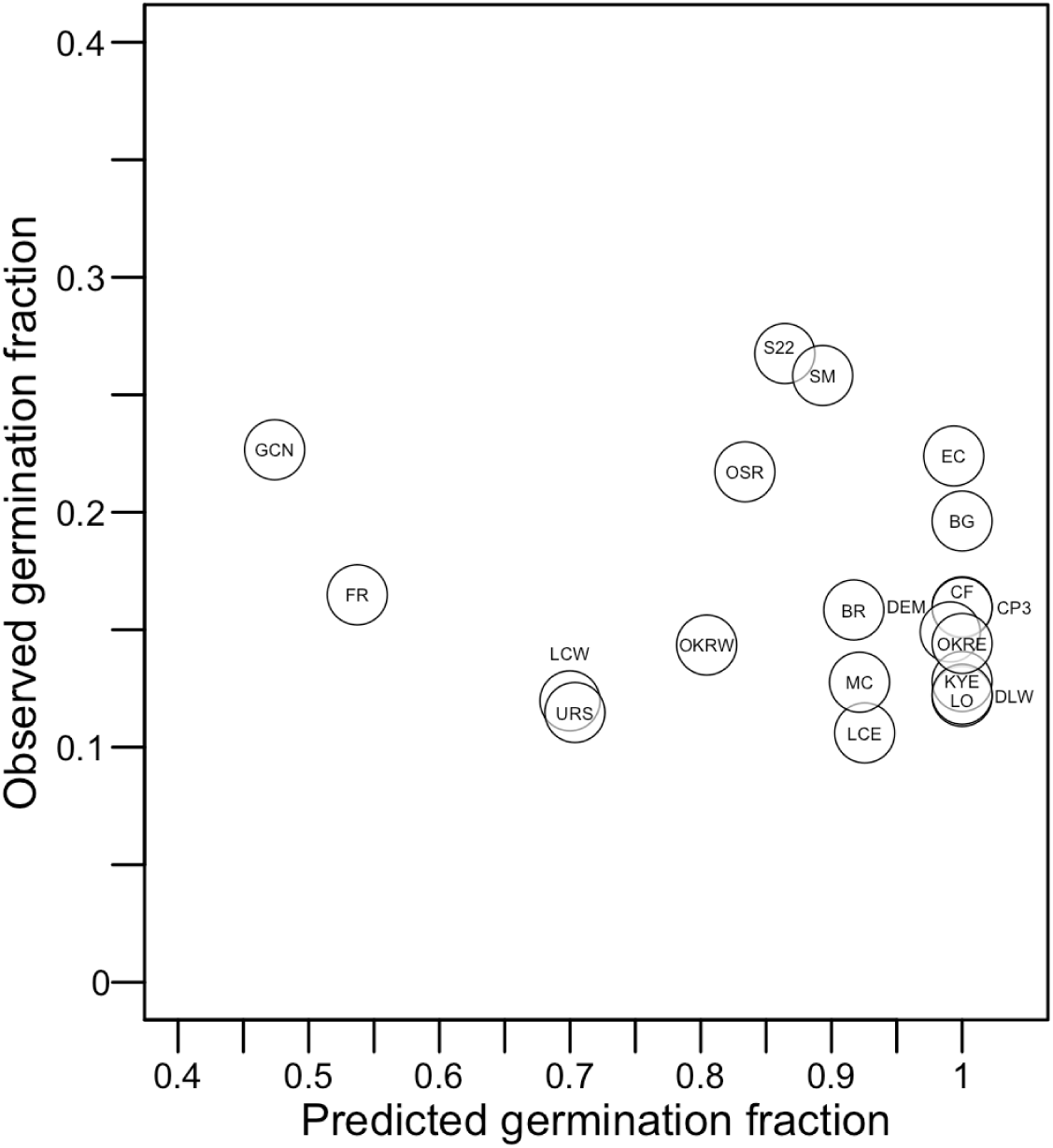
Comparison of observed and predicted, optimal germination fractions from a density-independent model of bet hedging. Each point is the population-specific mode of the posterior of *g*_1_ plotted against the predicted, optimal germination fractions. The observed germination fractions, *g*_1_, are estimated from the model for seed bank vital rates fit to data from the 2005-2008 seed bag experiments. For each population, we found the predicted, optimal germination fraction for a density-independent population model in which we resampled the annual estimates of per capita reproductive success.

### Germination and seed survival are uncorrelated

To assess the relationship between germination and risk experienced by seeds that remain in the seed bank, we calculated the correlation between germination fraction and seed survival. We did not observe a correlation between germination and seed survival in the seed bank (Fig. 5A). The 95% credible interval for the posterior distribution of the correlation between germination and seed survival overlapped zero.

**Figure 5:**
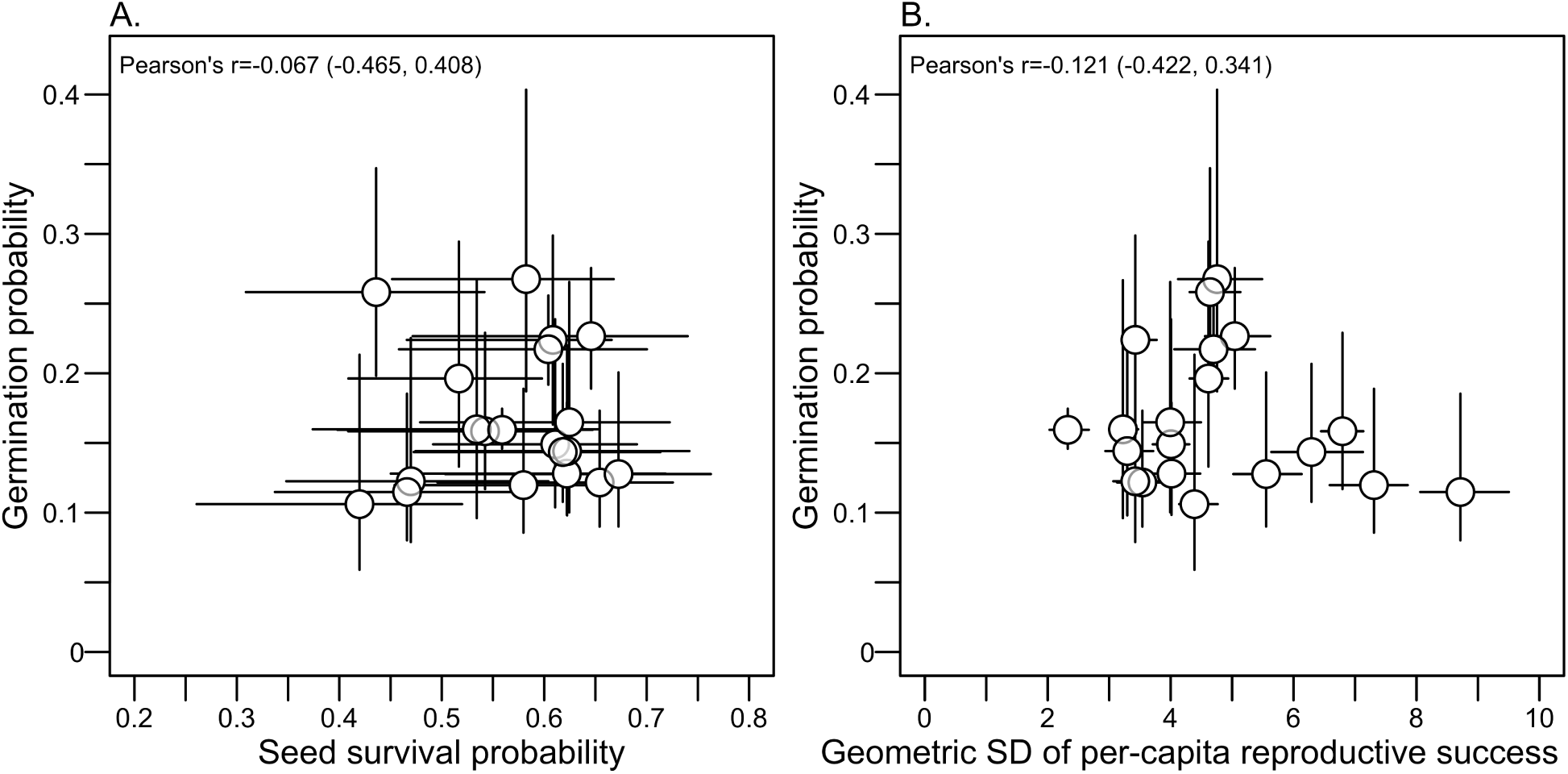
Relationship between germination and seed survival, and between germination and the geometric standard deviation of per capita reproductive success. (A) The observed germination probability, *g*_1_, plotted against the probability of seed survival, *s*_2_*s*_3_. (B) Correlation between observed germination probability, *g*_1_, and the geometric standard deviation of per capita reproductive success, a measure of the temporal temporal variability in per capita reproductive success. In both panels, points are the posterior modes; error bars are the 68% highest posterior density intervals (under a normal distribution, 68% of the distribution is within *±*1 standard deviation). The inset text gives the correlation and associated 95% credible interval.

### Germination and variability in per capita reproductive success are uncorrelated

To assess the relationship between germination and risk experienced after germination, we calculated the correlation between germination fraction and geometric standard deviation in per capita reproductive success. The correlation between germination and geometric standard deviation in per capita reproductive success was negative (Fig. 5B). However, the 95% credible interval for the posterior distribution of the correlation overlapped zero, indicating that there was not strong support for a non-zero correlation between germination and variability in reproductive success.

## Discussion

We used an extensive demographic dataset to conduct an unusually comprehensive test of whether bet hedging explained germination patterns among populations of *Clarkia xantiana* ssp. *xantiana*. All 20 populations in our study exhibited delayed germination. However, we found weak support for the expected trade-off between arithmetic and geometric mean fitness, mixed support that delayed germination acted as bet hedging, and no evidence that observed germination fractions were adaptive under a density-independent bet hedging model. Observed germination fractions were also uncorrelated with risk experienced by seeds that remain in the seed bank, or by plants after germination. Collectively, we interpret our results to suggest that delayed germination acting as bet hedging alone is insufficient to explain germination patterns among our study populations.

### Demographic test of bet hedging

To determine if delayed germination functions as bet hedging in each population, we tested for a trade-off between arithmetic and geometric mean population growth rate mediated by reduced variability in population growth rate (Cohen 1966; Philippi and Seger 1989). We observed average germination fractions below 0.3 in all populations. However, our demographic analysis failed to demonstrate the expected trade-off between mean and stochastic population growth rate for most populations, despite 15 years of observations of reproductive success (Table 3). We evaluated a strategy with the observed germination fraction against a strategy with no seed bank (Evans et al. 2007). Delayed germination reduced average population growth rate (Fig. 3A) and variance in population growth rates (Fig. 3B). But for most populations, delayed germination did not increase long-term stochastic population growth rate (Fig. 3C). Assuming quasi-complete germination, rather than complete germination, did not qualitatively change our results (Fig. S19).

**Table 3:** Summary of key results for tests of bet hedging.

|  |  |
| --- | --- |
| DEMOGRAPHIC TEST OF BET HEDGING | Summary |
| Reproductive success accounting for complete reproductive failure |  |
| 1. $\lambda_a(\text{noSB}) > \lambda_a(\text{SB})$ | 20/20 populations |
| 2. $\text{Var}(\lambda(\text{noSB})) > \text{Var}(\lambda(\text{SB}))$ | 20/20 populations |
| 3. $\lambda_s(\text{noSB}) < \lambda_s(\text{SB})$ | 7/20 populations |
| PREDICTED VS. OBSERVED GERMINATION | Summary |
| Germination fractions less than 1 | 13/20 populations |
| LIFE HISTORY COMPONENTS | Posterior mode (95% credible interval) |
| Correlation between germination and seed survival | $\rho_{g,s_2s_3} = -0.067 (-0.465, 0.408)$ |
| Correlation between germination and geometric standard deviation of per capita reproductive success | $\rho_{g,\text{GSD}} = -0.121 (-0.422, 0.341)$ |

### Observed germination fractions are lower than predicted by bet hedging models

To complement the demographic test of bet hedging, we calculated the optimal germination fractions that maximize each population’s growth rate (Childs et al. 2010; Simons 2011). We derived these optimal germination fractions by parameterizing density-independent population models with estimates of seed survival and reproductive success (Gremer and Venable 2014). The optimal germination fractions that we calculated suggest that the observed levels of seed mortality and temporal variability in reproductive success favored delayed germination in 13 populations (Fig. 4; Table 3). In comparison to the demographic test of bet hedging (Fig. 3), the optimal germination fractions (Fig. 4) thus provided slightly more support for the idea that delayed germination acts as bet hedging.

However, our predicted optimal germination fractions were much higher than the germination fractions observed in the field. We may have underestimated germination if we missed seedlings that died before, or if there was additional germination after, our annual census of seed bags. But the predicted germination fractions were 2 to 5 times the observed fractions, and we think it is unlikely that we underestimated germination to this extent. We also did not find the expected positive correlation between observed and predicted germination fractions. Jointly, we interpret these results to suggest that even when delayed germination is favored, the observed germination fractions are lower than would be adaptive under density-independent bet hedging alone. Our results parallel the findings in Gremer and Venable (2014) that density-independent models tend to predict higher germination fractions than observed in the field.

### Germination and risk across the life cycle

Under bet hedging, we expected that seeds from populations that experienced a greater degree of risk in the seed bank would have lower germination fractions (Cohen 1966; Gremer and Venable 2014; Venable 2007). High mortality risk in the soil seed bank should select against delayed germination, but we did not find support for the expected relationship between germination and seed survival (Fig. 5A; Table 3). Some populations with low seed survival exhibited low germination (e.g., FR, BR, SM), while some populations with high seed survival exhibited high germination (e.g., S22, CP3).

High variability in per capita reproductive success should also select for delayed germination (Cohen 1966). However, in our study populations, variability in reproductive success was uncorrelated with germination (Fig. 5B; Table 3). We observed similar germination fractions (approximately 0.1) for populations with very different levels of variability in reproductive success (similar germination probabilities for a range of geometric standard deviations from 3-9 in Fig. 5B).

Observed germination fractions varied more than twofold among populations but spanned a relatively small range (approximately 0.1-0.3; compared to the range of roughly 0.1-0.8 for the interspecific comparison in Venable 2007). The limited range of germination fractions may have made it more challenging to find support for the predicted correlations between germination and components of the life cycle. However, all populations had low germination fractions and this should have made it more likely to observe a trade-off between arithmetic and geometric mean fitness, and to find a greater adaptive value for delayed germination. Absence of evidence for negative correlations between germination and seed survival, and between germination and variability in reproductive success, is thus consistent with other results in our study. Populations with low variability in reproductive success and low germination were often the same populations that did not experience complete reproductive failure (Fig. 1B), for which stochastic population growth rates were higher without a seed bank, and for which we predicted high optimal germination fractions (e.g., OKRE, CP3 in Fig. 4).

### Temporal variability in reproductive success

Delayed germination decreases arithmetic mean fitness, and the variance in fitness, because it dampens the effect of years with low per capita reproductive success. To meet the criteria for bet hedging, delayed germination should also increase geometric mean fitness; whether it does so depends strongly on the minimum reproductive success or probability of reproductive failure (Childs et al. 2010; Cohen 1966; Evans et al. 2007). At the extreme, if there is no risk of reproductive failure, a strategy with delayed germination should always have lower geometric mean fitness than one with full germination. All populations in which stochastic population growth rate without a seed bank was lower than with a seed bank (URS, LCW, LCE, OKRW, FR, GCN, SM; Fig. 3C) either experienced reproductive failure or had no seedlings survive in permanent plots in at least one year (Fig. 1B). In contrast, populations in which stochastic population growth rate without a seed bank was higher than with a seed bank included those populations that either had some plants survive in permanent plots (LO) or populations in which plants set seed in all years. Although our demographic observations were exceptionally broad, 15 years of observations may have been insufficient to encounter reproductive failure in some populations. It is also possible that our observations and experiments underestimate the magnitude of temporal variability in per capita reproductive success of germinants. For example, our seed bag experiments capture seed germination but provide little information about events very early in a seedling’s life between germination and emergence (Chambers and MacMahon 1994). Early seedling mortality that exhibits substantial temporal variability would favor delayed germination. Our measurements may thus be conservative for testing predictions of bet hedging theory. At the same time, California is experiencing an ongoing drought and the 2005-2020 study period included precipitation anomalies with severe ecological impacts (Cook et al. 2015; Prugh et al. 2018; Williams et al. 2022). Studies of bet hedging through delayed germination often assume precipitation variability is a primary driver of variability in fitness (e.g., Clauss and Venable 2000; Philippi 1993*b*; Tielbö rger et al. 2012; Venable 2007). If precipitation variability were the dominant determinant of reproductive success in *C. x.* ssp. *xantiana*, we expect that our study would have had a high chance of observing its effects. We conducted an exploratory analysis to examine to what extent precipitation variability drives variability in reproductive success in our study populations (Appendix S6.5). We focused on the relationship between growing season precipitation and fitness, and found that the sensitivity of per capita reproductive success to growing season precipitation differed among populations (Fig. S21). Our analysis suggests that differences in precipitation variability alone are insufficient to explain differences in variability in per capita reproductive success among our study populations. Instead of aiming to identify specific environmental drivers of demographic variability, a complementary approach might be to model vital rates as a function of an unidentified, latent variable that describes environmental variability (Hindle et al. 2018). By adopting a model that describes the temporal covariance of vital rates, it would be possible to explore how population dynamics and optimal germination fractions respond to a range of variability beyond what we observed during the study period.

### Seed mortality across the life cycle

High seed mortality in the seed bank selects against delayed germination (Brown and Venable 1986; Cohen 1966; Donohue et al. 2010), but seed mortality between seed set and the opportunity to germinate or enter the seed bank can favor the evolution of delayed germination (Brown and Venable 1991). Seed mortality after the germination opportunity is a risk borne by seeds that remain in the seed bank. In contrast, seed mortality after seed set but before those new seeds germinate or enter the seed bank can reduce reproductive success (Brown and Venable 1991). It may thus be safer for a seed to remain in the seed bank if there is substantial seed mortality between seed set and the opportunity to germinate.

We conducted a follow-up analysis that shows the optimal germination fractions we predicted are more sensitive to estimates of seed survival before than after germination (Appendix S6.6). Optimal germination fractions could thus be lower than we predicted if we overestimated seed survival before germination (*s*_0_ or *s*_1_). To estimate survival from seed set in June/July to burial in October, *s*_0_, we combined observations from surveys and field experiments. We may have overestimated survival if our approach failed to fully capture mortality due to seed predation. In addition, the seed bag burial experiments could have overestimated seed survival from October to January, *s*_1_, if deep burial of seeds is a major source of loss from the seed bank, as bags prevent seeds from mixing into the soil. However, the experiments may underestimate survival if seed densities in bags are high enough to promote the growth of pathogenic fungi (Van Mourik et al. 2005). These caveats could also affect estimates of seed survival after germination (*s*_2_ or *s*_3_), but the optimal germination fraction is not as sensitive to these parameters.

### Intra- and interspecific interactions shape optimal germination fractions

In this study, we used a density-independent model of bet hedging, which is particularly sensitive to variability in reproductive success resulting from complete reproductive failure (Cohen 1966). Our estimates of per capita reproductive success implicitly incorporate the effects of density (Ellner 1985*b*), but we did not explicitly model density-dependence. Density-dependence can affect the value of delaying germination because competitors may alter reproductive success; years that would otherwise be good for growth and reproduction may become less favorable if there is strong competition (Ellner 1985*a*,*b*). In a density-dependent model, germination fractions are expected to be sensitive to variability in reproductive success rather than to just the probability of reproductive failure (Ellner 1985*a*). In the density-dependent case, optimal germination fractions are also expected to show a strong relationship with seed mortality rates, especially when mortality is low (Ellner 1985*a*). Optimal germination fractions may thus be lower than we predicted in this study if we were to calculate evolutionary stable strategies that account for competition (e.g., Gremer and Venable 2014).

To evaluate support for the hypothesis that density-dependence could play a role in our populations, we conducted exploratory analyses that examine whether transitions from seed to seedling (Appendix S6.7) and from seedling to fruiting plant (Appendix S6.8) exhibited density-dependence. The results of our exploratory analyses suggest that all, or the majority, of populations are subject to density-dependence in both transitions. Density-dependence could also act on other components of the life cycle; for example, competition from neighbors might also affect plant growth and fruit production. The mismatch between observed and predicted germination fractions that we report here might thus be partially explained by density-dependence in per capita reproductive success.

More broadly, competitive and facilitative interactions with intra- and inter-specific plant neighbors, as well as with pollinators, herbivores, and seed predators, could all modify the temporal variability of reproductive success. Reproductive success in *C. x.* ssp. *xantiana* is affected by insect pollinators (Moeller 2004), mammalian herbivores (Benning et al. 2019), and plant neighbors (James and Geber 2022). If these interactions amplify variability in per capita reproductive success, they could also favor lower germination fractions than those we predicted here (Brown and Venable 1991). Crucially, we would need to measure and model the temporal variability in the effect of these interactions in order to understand their impact on the evolution of delayed germination.

### Phenotypic plasticity in germination

To test bet hedging theory, we estimated fixed, population-level germination fractions with field experiments in which we collected and buried seeds in the same population. While we assumed that the germination fractions reflected genetic differentiation among populations, germination phenotypes can be influenced by seed genotype, maternal genotype, and offspring or maternal environment (Clauss and Venable 2000; Lampei et al. 2017; Philippi 1993*a*; Tielbö rger and Petrů 2010; Tielbörger et al. 2012). We could not partition the relative contribution of these influences in this study but, in general, germination phenotypes of *C. x.* ssp. *xantiana* do exhibit plasticity. In the field, germination varies interannually with rainfall (Geber and Eckhart 2005) and among microsites (James et al. 2020). In the lab, germination responds to water potential and temperature (I. Vergara and V. M. Eckhart, *unpublished data*). If germination reflects a response to environmental cues such as these, the distribution of those cues in the study years would determine the observed germination fractions (Clauss and Venable 2000). Studies that experimentally partition phenotypic variation in germination phenotypes of *C. x.* ssp. *xantiana* would be extremely valuable in complementing the present work.

Our results suggest that variation in germination fractions among populations of *C. x.* ssp. *xantiana* is unlikely to be explained exclusively by bet hedging. We hypothesize that germination strategies in these populations are likely shaped by the combined influence of bet hedging and plasticity. Bet hedging assumes that reproductive success is unpredictable at the time of germination (Cohen 1966). If germination responds to environmental cues that also predict reproductive success, plasticity should evolve in accordance to the correlation between the cue and fitness; such adaptive germination plasticity is termed predictive germination (Cohen 1967; Venable and Lawlor 1980). Bet hedging and predictive germination can operate both among or within years, providing seeds with multiple avenues to reduce the risk associated with germination (Donohue et al. 2010; Gremer et al. 2016). More generally, strategies are expected to combine bet hedging and plasticity in proportion to the uncertainty and predictability in the environment (Donaldson-Matasci et al. 2013; Tufto 2015). Predictive germination and density-dependence in per capita reproductive success may also interact to alter selection for delayed germination (Kortessis and Chesson 2019).

Empirical studies with other species suggest that germination strategies may often be a mix of bet hedging and predictive germination (Clauss and Venable 2000; Evans et al. 2007; Gremer et al. 2016; Simons 2014). To incorporate predictive germination into our bet hedging model, we could build on the approach taken by Gremer et al. (2016). Briefly, we would estimate annual germination fractions and retain the observed correlation between germination and reproductive success when calculating population growth rates. Estimating the correlation between germination and reproductive success would require more data than we have with the three years of seed bag burial experiments. Examining how bet hedging and plasticity jointly contribute to the evolution of delayed germination in *C. x.* ssp. *xantiana* would be an excellent task for future work.

## Acknowledgments

We thank A. A. Agrawal, S. P. Ellner, T. E. X. Miller, and W. F. Morris for conversations and feedback on the study. We appreciate feedback from K. E. Eisen, A. R. M. James, and R. Petipas during project development. We thank Erol Akçay, Rachel B. Spigler, Pedro F. Quintana-Ascencio, and an anonymous reviewer for helpful comments on the manuscript. We thank the field assistants, and undergraduate students and graduate students from Cornell University, Grinnell College, and the University of Minnesota, who contributed to the field surveys and experiments described in this study. G. Siegmund was supported by a Graduate Research Fellowship (DGE-1144153) from the U.S. National Science Foundation, and a Presidential Life Science Fellowship and Cornell Fellowship from Cornell University. The research was funded by U.S. National Science Foundation grants DEB-0515428, DEB-1256288 and DEB-1754299 to M. A. Geber, DEB-0515409, DEB-1256316, and DEB 1754157 to V. M. Eckhart, and DEB-0515466, DEB-1255141 and DEB-1754026 to D. A. Moeller.

## Author contributions

GS and MAG conceived of the ideas and analysis, using data collected by MAG, VME, and DAM. GS analyzed the data with input from MAG; GS wrote the manuscript with input from MAG. All authors contributed critically to the drafts.

