## Supplement A for "Bet hedging is not sufficient to explain germination patterns of a winter annual plant"

David A. Moeller[2,]

Vincent M. Eckhart[3,]

Monica A. Geber[1]

1. Department of Ecology and Evolutionary Biology, Cornell University, Ithaca, New York 14853;

2. Department of Plant and Microbial Biology, University of Minnesota, St. Paul, Minnesota 55108;

3. Department of Biology, Grinnell College, Grinnell, Iowa 50112

**S3 Priors** **28**

**S4 Model convergence** **30**

**S5 Computing vital rates** **34**

**S7 Glossary** **70**

### S1 Data summary

Table S1: Population names and geographic position.

| Population | Population name | Easting (km) | Northing (km) |
|---|---|---|---|
| LO | Live Oak | 342.78 | 3927.31 |
| URS | Upper Richbar South | 344.14 | 3926.22 |
| LCW | Lucas Creek West | 344.95 | 3927.66 |
| LCE | Lucas Creek East | 345.22 | 3927.83 |
| CF | Cow Flat | 3509 | 3932.83 |
| DEM | Democrat | 352.53 | 3932.92 |
| DLW | Delonegha West | 353.13 | 3934.38 |
| MC | Mill Creek | 353.37 | 3933.46 |
| OKRW | Old Kern Road West | 356.87 | 3936.26 |
| OKRE | Old Kern Road East | 3587 | 3936.69 |
| FR | Freeway Ridge | 359.38 | 3938.57 |
| BG | Black Gulch | 361.56 | 3939.68 |
| BR | Borel Road West | 362.54 | 3939.35 |
| KYE | Keyesville East | 362.86 | 39439 |
| OSR | Old State Road | 365.78 | 3951.48 |
| EC | Erskine Creek | 3696 | 3939.42 |
| S22 | Site 22 | 3698 | 3966.92 |
| CP3 | Camp 3 | 369.11 | 3962.98 |
| GCN | Golf Course North | 370.58 | 3954.57 |
| SM | Squirrel Mountain | 371.68 | 3940.78 |

Table S2: Sample sizes of dataset from seed bag burial experiment.

| | Age 0 | | | Age 1 | | Age 2 |
|---|---|---|---|---|---|---|
| Population | 2006 | 2007 | 2008 | 2006 | 2007 | 2008 |
| LO | 10 | 9 | 10 | 10 | 11 | 9 |
| URS | 10 | 9 | 10 | 5 | 10 | 4 |
| LCW | 10 | 10 | 10 | 9 | 10 | 9 |
| LCE | 10 | 10 | 9 | 9 | 7 | 9 |
| CF | 10 | 10 | 10 | 10 | 10 | 10 |
| DEM | 9 | 10 | 10 | 7 | 7 | 6 |
| DLW | 10 | 9 | 9 | 8 | 10 | 6 |
| MC | 9 | 10 | 10 | 8 | 9 | 9 |
| OKRW | 10 | 10 | 10 | 10 | 9 | 8 |
| OKRE | 10 | 11 | 10 | 10 | 9 | 9 |
| FR | 10 | 8 | 10 | 8 | 10 | 5 |
| BG | 10 | 10 | 10 | 7 | 10 | 5 |
| BR | 10 | 10 | 10 | 9 | 10 | 10 |
| KYE | 10 | 10 | 10 | 10 | 10 | 9 |
| OSR | 10 | 10 | 10 | 8 | 10 | 9 |
| EC | 11 | 10 | 10 | 8 | 10 | 8 |
| S22 | 10 | 10 | 10 | 8 | 10 | 8 |
| CP3 | 10 | 10 | 10 | 9 | 6 | 7 |
| GCN | 10 | 10 | 10 | 9 | 9 | 7 |
| SM | 10 | 10 | 9 | 8 | 10 | 10 |

Note: The table shows the number of seed bags in which seeds and/or seedlings were counted in each population, for each experimental round of the seed bag burial experiment. Seed bags are tallied in this table if they were recovered in both January and October of an experimental round. The number of seed bags varies because bags were lost or damaged in the field.

Table S3: Sample sizes of dataset on viability of seeds from seed bag burial experiment.

| | Age 0 | | | Age 1 | | Age 2 |
|---|---|---|---|---|---|---|
| Population | 2006 | 2007 | 2008 | 2006 | 2007 | 2008 |
| LO | 10 | 9 | 10 | 10 | 11 | 9 |
| URS | 7 | 9 | 9 | 5 | 9 | 4 |
| LCW | 10 | 10 | 5 | 9 | 7 | 8 |
| LCE | 10 | 10 | 9 | 9 | 7 | 9 |
| CF | 10 | 10 | 10 | 10 | 10 | 10 |
| DEM | 8 | 9 | 10 | 7 | 7 | 6 |
| DLW | 9 | 9 | 9 | 8 | 9 | 6 |
| MC | 9 | 10 | 10 | 8 | 9 | 9 |
| OKRW | 10 | 10 | 8 | 9 | 9 | 8 |
| OKRE | 10 | 11 | 10 | 10 | 7 | 9 |
| FR | 9 | 8 | 10 | 8 | 10 | 4 |
| BG | 7 | 10 | 10 | 6 | 10 | 3 |
| BR | 10 | 10 | 10 | 9 | 10 | 9 |
| KYE | 10 | 10 | 10 | 9 | 9 | 9 |
| OSR | 10 | 10 | 10 | 8 | 9 | 9 |
| EC | 9 | 10 | 10 | 8 | 10 | 8 |
| S22 | 9 | 10 | 10 | 8 | 10 | 8 |
| CP3 | 7 | 10 | 9 | 9 | 6 | 7 |
| GCN | 10 | 10 | 10 | 9 | 9 | 7 |
| SM | 9 | 10 | 9 | 8 | 10 | 10 |

Note: The table shows the number of seed bags from which seeds were used in lab viability assays in each population, for each experimental round. Seed bags are tallied in this table if they were recovered in October of an experimental round. The number of seed bags varies because bags were lost or damaged in the field.

Table S4: Sample sizes of dataset on seedling survival to fruiting.

| Population | 2006 | 2007 | 2008 | 2009 | 2010 | 2011 | 2012 | 2013 | 2014 | 2015 | 2016 | 2017 | 2018 | 2019 | 2020 |
|---|---|---|---|---|---|---|---|---|---|---|---|---|---|---|---|
| LO | 12 | 15 | 28 | 29 | 27 | 2 | 1 | 19 | 5 | 11 | 6 | 19 | 10 | 25 | 30 |
| URS | 4 | 17 | 10 | 7 | 12 | 14 | 3 | 5 | 2 | 1 | – | 5 | – | 4 | 3 |
| LCW | 16 | 27 | 27 | 27 | 21 | 4 | – | 15 | – | 1 | – | 4 | 5 | 20 | 18 |
| LCE | 20 | 12 | 18 | 19 | 19 | 1 | 1 | 3 | 1 | 8 | 7 | 19 | 17 | 24 | 20 |
| CF | 20 | 21 | 28 | 29 | 29 | 21 | 23 | 27 | 15 | 15 | 5 | 22 | – | 19 | 26 |
| DEM | 18 | 17 | 14 | 21 | 24 | 25 | 18 | 22 | 3 | 9 | 4 | 21 | 18 | 28 | 26 |
| DLW | 16 | 18 | 14 | 15 | 17 | 22 | 16 | 19 | 1 | 13 | 5 | 11 | 4 | 19 | 22 |
| MC | 17 | 11 | 22 | 25 | 27 | 30 | 29 | 27 | 6 | 18 | 8 | 15 | – | 13 | 22 |
| OKRW | 19 | 19 | 22 | 20 | 19 | 12 | 9 | 13 | – | 3 | 1 | 3 | 1 | 12 | 8 |
| OKRE | 14 | 10 | 8 | 19 | 21 | 17 | 7 | 19 | 6 | 10 | 5 | 15 | 5 | 15 | 13 |
| FR | 20 | 28 | 27 | 27 | 30 | 30 | 24 | 25 | 7 | 15 | 3 | 17 | 8 | 28 | 30 |
| BG | 18 | 21 | 22 | 26 | 24 | 26 | 20 | 23 | 3 | 26 | 5 | 16 | 12 | 25 | 24 |
| BR | 19 | 30 | 29 | 30 | 30 | 30 | 29 | 30 | 9 | 27 | 5 | 26 | 25 | 29 | 30 |
| KYE | 18 | 28 | 28 | 30 | 30 | 30 | 27 | 28 | 1 | 27 | 9 | 12 | 5 | 10 | 10 |
| OSR | 15 | 13 | 9 | 9 | 23 | 26 | 18 | 20 | 1 | 14 | – | 1 | – | 5 | 21 |
| EC | 20 | 28 | 30 | 30 | 30 | 30 | 30 | 24 | 2 | 10 | 9 | 8 | 2 | 9 | 19 |
| S22 | 17 | 10 | 21 | 18 | 28 | 17 | 27 | 26 | – | 17 | 4 | 10 | 1 | 10 | 20 |
| CP3 | 18 | 19 | 19 | 13 | 19 | 8 | – | 10 | 1 | 7 | – | 6 | 1 | 11 | 15 |
| GCN | 18 | 20 | 15 | 20 | 28 | 29 | 22 | 27 | 5 | 17 | – | 1 | 5 | 19 | 21 |
| SM | 15 | 8 | 13 | 18 | 23 | 25 | 18 | 24 | – | 19 | 8 | 13 | – | 14 | 24 |

Note: The table shows the number of permanent plots, located along transects, in each population for which we recorded observations of seedlings and/or fruiting plants in each year.

Table S5: Summary of undercounting in the dataset on seedling survival to fruiting.

| Population | 2006 | 2007 | 2008 | 2009 | 2010 | 2011 | 2012 | 2013 | 2014 | 2015 | 2016 | 2017 | 2018 | 2019 | 2020 |
|---|---|---|---|---|---|---|---|---|---|---|---|---|---|---|---|
| LO | 0 | 33 | 7.1 | 10 | 0 | – | 0 | 0 | 0 | 9.1 | 33 | 11 | 50 | 20 | 0 |
| URS | 0 | 5.9 | 0 | 14 | 17 | 7.1 | 0 | 0 | 0 | 0 | – | 0 | – | 50 | 67 |
| LCW | 0 | 3.7 | 0 | 0 | 4.8 | 25 | – | 0 | – | 0 | – | 0 | 0 | 0 | 0 |
| LCE | 0 | 50 | 5.6 | 37 | 0 | 0 | 0 | 0 | 0 | 0 | 14 | 5.3 | 29 | 4.2 | 5 |
| CF | 0 | 9.5 | 7.1 | 3.4 | 17 | 9.5 | 0 | 0 | 6.7 | 0 | 0 | 4.5 | – | 16 | 0 |
| DEM | 0 | 35 | 14 | 0 | 29 | 4 | 0 | 0 | 0 | 0 | 0 | 0 | 22 | 7.1 | 0 |
| DLW | 0 | 11 | 14 | 13 | 29 | 4.5 | 6.2 | 0 | 0 | 0 | 40 | 0 | 50 | 5.3 | 9.1 |
| MC | 0 | 27 | 9.1 | 8 | 7.4 | 0 | 0 | 0 | 33 | 0 | 38 | 6.7 | – | 7.7 | 4.5 |
| OKRW | 0 | 5.3 | 0 | 5 | 37 | 33 | 0 | 0 | – | 0 | – | 0 | – | 8.3 | 0 |
| OKRE | 0 | 20 | 12 | 11 | 14 | 18 | 0 | 0 | 17 | 0 | 20 | 6.7 | 40 | 0 | 0 |
| FR | 0 | 3.6 | 7.4 | 3.7 | 3.3 | 0 | 0 | 0 | 43 | 0 | 33 | 0 | 0 | 11 | 0 |
| BG | 0 | 14 | 14 | 0 | 12 | 0 | 5 | 0 | 0 | 0 | 0 | 0 | 83 | 0 | 0 |
| BR | 0 | 3.3 | 10 | 0 | 33 | 0 | 3.4 | 0 | 44 | 0 | 20 | 0 | 76 | 17 | 0 |
| KYE | 0 | 3.6 | 29 | 0 | 43 | 3.3 | 0 | 3.6 | – | 3.7 | 0 | 0 | 20 | 30 | 10 |
| OSR | 0 | 7.7 | 11 | 0 | 39 | 15 | 0 | 0 | – | 0 | – | 0 | – | 40 | 0 |
| EC | 0 | 29 | 33 | 0 | 20 | 0 | 0 | 21 | 50 | 0 | 11 | 0 | 0 | 44 | 0 |
| S22 | 0 | 0 | 19 | 5.6 | 18 | 18 | 3.7 | 0 | – | 0 | 50 | 0 | 0 | 10 | 10 |
| CP3 | 0 | 5.3 | 21 | 15 | 0 | 12 | – | 0 | – | 0 | – | 0 | 0 | 0 | 13 |
| GCN | 0 | 0 | 27 | 0 | 32 | 17 | 0 | 0 | – | 0 | – | 0 | 40 | 11 | 0 |
| SM | 0 | 0 | 31 | 0 | 61 | 20 | 0 | 4.2 | – | 0 | 0 | 0 | – | 50 | 17 |

Note: Values are the percentage of permanent plots in which we observed fewer seedlings than fruiting plants in our field surveys.

Table S6: Sample sizes of dataset on total fruit equivalents per plant from all plots.

| Population | 2006 | 2007 | 2008 | 2009 | 2010 | 2011 | 2012 |
|---|---|---|---|---|---|---|---|
| LO | 98 | 114 | 253 | 472 | 199 | 40 | 3 |
| URS | 32 | 43 | 41 | 54 | 155 | 58 | 7 |
| LCW | 243 | 269 | 340 | 194 | 224 | 53 | 3 |
| LCE | 246 | 215 | 147 | 146 | 133 | 29 | 0 |
| CF | 282 | 165 | 225 | 186 | 357 | 238 | 97 |
| DEM | 177 | 111 | 56 | 203 | 387 | 128 | 93 |
| DLW | 208 | 130 | 117 | 150 | 187 | 110 | 73 |
| MC | 163 | 160 | 142 | 151 | 206 | 278 | 77 |
| OKRW | 280 | 60 | 71 | 81 | 208 | 101 | 6 |
| OKRE | 100 | 46 | 43 | 140 | 105 | 122 | 5 |
| FR | 261 | 188 | 154 | 176 | 427 | 151 | 17 |
| BG | 153 | 160 | 222 | 155 | 150 | 141 | 63 |
| BR | 349 | 230 | 744 | 237 | 491 | 205 | 125 |
| KYE | 285 | 211 | 324 | 211 | 403 | 225 | 75 |
| OSR | 277 | 301 | 266 | 186 | 314 | 274 | 149 |
| EC | 370 | 196 | 133 | 287 | 389 | 386 | 83 |
| S22 | 319 | 111 | 92 | 187 | 245 | 105 | 115 |
| CP3 | 279 | 227 | 146 | 200 | 200 | 107 | 25 |
| GCN | 240 | 169 | 156 | 125 | 267 | 258 | 153 |
| SM | 217 | 25 | 79 | 121 | 173 | 199 | 51 |

Note: The table shows the number of plants on which fruits were counted in permanent and haphazardly located plots.

Table S7: Sample sizes of dataset on undamaged and damaged fruits per plant from extra plots.

| Population | 2013 | 2014 | 2015 | 2016 | 2017 | 2018 | 2019 | 2020 |
|---|---|---|---|---|---|---|---|---|
| LO | 8 | 48 | 1 | 8 | 187 | 47 | 141 | 210 |
| URS | 0 | 0 | 0 | 79 | 38 | 0 | 12 | 8 |
| LCW | 0 | 0 | 0 | 0 | 49 | 155 | 115 | 98 |
| LCE | 25 | 53 | 60 | 149 | 118 | 160 | 151 | 198 |
| CF | 70 | 114 | 52 | 96 | 198 | 150 | 152 | 165 |
| DEM | 31 | 46 | 45 | 67 | 255 | 256 | 286 | 260 |
| DLW | 68 | 35 | 61 | 60 | 242 | 169 | 145 | 148 |
| MC | 5 | 77 | 44 | 46 | 132 | 113 | 57 | 88 |
| OKRW | 0 | 8 | 0 | 31 | 130 | 35 | 75 | 69 |
| OKRE | 68 | 31 | 32 | 38 | 97 | 36 | 114 | 43 |
| FR | 6 | 59 | 41 | 53 | 198 | 74 | 107 | 148 |
| BG | 41 | 92 | 52 | 56 | 102 | 164 | 169 | 194 |
| BR | 111 | 180 | 65 | 84 | 180 | 274 | 464 | 248 |
| KYE | 60 | 136 | 120 | 57 | 143 | 133 | 120 | 153 |
| OSR | 47 | 160 | 104 | 99 | 150 | 108 | 116 | 167 |
| EC | 54 | 42 | 96 | 66 | 151 | 6 | 172 | 229 |
| S22 | 1 | 29 | 69 | 102 | 259 | 18 | 197 | 242 |
| CP3 | 151 | 88 | 59 | 69 | 142 | 11 | 137 | 177 |
| GCN | 11 | 42 | 58 | 64 | 104 | 134 | 202 | 195 |
| SM | 61 | 3 | 28 | 0 | 52 | 18 | 138 | 297 |

Note: The table shows the number of plants on which undamaged & damaged fruits were counted, in permanent and haphazardly located plots.

Table S8: Sample sizes of dataset on seeds per undamaged fruit.

| Population | 2006 | 2007 | 2008 | 2009 | 2010 | 2011 | 2012 | 2013 | 2014 | 2015 | 2016 | 2017 | 2018 | 2019 | 2020 |
|---|---|---|---|---|---|---|---|---|---|---|---|---|---|---|---|
| LO | 32 | 44 | 30 | 30 | 37 | 2 | 2 | 24 | 30 | 0 | 30 | 28 | 28 | 25 | 30 |
| URS | 18 | 30 | 25 | 30 | 30 | 27 | 5 | 0 | 0 | 0 | 29 | 16 | 0 | 24 | 3 |
| LCW | 20 | 50 | 28 | 30 | 35 | 32 | 4 | 0 | 0 | 0 | 0 | 28 | 33 | 25 | 30 |
| LCE | 20 | 30 | 30 | 30 | 32 | 12 | 0 | 30 | 29 | 38 | 30 | 26 | 37 | 47 | 28 |
| CF | 20 | 45 | 30 | 29 | 34 | 30 | 27 | 28 | 30 | 26 | 28 | 31 | 33 | 31 | 28 |
| DEM | 20 | 32 | 29 | 30 | 32 | 27 | 27 | 30 | 24 | 28 | 30 | 25 | 29 | 18 | 29 |
| DLW | 20 | 29 | 22 | 30 | 31 | 28 | 25 | 33 | 1 | 30 | 29 | 32 | 35 | 39 | 30 |
| MC | 20 | 50 | 29 | 30 | 35 | 30 | 26 | 24 | 46 | 35 | 30 | 34 | 30 | 41 | 28 |
| OKRW | 20 | 28 | 33 | 30 | 34 | 28 | 4 | 0 | 9 | 0 | 27 | 26 | 29 | 34 | 29 |
| OKRE | 20 | 40 | 26 | 30 | 30 | 28 | 3 | 30 | 18 | 24 | 31 | 35 | 22 | 27 | 21 |
| FR | 20 | 34 | 31 | 30 | 31 | 31 | 10 | 2 | 46 | 30 | 38 | 31 | 31 | 32 | 33 |
| BG | 21 | 19 | 41 | 30 | 30 | 28 | 29 | 29 | 30 | 29 | 29 | 32 | 27 | 31 | 28 |
| BR | 20 | 29 | 32 | 30 | 29 | 18 | 29 | 39 | 31 | 31 | 30 | 27 | 32 | 42 | 27 |
| KYE | 20 | 30 | 30 | 30 | 30 | 30 | 28 | 25 | 30 | 29 | 27 | 31 | 30 | 27 | 42 |
| OSR | 20 | 32 | 32 | 30 | 30 | 28 | 29 | 29 | 30 | 37 | 32 | 33 | 30 | 31 | 31 |
| EC | 20 | 17 | 29 | 30 | 31 | 26 | 22 | 30 | 31 | 31 | 30 | 30 | 4 | 33 | 29 |
| S22 | 20 | 40 | 33 | 30 | 28 | 23 | 30 | 30 | 23 | 30 | 30 | 30 | 17 | 33 | 32 |
| CP3 | 20 | 36 | 41 | 30 | 30 | 29 | 21 | 30 | 30 | 21 | 29 | 29 | 11 | 42 | 27 |
| GCN | 20 | 29 | 29 | 30 | 32 | 30 | 29 | 27 | 28 | 29 | 30 | 30 | 30 | 38 | 32 |
| SM | 20 | 44 | 31 | 29 | 32 | 30 | 27 | 30 | 3 | 8 | 0 | 30 | 3 | 28 | 34 |

Note: The table shows the number of undamaged fruits that were collected to count seeds per population, per year.

Table S9: Sample sizes of dataset on seeds per damaged fruit.

| Population | 2013 | 2014 | 2015 | 2016 | 2017 | 2018 | 2019 | 2020 |
|---|---|---|---|---|---|---|---|---|
| LO | 4 | 14 | 0 | 27 | 29 | 4 | 19 | 25 |
| URS | 0 | 0 | 0 | 19 | 20 | 0 | 5 | 9 |
| LCW | 0 | 0 | 0 | 0 | 16 | 15 | 32 | 32 |
| LCE | 1 | 11 | 15 | 24 | 16 | 7 | 17 | 32 |
| CF | 22 | 29 | 27 | 29 | 28 | 28 | 20 | 26 |
| DEM | 5 | 14 | 25 | 30 | 20 | 28 | 12 | 30 |
| DLW | 8 | 0 | 30 | 30 | 30 | 33 | 17 | 25 |
| MC | 4 | 15 | 15 | 30 | 24 | 31 | 17 | 32 |
| OKRW | 0 | 4 | 0 | 21 | 24 | 5 | 23 | 23 |
| OKRE | 13 | 8 | 9 | 18 | 30 | 7 | 11 | 17 |
| FR | 2 | 25 | 15 | 32 | 26 | 17 | 5 | 27 |
| BG | 17 | 20 | 11 | 30 | 28 | 28 | 6 | 31 |
| BR | 24 | 25 | 23 | 30 | 26 | 26 | 8 | 30 |
| KYE | 23 | 34 | 15 | 28 | 32 | 31 | 36 | 45 |
| OSR | 1 | 19 | 26 | 36 | 20 | 25 | 17 | 30 |
| EC | 12 | 22 | 8 | 30 | 30 | 1 | 20 | 30 |
| S22 | 1 | 3 | 2 | 7 | 10 | 1 | 7 | 12 |
| CP3 | 23 | 11 | 9 | 14 | 20 | 4 | 37 | 18 |
| GCN | 1 | 0 | 3 | 7 | 22 | 30 | 24 | 24 |
| SM | 1 | 3 | 0 | 0 | 0 | 0 | 2 | 32 |

Note: The table shows the number of damaged fruits that were collected to count seeds per population, per year.

### S2 Statistical models

We use observational and experimental data from 20 populations of *Clarkia xantiana* ssp. *xantiana* to estimate and calculate transition probabilities across the life cycle. We first obtain parameter estimates for multilevel, statistical models that we fit to experiments with the seed bank and observations of seedlings, fruiting plants, fruits and seeds. We built separate statistical models for each population because we were interested in describing the life histories of individual populations. We subsequently calculated the components used to construct the population model. More specifically, we use the parameter estimates from the statistical model to calculate the components of the population model. We calculate population-specific vital rates for belowground parts of the life cycle, and year- and population-specific vital rates for aboveground parts of the life cycle. Here, we provide an overview of the relevant sections of the supplement that elaborate on these details of our modeling approach.

First, we describe the hierarchical structure of our models in terms of linear mixed models. All of our models share the same underlying structure, so this description is generic and not associated with any particular observations or experimental data. Second, we describe specific models for the observations we used in this study. For each of the specific models, we describe the observations and define the likelihood associated with the observations. We use this combination to construct specific hierarchical models. We illustrate each model with a directed acyclic graph (DAG). In each DAG, solid arrows depict the relationships among random variables and dashed arrows depict the deterministic relationships. We also write the posterior and joint distributions for each model, which fully define the sampling statement. Third, we describe the priors that we place on parameters in the models. In the main text, we discussed the general principles that we used to choose priors; here, we provide more detailed justification and references.

With these parameters in hand, we then calculate the vital rates that we use to construct the population model. We present the calculations that we use to translate the parameter estimates for the statistical models to the derived quantities (Hobbs and Hooten 2015, p. 194-6) that describe the life history of the *C. x.* ssp. *xantiana* populations in this study. By calling these derived quantities, we mean that they can not be estimated directly but are instead calculated from two or more other values. We emphasize the importance of this portion of the analysis by discussing the calculations in the main text, and we provide more details in the supplement.

#### *S2.1 Hierarchical model structure*

In this study, we estimate each population's average probability of seed survival and germination, and the average per capita reproductive success in each population in each year. We fit multilevel models to obtain population-specific estimates for belowground vital rates, and year- and population-specific estimates for aboveground vital rates.

We describe our approach in terms of linear mixed models before defining the specific models used in our analysis (Evans et al. 2010; Ogle and Barber 2020). For a single population, we generally had $i$ observations of a response in year $j$, which we write as $y_{ij}$. We assume that the mean of observations in year $j$, $\theta_j$, is drawn from a normal distribution with mean $\mu_0^{\text{pop}}$ and variance $(\sigma^{\text{pop}})^2$ (Fig. S1). The corresponding equation is

$$\theta_j = \mu_0^{\text{pop}} + \epsilon_{(j)}. \tag{S1}$$

The model includes a population-level intercept $\mu_0^{\text{pop}}$ and random effects $\epsilon_{(j)}$. The random effects can be written as $\epsilon_{(j)} \sim N(0, (\sigma^{\text{pop}})^2)$. For the moment, we focus on describing the hierarchical structure of the model but note that we use link functions for transformation to parameters that are appropriate for the likelihoods we use to model different observations (e.g., binomial for seed bag experiments; Poisson for counts of seed per fruit). The binomial and Poisson both have a single parameter, so we describe the hierarchical structure for this kind of structure. Such a linear mixed effects model with random intercepts for years is one method recommended for modeling inter-annual variation in demographic rates (Metcalf et al. 2015).

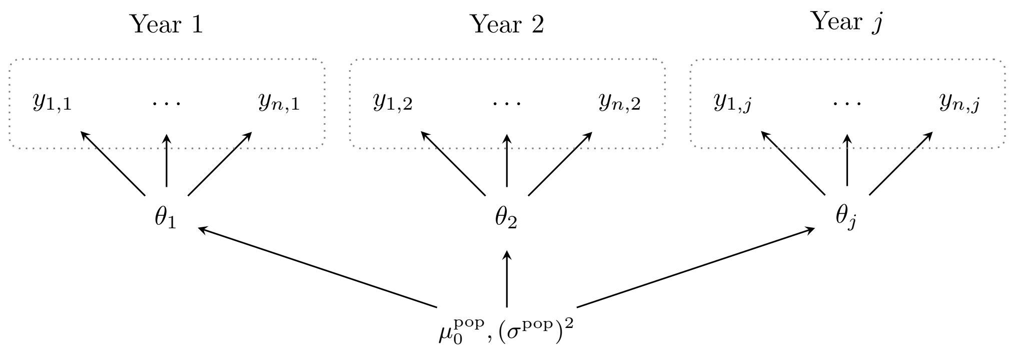

Figure S1: Graph depicting the general structure for the hierarchical models, for one population. Observations from each year, $y_{ij}$, are shown grouped and outlined by dotted lines. The observations are drawn from year-level parameters, $\theta_j$, which in turn are drawn from a distribution defined by population-level parameters, $\mu_0^{\text{pop}}$ and $\sigma^{\text{pop}}$.

We can rewrite the model with hierarchical centering (Ogle and Barber 2020) as

$$\theta_j \sim N(\mu_0^{\text{pop}}, (\sigma^{\text{pop}})^2). \tag{S2}$$

We are simply drawing year-level means from the population-level mean. For a single population, we write the posterior proportional to the joint distribution for this model as

$$[\theta_j, \mu_0^{\text{pop}}, (\sigma^{\text{pop}})^2 | y_{ij}] \propto [y_{ij}|\theta_j][\theta_j|\mu_0^{\text{pop}}, (\sigma^{\text{pop}})^2][\mu_0^{\text{pop}}][(\sigma^{\text{pop}})^2]. \tag{S3}$$

The distribution of the observations $y_{ij}$ is conditional on the year-specific parameter $\theta_j$. In turn, the year-specific parameter $\theta_j$ is conditional on the population-specific parameters $\mu_0^{\text{pop}}$ and $(\sigma^{\text{pop}})^2$. We place priors on all parameters found on the right hand side of conditional statements ($\mu_0^{\text{pop}}, (\sigma^{\text{pop}})^2$). In practice, we implement this model by specifying the population- and year-levels of the model with normal distributions; for example, $[\theta_j|\mu_0^{\text{pop}}, (\sigma^{\text{pop}})^2]$ is written as $\theta_j \sim N(\mu_0^{\text{pop}}, (\sigma^{\text{pop}})^2)$. The model thus describes a structure in which years are nested within populations.

#### S2.2 Posteriors and joint likelihoods

In the following sections, we use the general, hierarchical model structure described above to define statistical models for all the vital rates in this study. We describe models seedling survival to fruiting (Appendix S2.2.1), fruit and seed production (Appendix S2.2.2), seed vital rates (Appendix S2.2.3), and seed viability (Appendix S2.2.4). In each section, we define the model in three complementary ways: with a verbal description, with a directed acyclic graph, and with the posterior and joint distribution. The observations used in the models are defined in the text, and the parameters and their associated priors are given in Table S10.

##### S2.2.1 Seedling survival to fruiting

Seedlings can perish from a multitude of causes including end-of-season drought (Geber and Eckhart 2005), intra- and inter-specific competition (e.g., Geber and Eckhart 2005; James et al. 2020), small mammal herbivory (Benning et al. 2019), and fungal rust morality (Geber and Eckhart 2005). To describe variation in seedling survival, we wrote a model that has year-level means drawn from the population-level distribution for each study population (Fig. S2; Eq. S7). In the field, we counted seedlings ($n_{ijk}^{\text{seedlings}}$) and fruiting plants ($y_{ijk}^{\text{fruiting}}$) in plot $i$, year $j$, and population $k$. For each of 20 populations, indexed by $k$, we had observations for up to 15 years, indexed by $j$. Each year, we had $m_{jk}$ observations, which are indexed by $i$. In each year $j$ at population $k$, the average probability of seedling survival, $\mu_{S,jk}$, is drawn from a distribution defined by the population-level mean, $\mu_{S,k}^{\text{pop}}$, and variance, $(\sigma_{S,k}^{\text{pop}})^2$. The model has a binomial likelihood, and we used a logit link to model the probability of seedling survival. The full model is written as

$$\begin{aligned}
[\boldsymbol{\mu}_S, \boldsymbol{\mu}_S^{\textbf{pop}}, \boldsymbol{\sigma}_S^{\textbf{pop}} | \boldsymbol{y}^{\textbf{fruiting}}] \propto & \prod_{k=1}^{20}\prod_{j=1}^{J}\prod_{i=1}^{m_{jk}} \text{binomial}(y_{ijk}^{\text{fruiting}} | n_{ijk}^{\text{seedling}}, \text{logit}^{-1}(\mu_{S,jk})) \\
& \times \text{normal}(\mu_{S,jk} | \mu_{S,k}^{\text{pop}}, (\sigma_{S,k}^{\text{pop}})^2) \\
& \times \text{normal}(\mu_{S,k}^{\text{pop}} | 0, 1)\text{half-normal}(\sigma_{S,k}^{\text{pop}} | 0, 1).
\end{aligned} \tag{S4}$$

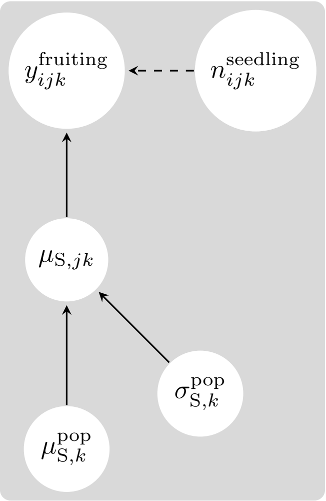

Figure S2: Directed acyclic graphs for the model for seedling survival to fruiting. All symbols are defined in the text.

##### *S2.2.2 Fruits per plant & seeds per fruit*

Seed production is the product of flowering, pollination, fruit production, and successful seed set. We used multiple types of observations to estimate seed production: total fruit equivalents per plant (2006-2012), total fruits per plant and proportion of fruits damaged by herbivores (2013-2020), seeds per undamaged fruit (2006-2020), and seeds per damaged fruit (2013-2020). We first describe the model for observations of total fruit equivalents, and then describe how we adapted this model for other observations.

In the field, we counted of total fruit equivalents ($y_{ijk}^{TFE}$) on plant $i$, in year $j$, and in population $k$. For each of 20 populations, indexed by $k$, we had observations for up to 7 years, indexed by $j$. At each population and in each year, we counted fruits on $m_{jk}$ plants, which are indexed by $i$. We modeled observations with a Poisson likelihood with a log-link. In each year $j$ at population $k$, the average number of total fruit equivalents per plant is $\mu_{\text{TFE},jk}$. Fruit counts are non-negative, so we assume that the annual mean total fruit equivalents per plant are drawn from a log-normal with parameters for the population-level mean, $\mu_{\text{TFE},k}^{\text{pop}}$ and variance on the log scale, $(\sigma_{\text{TFE},k}^{\text{pop}})^2$. To ensure that the population-level means are non-negative as required for the log-normal, we used a gamma distribution for the population-level means, $\mu_{\text{TFE},k}^{\text{pop}}$. Finally, we included observation-level random effects because counts of total fruit equivalents were overdispersed. We included an observation-level random effect, $\epsilon_{\text{TFE},ijk}$, defined separately for each population so that $\epsilon_{\text{TFE},ijk} \sim N(0, \sigma^2_{\epsilon,\text{TFE},k})$, where $\sigma^2_{\epsilon,\text{TFE},k}$ is the variance of the observation-level random effect. We represent the model total fruit equivalents per plant as a DAG (Fig. S3) and write the full model as

$$\begin{aligned}
&f(\boldsymbol{\mu}_{\textbf{TFE}}, \boldsymbol{\epsilon}_{\textbf{TFE}}) = \exp(\log(\mu_{\text{TFE},jk}) + \epsilon_{\text{TFE},ijk}) \\
&[\boldsymbol{\mu}_{\textbf{TFE}}, (\boldsymbol{\sigma}^{\textbf{pop}}_{\textbf{TFE}})^2, \boldsymbol{\nu}_{\textbf{TFE}} | \boldsymbol{y}^{\textbf{TFE}}] \propto \\
&\prod_{k=1}^{20}\prod_{j=1}^{J}\prod_{i=1}^{m_{jk}} \text{Poisson}(y^{\text{TFE}}_{ijk} | f(\mu_{\text{TFE},jk}, \epsilon_{\text{TFE},ijk})) \text{ lognormal}(\mu_{\text{TFE},jk} | \nu_{\text{TFE},k}, (\sigma^{\text{pop}}_{\text{TFE},k})^2) \\
&\times \text{normal}(\epsilon_{\text{TFE},ijk} | N(0, \sigma^2_{\epsilon,\text{TFE},k}) \\
&\times \text{gamma}(\nu_{\text{FE},k} | 1, 1) \text{ half-normal}((\sigma^{\text{pop}}_{\text{TFE},k})^2 | 0, 1) \text{half-normal}(\sigma^2_{\epsilon,\text{TFE},k} | 0, 1)
\end{aligned} \tag{S5}$$

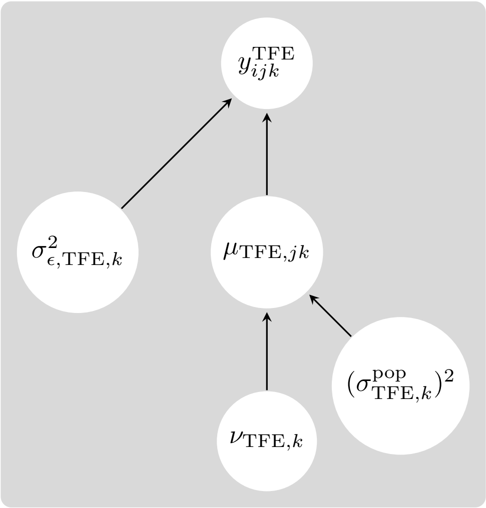

Figure S3: Directed acyclic graphs for the model for total fruit equivalents. All symbols are defined in the text.

From 2013-2020, we separately recorded the number of undamaged ($y^{UF}$) and damaged ($y^{DF}$) fruits on a plant. We summed these to compute the total number of fruits per plant ($y^{TOT}_{ijk}$) on plant $i$, in year $j$, and in population $k$. For each of 20 populations, indexed by $k$, we had observations for up to 8 years, indexed by $j$. At each population and in each year, we counted fruits on $m_{jk}$ plants, which are indexed by $i$. We modeled observations of total fruits the same way we modeled total fruit equivalents (see above). We write the model for total fruits below; the DAG has the same structure as in the model for total fruit equivalents so we do not repeat it here.

$$
\begin{aligned}
& f(\boldsymbol{\mu}_{\mathbf{TOT}}, \boldsymbol{\epsilon}_{\mathbf{TOT}}) = \exp(\log(\mu_{\text{TOT},jk}) + \epsilon_{\text{TOT},ijk}) \\
& [\boldsymbol{\mu}_{\mathbf{TOT}}, (\boldsymbol{\sigma}^{\mathbf{pop}}_{\mathbf{TOT}})^2, \boldsymbol{\nu}_{\mathbf{TOT}} | \boldsymbol{y}^{\mathbf{TOT}}] \propto \\
& \prod_{k=1}^{20} \prod_{j=1}^{J} \prod_{i=1}^{m_{jk}} \text{Poisson}(y^{\text{TOT}}_{ijk} | f(\mu_{\text{TOT},jk}, \epsilon_{\text{TOT},ijk})) \ \text{lognormal}(\mu_{\text{TOT},jk} | \nu_{\text{TOT},k}, (\sigma^{\text{pop}}_{\text{TOT},k})^2) \\
& \times \text{normal}(\epsilon_{\text{TOT},ijk} | N(0, \sigma^2_{\epsilon,\text{TOT},k}) \\
& \times \text{gamma}(\nu_{\text{TOT},k} | 1, 1) \ \text{half-normal}((\sigma^{\text{pop}}_{\text{TOT},k})^2 | 0, 1) \text{half-normal}(\sigma^2_{\epsilon,\text{TOT},k} | 0, 1)
\end{aligned}
\tag{S6}
$$

We then sought to estimate the proportion of fruits that were damaged by herbivores in the observations from 2013-2020. We modeled the number of damaged fruits on a plant ($y^{DF}$) as a sample from a binomial distribution with the number of trials equal to the total number of fruits on that plant ($y^{TOT}$). Damaged fruits were thus considered 'successes', and we modeled the proportion of fruits that were damaged per plant. For each of 20 populations, indexed by $k$, we had observations on total, $y^{TOT}_{ijk}$, and damaged, $y^{DF}_{ijk}$, fruits per plant for up to 8 years, indexed by $j$. At each population and in each year, we counted fruits on $m_{jk}$ plants, which are indexed by $i$. In each year $j$ at population $k$, the average probability of a fruit being damaged, $\mu_{\text{D},jk}$, is drawn from a distribution defined by the population-level mean, $\mu^{\text{pop}}_{\text{D},k}$, and variance, $(\sigma^{\text{pop}}_{\text{D},k})^2$. The model has a binomial likelihood, and we used a logit link to model the probability of fruit damage. The model has the same structure as the model for seedling survival to fruiting, so we do not repeat the DAG here. The model is written as

$$
\begin{aligned}
[\boldsymbol{\mu}_{\boldsymbol{D}}, \boldsymbol{\mu}^{\mathbf{pop}}_{\boldsymbol{D}}, \boldsymbol{\sigma}^{\mathbf{pop}}_{\boldsymbol{D}} | \boldsymbol{y}^{\mathbf{DF}}] \propto & \prod_{k=1}^{20} \prod_{j=1}^{J} \prod_{i=1}^{m_{jk}} \text{binomial}(y^{\text{DF}}_{ijk} | n^{\text{TOT}}_{ijk}, \text{logit}^{-1}(\mu_{D,jk})) \\
& \times \text{normal}(\mu_{D,jk} | \mu^{\text{pop}}_{D,j}, (\sigma^{\text{pop}}_{D,j})^2) \\
& \times \text{normal}(\mu^{\text{pop}}_{D,j} | 0, 1) \text{half-normal}(\sigma^{\text{pop}}_{D,j} | 0, 1).
\end{aligned}
\tag{S7}
$$

From 2006-2020, we counted the number of seeds per undamaged fruit ($y^{US}_{ijk}$) in fruit $i$, in year $j$, and in population $k$. For each of 20 populations, indexed by $k$, we had observations for up to 15 years, indexed by $j$. At each population and in each year, we counted seeds on $m_{jk}$ fruits, which are indexed by $i$. We modeled observations of seeds per undamaged fruit the same way we modeled total fruit equivalents and total fruits per plant (see above). In a given year, the mean seeds per undamaged fruit, $\mu^{\text{pop}}_{\text{US},jk}$ is drawn from a log-normal with parameters for the population-level mean, $\mu^{\text{pop}}_{\text{US},k}$ and variance on the log scale, $(\sigma^{\text{pop}}_{\text{US},k})^2$. We again included an observation-level term for overdispersed counts, with a variance of $\sigma^2_{\epsilon,\text{US},k}$.

Starting in 2013, we also collected damaged fruits ($y^{DS}_{ijk}$). At each population and in each year, we counted seeds on $n_{jk}$ fruits, which are indexed by $i$. To estimate the fraction of seeds lost to herbivory per fruit, we combined observations of seeds per undamaged and damaged fruits. We assumed that, on average, damaged fruits had a fraction of the seeds that undamaged fruits had. In each year $j$ at population $k$, the average proportion of seeds lost to herbivory, $\theta_{\text{DS},jk}$ is the inverse logit of $\mu_{\text{DS},jk}$ which is drawn from a distribution defined by the population-level mean, $\mu^{\text{pop}}_{\text{DS},k}$, and variance, $(\sigma^{\text{pop}}_{\text{DS},k})^2$. With the Poisson distribution as the likelihood, we model the mean seeds per damaged fruit as the product of proportion of seeds lost to herbivory and the estimated seeds per undamaged fruit. We combine the models for seeds per undamaged fruits and damaged fruits (Fig. S4), which we write as

$$
\begin{aligned}
&f(\boldsymbol{\mu}_{\mathbf{US}}, \boldsymbol{\epsilon}_{\mathbf{US}}) = \exp(\log(\mu_{\text{US},jk}) + \epsilon_{\text{US},ijk}) \\
&\text{logit}(\theta_{\text{DS},jk}) = \mu_{\text{DS},jk} \\
&[\boldsymbol{\mu}_{\mathbf{US}}, (\boldsymbol{\sigma}^{\mathbf{pop}}_{\mathbf{US}})^2, \boldsymbol{\nu}_{\mathbf{US}}, \boldsymbol{\mu}_{\mathbf{DS}}, (\boldsymbol{\sigma}^{\mathbf{pop}}_{\mathbf{DS}})^2, \boldsymbol{\mu}_{\mathbf{DS}} | \boldsymbol{y}^{\mathbf{US}}, \boldsymbol{y}^{\mathbf{DS}}] \propto \\
&\prod_{k=1}^{20}\prod_{j=1}^{J}\Big\{\prod_{i=1}^{m_{jk}}\text{Poisson}(y^{\text{US}}_{ijk}|f(\mu_{\text{US},jk}, \epsilon_{\text{US},ijk}))\ \text{lognormal}(\mu_{\text{US},jk}|\nu_{\text{US},k}, (\sigma^{\text{pop}}_{\text{US},k})^2) \\
&\times \text{normal}(\epsilon_{\text{US},ijk}|N(0, \sigma^2_{\epsilon,\text{US},k}) \\
&\times \text{gamma}(\nu_{\text{FE},k}|1,1)\ \text{half-normal}((\sigma^{\text{pop}}_{\text{US},k})^2|0,1)\text{half-normal}(\sigma^2_{\epsilon,\text{US},k}|0,1)\Big\} \\
&\times \Big\{\prod_{i=1}^{n_{jk}}\text{binomial}(y^{\text{DS}}_{ijk}|\theta_{\text{DS},jk} \times f(\mu_{\text{US},jk}, \epsilon_{\text{US},ijk}))\ \text{normal}(\mu_{DS,jk}|\mu^{\text{pop}}_{DS,k}, (\sigma^{\text{pop}}_{DS,k})^2) \\
&\times \text{normal}(\mu^{\text{pop}}_{S,j}|0,1)\text{half-normal}(\sigma^{\text{pop}}_{DS,k}|0,1)\Big\}.
\end{aligned}
\tag{S8}
$$

##### *S2.2.3 Seed persistence and emergence*

We combined observations of intact seeds and seedlings in seed bags with observations of seed rain and seedlings in permanent plots to study seed bank dynamics in the field. To infer seed persistence from fall (October) to winter (January/February), seedling emergence in the winter, and seed persistence from winter to fall, we used experiments in which we buried seeds in mesh

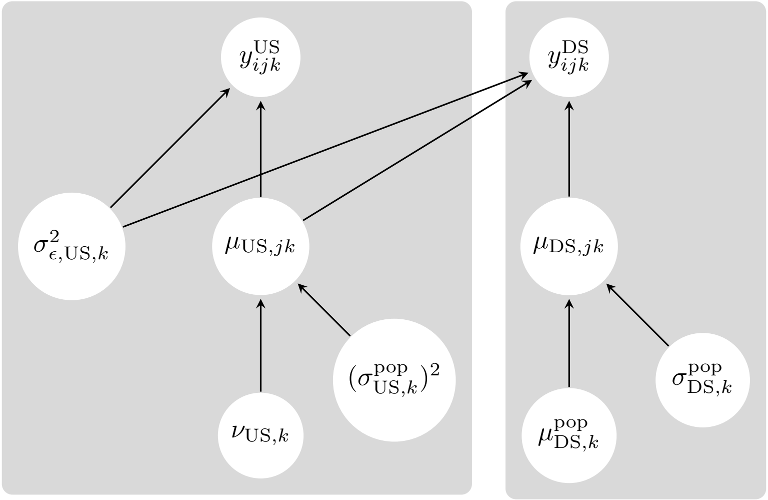

Figure S4: Directed acyclic graphs for the model for seeds per undamaged and damaged fruit. All symbols are defined in the text.

bags the field. To determine seed survival from seed production in July to four months later in October when seed bags were placed in the field, we combined observations from the seed bag burial experiment with observations of seed production and seedling emergence in permanent plots. In practice, we constructed a single statistical model to estimate seed bank dynamics; we discuss this model in detail in this section (Fig. S5; Eq. S11).

We use observations from the field experiment to estimate seed persistence and emergence, but these estimates do do not account for loss of viability. In the seed bag burial experiment, we counted intact seeds for up to three years, and counted seedlings after winter rains once per year. It is not possible to distinguish intact, and viable, seeds from intact, but not viable, seeds in the field. Seeds that experience physiological death may remain intact but may not be viable. To calculate seed survival and germination, which do account for loss of viability, we adjust persistence and emergence with estimates of viability obtained with the lab germination trial and viability assays (Appendix S2.2.4 and S5.2.3).

Seeds remain intact in the seed bank if they persist, i.e. remain intact, and do not emerge. Seeds leave the seed bank either by not persisting – experiencing mortality – or by emerging to become seedlings. In our statistical model for observations from the seed bag burial experiment, we linked the observations (intact seeds, seedlings) by describing seed loss from the seed bank as the product of two separate processes, mortality and emergence. We used methods from event history analysis (also known as survival analysis or failure time analysis) to describe seed loss from the seed bank by mortality and by emergence as discrete functions (Siegmund and Geber 2022). By taking the product of these functions, we define a single statistical model for observations of intact seeds and seedlings.

Our approach allowed us to use all of the data from the seed bag burial experiments simultaneously. In our experiment, we did not return the bags that we collected in October to the field because we tested the seeds in those bags for viability–sampling was thus destructive. This mean that we sampled a new set of bags in the second year of an experimental round, and a new set of bags in the third year. Due to variability among bags, seed counts could increase from one observation period to the next when we sampled a new set of bags. For example, the seed counts in the second year of a round could be higher than the seed counts at the end of the first year. The model that we constructed allowed us to reconcile such differences across observation periods because the model decided what amount of this was due to sampling variation and what was due to the processes of persistence and emergence.

To estimate emergence, the probability that an intact seed becomes a seedling, we model observations of seedlings from the seed bag burial experiment. We construct a model with a binomial likelihood and logit-link for the latent probability of emergence. The latent probability of emergence at each age is described by two hierarchical levels: the first level is the experimental years and the second (upper) level is the population.

To estimate seed persistence, the probability that a seed remains intact in the seed bank, we model observations of intact seeds from the seed bag burial experiment. For seeds to remain intact, they must not die and not emerge as seedlings. We thus model persistence, the probability of a seed remaining intact as the product of persistence and the complement of emergence (probability that seeds do not emerge). The latter describes the probability of not emerging as a seedling. In the language of survival analysis, the product of these processes defines a non-parametric 'survival function' (Klein and Moeschberger 2003). The non-parametric 'survival function' is simply the product of parameters that describe seeds persisting and not emerging at different points in time.

To model observations of intact, ungerminated seeds from the seed bag burial experiment, we thus combine parameters that describe seed persistence in the seed bank with parameters that describe seeds not emerging from the seed bank. We construct a model with a binomial likelihood and logit-link for the latent probability that a seed remains intact in the seed bank. The latent probability at each observation instance is described by the product of discrete components: the survival function and the discrete probabilities of germination, respectively. We assume that the parameters vary from year to year within each population. The latent probability of seed persistence at each age is described by two hierarchical levels: the first level is the experimental years and the second (upper) level is the population.

The seed bag burial experiment follows seeds from the start of the seed burial experiment in October after seed production. However, it does not provide direct information about what happens to seeds between seed production in July and October four months later. To estimate the survival of seeds from seed production to October, we augment the model described so far with one additional component. In addition to modeling seed persistence and seedling emergence in seed bags, we add a model for seedling emergence in permanent plots. We use estimates for the number of seeds produced in years $t-2$ and $t-1$ in permanent plots, seedlings emerging in permanent plots in year $t$, and our model for seed persistence and emergence to infer seed survival from seed production to October. We assume that the majority of seedlings in a plot emerge from seeds produced the previous two year and thus do not model emergence from older seeds.

To illustrate how this works, we write the number of seedlings that emerge in permanent plots in year $t$ as:

$$\text{seedlings}(t) = \text{seed rain}(t-1)\theta_0\theta_1\gamma + \text{seed rain}(t-2)\theta_0\theta_1(1-\gamma)\theta_2\theta_3\gamma \tag{S9}$$

In the above equation, $\theta$s are the probability of seed persistence for different time periods;

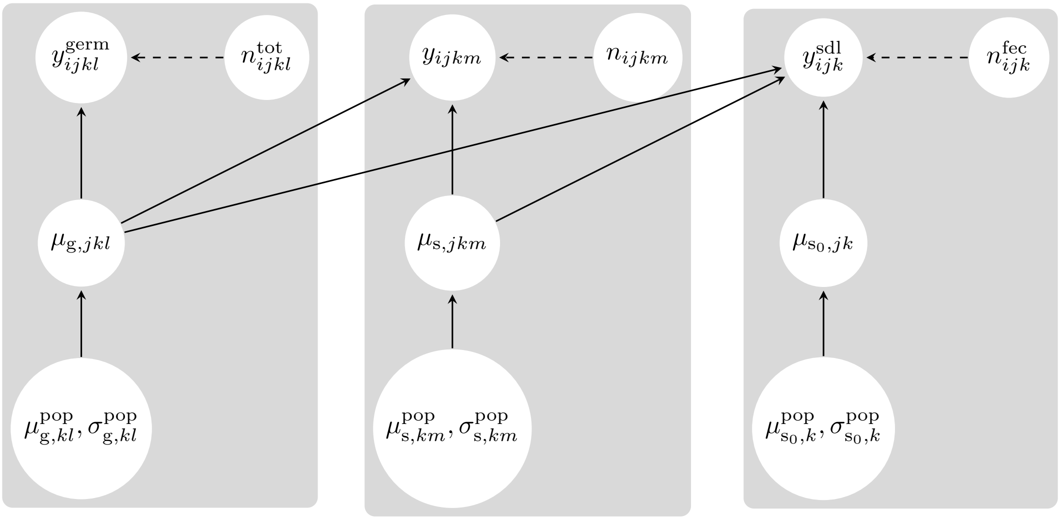

Figure S5: Directed acyclic graphs for the joint models for emergence, seed persistence, and seed survival from seed production to the October four months after seed production. From left to right, the directed acyclic graphs show models for emergence, seed persistence, and seed production. All symbols are defined in the text.

$\theta_0$: persistence from seed production in year $t$ to October in year $t$, $\theta_1$: persistence from October in year $t$ to January in year $t+1$, $\theta_2$: persistence from January in year $t+1$ to October in year $t+1$, $\theta_3$: persistence from October in year $t+1$ to January in year $t+2$. The $\gamma$ is the probability of emergence. By combining the estimates of seed persistence and emergence from the seed bag burial experiment with observations of seed rain and seedling emergence in permanent plots, we can infer seed persistence from seed production in July to October. Note that because we assume that all seeds are viable at the start of the seed bag burial experiment (Appendix S5.2.1), our estimate for seed persistence from July to October is simply seed survival from July to October. We do not further adjust this estimate for loss of viability.

We combine several datasets on aboveground plants to calculate seed rain. To calculate the number of fruits per plot, we multiply the average counts of fruits per plant from population-wide surveys with counts of plants per permanent plot. To calculate the number of seeds produced per plot, we multiply fruits per plot with the average number of seeds per fruit. We do this for aboveground data in 2006 and 2007. We link this data to the number of seedlings observed in those plots in 2008. We summed across plots in a transect and thus take the total number of seeds produced in a transect in year $t-1$ and $t-2$ as the number of trials in a binomial experiment for which the outcome is the number of seedlings observed in that transect in year $t$. The probability is the product of survival from seed production in July to October (estimated here), persistence from October to January, and emergence. We have estimates of the latter two probabilities from the seed bag experiments. We link the three components and estimate the probability of seed survival from seed production in July to October in 2006 and 2007.

Specifically, we observed seeds and seedlings in bags $i$, in year $j$, and at population $k$. We made observations in 3 years, indexed by $j$, at 20 populations, indexed by $k$. Each year, we observed $n_{jk}$ bags, which were indexed by $i$. For an experiment that started with $n$ bags in October, year $t$, observations of intact seeds, $y_{ijkm}$, were made at 6 times indexed by $m$ that describe seeds persisting up to...

- January $t+1$ ($m=1$)
- October $t+1$ ($m=2$)
- January $t+2$ ($m=3$)
- October $t+2$ ($m=4$)
- January $t+3$ ($m=5$)
- October $t+3$ ($m=6$)

We modeled the number of intact seeds, $y_{ijkm}$, at each time point as a sample from a binomial distribution with the number of seeds in the seed bag at the start of the experiment, $n_{ijkm}$, as the number of trials. The probability of remaining intact for up to $m$ intervals is described by a function $f(\dots)$ that accounts for both the probability of remaining intact and the probability of germinating. In each year $j$, population $k$, and interval $m$ the average probability of a seed persisting, $\mu_{\text{s},jkm}$, is drawn from a distribution defined by the population-level mean for a particular time interval, $\mu^{\text{pop}}_{\text{s},km}$, and variance, $(\sigma^{\text{pop}}_{\text{s},km})^2$.

We modeled the number of seedlings, $y_{\text{g},ijkl}$, as a sample from a binomial distribution with the total number of seeds before germination, $n_{\text{gijkl}}$, as the number of trials. In each year $j$, population $k$, and for each seed age $l$ the average probability of a seed emerging, $\mu_{\text{g},jkl}$, is drawn from a distribution defined by the population-level mean for a particular seed age, $\mu^{\text{pop}}_{\text{s},kl}$, and variance, $(\sigma^{\text{pop}}_{\text{s},kl})^2$.

Finally, we modeled the number of seedlings in permanent plots in 2008, $y_{\text{p},3k}$, as the sum of two sets of seedlings. We assume that the seedlings in 2008 are either from seeds produced in 2007, $y^{[t-2]}_{\text{p},3k}$, or 2006, $y^{[t-1]}_{\text{p},3k}$. We use the superscript to index the year in which seeds were produced, and keep the subscript consistent with the rest of the model: 3 refers to the third year of observations in the seed bag experiment and $k$ indexes populations. Each set of seedlings is a sample from a binomial distribution, with the seed rain in the appropriate year as the number of trials. In each year $j$ and population $k$ the average probability of a seed surviving from July to October, $\mu_{\text{s}_0,jk}$, is drawn from a distribution defined by the population-level mean, $\mu^{\text{pop}}_{\text{s}_0,k}$, and variance, $(\sigma^{\text{pop}}_{\text{s}_0,k})^2$.

For example, for seeds from 2007, we take the number of seeds produced in permanent plots in year 2 of the seed bag experiment (2007), $n_{\text{p},2k}$, as the number of trials. The average probability of a seed persisting from July to October, October to January, and then emerging as a seedling is the product of three terms. The first term is the probability of a seed surviving from July in year $t$ to October of year $t$; the second term is the probability of a seed persisting from October of year $t$ to October of year $t+1$; the third term is the probability of a seed that persisted to January $t+1$ emerging that January:

$$\left(\text{logit}^{-1}(\mu_{\text{s}_0,2k})\right) \times \left(f(\cdot, \mu_{\text{s},\cdot 2k,m=1})\right) \times \left(\text{logit}^{-1}(\mu_{\text{g},3k})\right) \tag{S10}$$

When we combine the model for observations of intact seeds in seed bags, seedling emergence in seed bags, and seedlings in permanent plots, the full model is written as

$$f(\mu_{\mathrm{g},jkl},\mu_{\mathrm{s},jkm}) = \begin{cases} \prod_{m=1}^{1} \mathrm{logit}^{-1}(\mu_{\mathrm{s},jkm}) \text{ if m=1} \\ \prod_{m=1}^{2}\prod_{l=1}^{1} \mathrm{logit}^{-1}(\mu_{\mathrm{s},jkm}) \times \left(1-\mathrm{logit}^{-1}(\mu_{\mathrm{g},jkl})\right) \text{ if } m=2 \\ \prod_{m=1}^{3}\prod_{l=1}^{1} \mathrm{logit}^{-1}(\mu_{\mathrm{s},jkm}) \times \left(1-\mathrm{logit}^{-1}(\mu_{\mathrm{g},jkl})\right) \text{ if } m=3 \\ \prod_{m=1}^{4}\prod_{l=1}^{2} \mathrm{logit}^{-1}(\mu_{\mathrm{s},jkm}) \times \left(1-\mathrm{logit}^{-1}(\mu_{\mathrm{g},jkl})\right) \text{ if } m=4 \\ \prod_{m=1}^{5}\prod_{l=1}^{2} \mathrm{logit}^{-1}(\mu_{\mathrm{s},jkm}) \times \left(1-\mathrm{logit}^{-1}(\mu_{\mathrm{g},jkl})\right) \text{ if } m=5 \\ \prod_{m=1}^{6}\prod_{l=1}^{3} \mathrm{logit}^{-1}(\mu_{\mathrm{s},jkm}) \times \left(1-\mathrm{logit}^{-1}(\mu_{\mathrm{g},jkl})\right) \text{ if } m=6 \end{cases}$$

$$\begin{aligned}
y_{\mathrm{p},3k} &= y_{\mathrm{p},3k}^{[t-1]} + y_{\mathrm{p},3k}^{[t-2]} \\
[\boldsymbol{\mu}_{\mathrm{s}}, \boldsymbol{\mu}_{\mathrm{s}}^{\mathrm{pop}}, \boldsymbol{\sigma}_{\mathrm{s}}^{\mathrm{pop}}, \boldsymbol{\mu}_{\mathrm{g}}, \boldsymbol{\mu}_{\mathrm{g}}^{\mathrm{pop}}, \boldsymbol{\sigma}_{\mathrm{g}}^{\mathrm{pop}}, \boldsymbol{\mu}_{\mathrm{s}_0}, \boldsymbol{\mu}_{\mathrm{s}_0}^{\mathrm{pop}}, \boldsymbol{\sigma}_{\mathrm{s}_0}^{\mathrm{pop}} &| \mathbf{y}_{\mathrm{g}}, \mathbf{y}] \propto \\
&\prod_{k=1}^{20}\prod_{j=1}^{3}\prod_{i=1}^{n_{jk}} \Bigg[ \Big[ \prod_{m=1}^{M} \mathrm{binomial}(y_{ijkm} | n_{ijkm}, f(\mu_{\mathrm{g},jkl}, \mu_{\mathrm{s},jkm})) \\
&\times \mathrm{normal}(\mu_{\mathrm{s},jkm} | \mu_{\mathrm{s},km}^{\mathrm{pop}}, \sigma_{\mathrm{s},km}^{\mathrm{pop}}) \\
&\times \mathrm{normal}(\mu_{\mathrm{s},km}^{\mathrm{pop}} | 0,1) \text{half-normal}(\sigma_{\mathrm{s},km}^{\mathrm{pop}} | 0,1) \Big] \\
&\Big[ \times \prod_{l=1}^{L} \mathrm{binomial}(y_{\mathrm{g},ijkl} | n_{\mathrm{g},ijkl}, \mathrm{logit}^{-1}(\mu_{\mathrm{g},jkl})) \\
&\times \mathrm{normal}(\mu_{\mathrm{g},jkl} | \mu_{\mathrm{g},kl}^{\mathrm{pop}}, \sigma_{\mathrm{g},kl}^{\mathrm{pop}}) \\
&\times \mathrm{normal}(\mu_{\mathrm{g},kl}^{\mathrm{pop}} | 0,1) \text{half-normal}(\sigma_{\mathrm{g},kl}^{\mathrm{pop}} | 0,1) \Big] \Bigg] \\
&\prod_{i=1}^{I}\prod_{j=1}^{J} \mathrm{binomial}(y_{\mathrm{p},3k}^{[t-1]} | n_{\mathrm{p},2k}, \mathrm{logit}^{-1}(\mu_{\mathrm{s}_0,2k}) f(\cdot, \mu_{\mathrm{s},\cdot 2k,m=1}) \mathrm{logit}^{-1}(\mu_{\mathrm{g},3k}) \\
&\times \mathrm{binomial}(y_{\mathrm{p},3k}^{[t-2]} | n_{\mathrm{p},3k}, \mathrm{logit}^{-1}(\mu_{\mathrm{s}_0,1k}) f(\mu_{\mathrm{g},\cdot 11}, \mu_{\mathrm{s},\cdot 1k,m=3}) \mathrm{logit}^{-1}(\mu_{\mathrm{g},3k}) \\
&\times \mathrm{normal}(\mu_{\mathrm{s}_0,jk} | \mu_{\mathrm{s}_0,k}^{\mathrm{pop}}, \sigma_{\mathrm{s}_0,k}^{\mathrm{pop}}) \\
&\times \mathrm{normal}(\mu_{\mathrm{s}_0,k}^{\mathrm{pop}} | 0,1) \text{half-normal}(\sigma_{\mathrm{s}_0,k}^{\mathrm{pop}} | 0,1).
\end{aligned} \tag{S11}$$

##### *S2.2.4 Viability*

We used the seed bag burial experiment to estimate the persistence of intact seeds in the field (Appendix S2.2.3). However, not all seeds that remain intact in the field are necessarily viable; some proportion may remain intact but be physiologically dead. Only some fraction of the seeds that are unearthed intact in the seed burial experiments are likely to be viable. To estimate the proportion of persistent, intact seeds from the seed bag burial experiment that are viable, we conducted lab assays on intact seeds when they were unearthed in October. As described in the main text, in October of each year of the seed bag burial experiment, the bags that had been dug up in the field were brought to the lab. The intact seeds remaining in the bag were tested in a two-stage lab viability trial. Specifically, we conducted lab germination trial and viability assays on subsets of the seeds from each bag to estimate the viability of the intact seeds.

The germination trials and viability assays form a sequence of binomial trials. Both the germination trials and viability assays are binomial trials, and we thus have counts of seeds at the start and end of each experiment. For each population, each models has the same structure for seeds from seed bag $i$ in experimental year $j$. If the number of seeds starting the trial (trials) is $n_{ij}$ and the number of seeds at the end of the trial (successes, either the number of seeds germinating or staining in the viability assay) is $y_{ij}$, we write a model that has a population-level mean and year-level means drawn from the population-level distribution. Broadly, this is two-level hierarchical model with a population-level mean, and year-level means drawn from the population-level distribution. The probability of success for each bag is drawn from this year- and population-level distribution. The model uses a binomial likelihood. The full models for germination trials (Fig. S6; Eq. S12) and viability assays (Fig. S6; Eq. S13) have a similar structure as the other models with binomial likelihoods (e.g., seedling survival to fruiting) and is written as

$$\begin{aligned}[\boldsymbol{\mu_G}, \boldsymbol{\mu_G^{\text{pop}}}, \boldsymbol{\sigma_G^{\text{pop}}} | \boldsymbol{y^{\text{tot}}}] \propto \prod_{i=1}^{I}\prod_{j=1}^{J} &\text{binomial}(y_{ij}^{\text{germ}} | n_{ijk}^{\text{test}_\text{g}}, \text{logit}^{-1}(\mu_{G,jk})) \\ &\times \text{normal}(\mu_{G,j} | \mu_{G,j}^{\text{pop}}, \sigma_{G,j}^{\text{pop}}) \\ &\times \text{normal}(\mu_{G,j}^{\text{pop}} | 0, 1)\text{half-normal}(\sigma_{G,j}^{\text{pop}} | 0, 1)\end{aligned} \tag{S12}$$

and

$$\begin{aligned}[\boldsymbol{\mu_V}, \boldsymbol{\mu_V^{\text{pop}}}, \boldsymbol{\sigma_V^{\text{pop}}} | \boldsymbol{y^{\text{tot}}}] \propto \prod_{i=1}^{I}\prod_{j=1}^{J} &\text{binomial}(y_{ij}^{\text{viab}} | n_{ij}^{\text{test}_\text{v}}, \text{logit}^{-1}(\mu_{V,j})) \\ &\times \text{normal}(\mu_{V,j} | \mu_{V,j}^{\text{pop}}, \sigma_{V,j}^{\text{pop}}) \\ &\times \text{normal}(\mu_{V,j}^{\text{pop}} | 0, 1)\text{half-normal}(\sigma_{V,j}^{\text{pop}} | 0, 1).\end{aligned} \tag{S13}$$

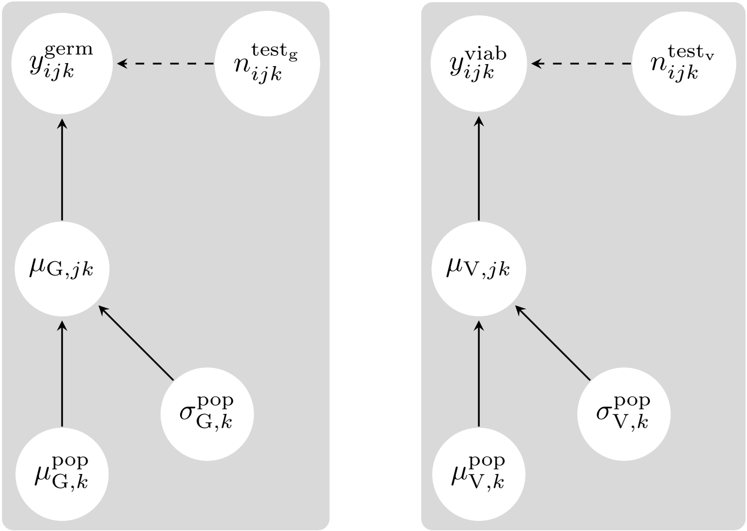

Figure S6: Directed acyclic graphs for the hierarchical models for lab trials. The left panel shows the DAG for the germination trials. The right panel shows the DAG for the viability assays.

### S3 Priors

Priors for all parameters in our statistical models are presented in Table S10. So that readers can understand the priors we used, we describe the principles we followed to place priors on parameters. First, we used weakly informative priors that avoid placing much probability on biologically unrealistic values (Gelman et al. 2017; Lemoine 2019; Wesner and Pomeranz 2021). Second, we placed positive, unbounded priors on variance terms, rather than priors with hard upper bounds (Gelman 2006). Third, we conducted prior predictive checks to confirm that the scale of priors translated to realistic values upon parameter transformation (Gabry et al. 2019; Hobbs and Hooten 2015; Wesner and Pomeranz 2021. Finally, we simulated prior predictive distributions to confirm that the joint likelihood generated data within the observed range of data (Conn et al. 2018; Gabry et al. 2019; Hobbs and Hooten 2015).

Table S10: Description of parameters in statistical models and associated prior distributions.

| Parameter | Description | Distribution |
|---|---|---|
| **Seed bag burial experiments** | | |
| $\mu^{\text{pop}}_{\text{g},jl}$ | Mean germination at population $k$ for seeds of age $l$ | normal(0, 1) |
| $\sigma^{\text{pop}}_{\text{g},jl}$ | S.D. of germination at population $k$ for seeds of age $l$ | half-normal(0, 1) |
| $\mu^{\text{pop}}_{\text{s},jm}$ | Mean seed survival at population $k$ for seeds in interval $m$ | normal(0, 1) |
| $\sigma^{\text{pop}}_{\text{s},jm}$ | S.D. of seed survival at population $k$ for seeds in interval $m$ | half-normal(0, 1) |
| $\mu^{\text{pop}}_{\text{s}_0,k}$ | Mean seed survival from July to October at population $k$ | normal(0, 1) |
| $\sigma^{\text{pop}}_{\text{s}_0,k}$ | S.D. of seed survival from July to October at population $k$ | half-normal(0, 1) |
| **Lab germination trials and viability assays** | | |
| $\mu^{\text{pop}}_{\text{G},k}$ | Mean germination in lab germination trials for seeds from population $k$ | normal(0, 1) |
| $\sigma^{\text{pop}}_{\text{G},k}$ | S.D. of germination in lab germination trials for seeds from population $k$ | half-normal(0, 1) |
| $\mu^{\text{pop}}_{\text{V},k}$ | Mean viability in lab viability assays for seeds from population $k$ | normal(0, 1) |
| $\sigma^{\text{pop}}_{\text{V},k}$ | S.D. of viability in lab viability assays for seeds from population $k$ | half-normal(0, 1) |
| **Seedling survival to fruiting** | | |
| $\mu^{\text{pop}}_{\text{S},k}$ | Mean seedling survival to fruiting at population $k$ | normal(0, 1) |
| $\sigma^{\text{pop}}_{\text{S},k}$ | S.D. of seedling survival to fruiting at population $k$ | half-normal(0, 1) |
| **Fruits per plant** | | |
| $\nu_{\text{TFE},k}$ | Mean total fruit equivalents per plant at population $k$ | gamma(1, 1) |
| $(\sigma_{\text{TFE},k})^2$ | Variance of total fruit equivalents per plant at population $k$ | half-normal(0, 1) |
| $(\sigma_{\text{TFE},k})^2$ | Variance of total fruit equivalents per plant at population $k$ | half-normal(0, 1) |
| $\nu_{\text{UF},k}$ | Mean undamaged fruits per plant at population $k$ | gamma(1, 1) |
| $(\sigma_{\text{UF},k})^2$ | Variance of undamaged fruits per plant at population $k$ | half-normal(0, 1) |
| $\nu_{\text{DF},k}$ | Mean damaged fruits per plant at population $k$ | gamma(1, 1) |
| $(\sigma_{\text{DF},k})^2$ | Variance of damaged fruits per plant at population $k$ | half-normal(0, 1) |
| **Seeds per fruit** | | |
| $\nu_{\text{US},k}$ | Mean seeds per undamaged fruit at population $k$ | gamma(1, 1) |
| $(\sigma_{\text{US},k})^2$ | Variance of seeds per undamaged fruit at population $k$ | half-normal(0, 1) |
| $\nu_{\text{DS},k}$ | Mean seeds per damaged fruit at population $k$ | gamma(1, 1) |
| $(\sigma_{\text{DS},k})^2$ | Variance of seeds per damaged fruit at population $k$ | half-normal(0, 1) |

### S4 Model convergence

We assessed convergence and mixing of the MCMC chains by (1) visually inspecting trace plots; by (2) calculating the Brooks-Gelman-Rubin diagnostic, $\hat{R}$; and by (3) calculating the Heidelberg-Welch diagnostic (Elderd and Miller 2016; Hobbs and Hooten 2015). Convergence refers to MCMC chains that start in different regions of parameter space honing in on the same range of values, and suggests that the chains are all sampling a high-likelihood region. Mixing refers to how the MCMC chain samples parameter space. A chain that exhibits good mixing moves around the posterior distribution, while a chain that exhibits poor mixing may get stuck at particular values or move around the posterior distribution slowly.

Trace plots are graphs that show the sequence of MCMC draws: iterations in the MCMC sampler are shown on the x-axis and parameter values are shown on the y-axis. When multiple chains are plotted together, it is possible to visually assess whether the chains in an MCMC run are converged and well mixed. To evaluate convergence, it is necessary to compare the behavior of multiple chains; a MCMC run that exhibits convergence will have multiple chains that broadly overlap in the trace plots; a run that does not exhibit convergence will not show overlap. Mixing can be evaluated at the level of individual chains. Chains that are well mixed will show few signs of auto-correlation; chains that are poorly mixed will exhibit auto-correlation or remain stuck around particular parameter values.

In addition to visual assessment, the Brooks-Gelman-Rubin diagnostic, $\hat{R}$, assesses convergence between chains, and the Heidelberg-Welch diagnostic evaluates stationarity within chains. The Brooks-Gelman-Rubin diagnostic, $\hat{R}$, is a ratio that compares variability between versus within chains: values of $\hat{R}$ close to 1 suggest convergence. Because variability between chains is typically greater than or equal to variability within chains, the lower limit of values is typically 1. Stationarity is closely related to whether a chain exhibits good mixing. Informally, the Heidelberg-Welch diagnostic tests whether a chain is stationary by checking whether the mean remains relatively unchanged across the MCMC chain.

Trace plots generally reflected convergence and good mixing across the different models that we fit (not shown). Because we fit separate models for each population, and some models contained parameters for multiple processes, we inspected many individual trace plots. To summarize the patterns in the trace plots, we used the $\hat{R}$ and Heidelberg-Welch diagnostic. Specifically, we collected all the parameters for a given dataset (e.g. all means and standard deviation parameters in the seedling survival to fruiting model) from all populations. For each parameter set, we calculated and plotted the distribution of $\hat{R}$, and calculated the percentage of these $\hat{R}$ values that were less than 1.05. We also calculated the Heidelberg-Welch diagnostic for the chains in this parameter set, and calculated the percentage of chains that were stationary. We used a p-value cutoff of 0.05 to determine whether a chain failed the Heidelberg-Welch diagnostic. With the repeated testing that we conducted in calculating this diagnostic for many chains, we would expect 5% of stationary chains to fail under this diagnostic.

Histograms of $\hat{R}$ values show that chains converged, with $\hat{R}$ for all chains less than 1.05. In addition, the distribution of $\hat{R}$ was skewed to the right in models for all datasets. The majority of $\hat{R}$ values were close to 1 and larger values were always part of a tail. Summaries of the Heidelberg-Welch diagnostic indicate that chains were well-mixed in almost all cases. The sole exception was that only 93% of one of the chains (chain 3) passed the Heidelberg-Welch diagnostic in the model for seeds per undamaged fruit.

To determine whether this was an issue that could complicate inferences about the data, we examined this model in greater detail. All the chains that failed the diagnostic test were associated with the parameter for the population-level variance or the variance for the observation-level term for overdispersed counts. We suspect that it may be challenging to sample the posterior for these parameters in some populations because of variability in sample sizes on seeds per undamaged fruit. It could be especially challenging to sample the posterior distribution for variance terms if certain years are outliers in terms of their sample variance due to small sample sizes.

In summary, we retained these models because the vast majority of diagnostic tests indicated convergence and mixing. However, we note that a small handful of variance terms exhibited poor mixing and that the models could be reevaluated if we the goal were to make inferences about the variance in seeds per undamaged fruit.

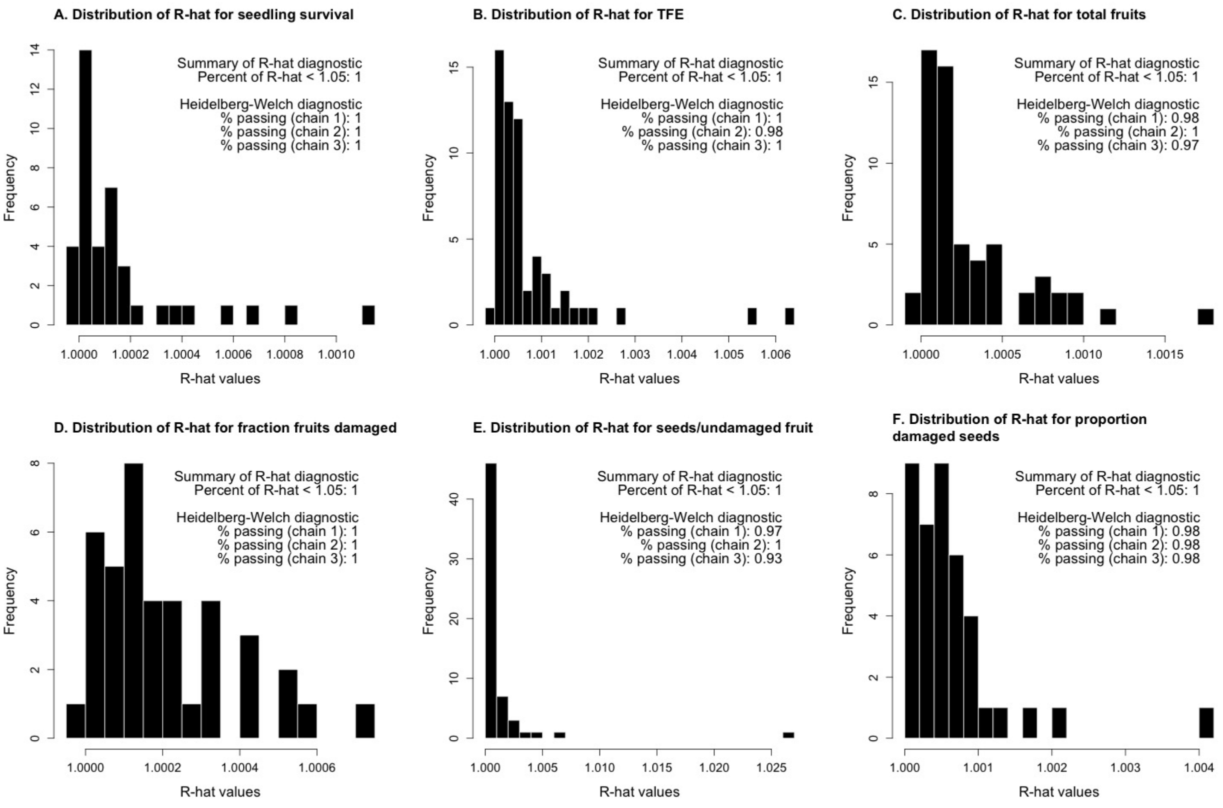

Figure S7: Histograms of $\hat{R}$ values for parameters associated with models for different datasets. In each panel, the histogram plots the frequency of $\hat{R}$ values. The inset text summarizes the percent of $\hat{R}$ values that are less than 1.05, and the percent of each chain that passes the Heidelberg-Welch diagnostic. See text for details. (A) Diagnostics for parameters from the model for seedling survival to fruiting. (B) Diagnostics for parameters associated with the model for total fruit equivalents per plant (2006-2012). (C) Diagnostics for parameters associated with the model for total fruits per plant (2013-2020). (D) Diagnostics for parameters associated with the model for fraction of total fruits per plant that are damaged (2013-2020). (E) Diagnostics for parameters associated with the model for number of seeds per undamaged fruit (2006-2020). (F) Diagnostics for parameters associated with the model for proportion of seeds lost to damage in damaged fruit (2013-2020).

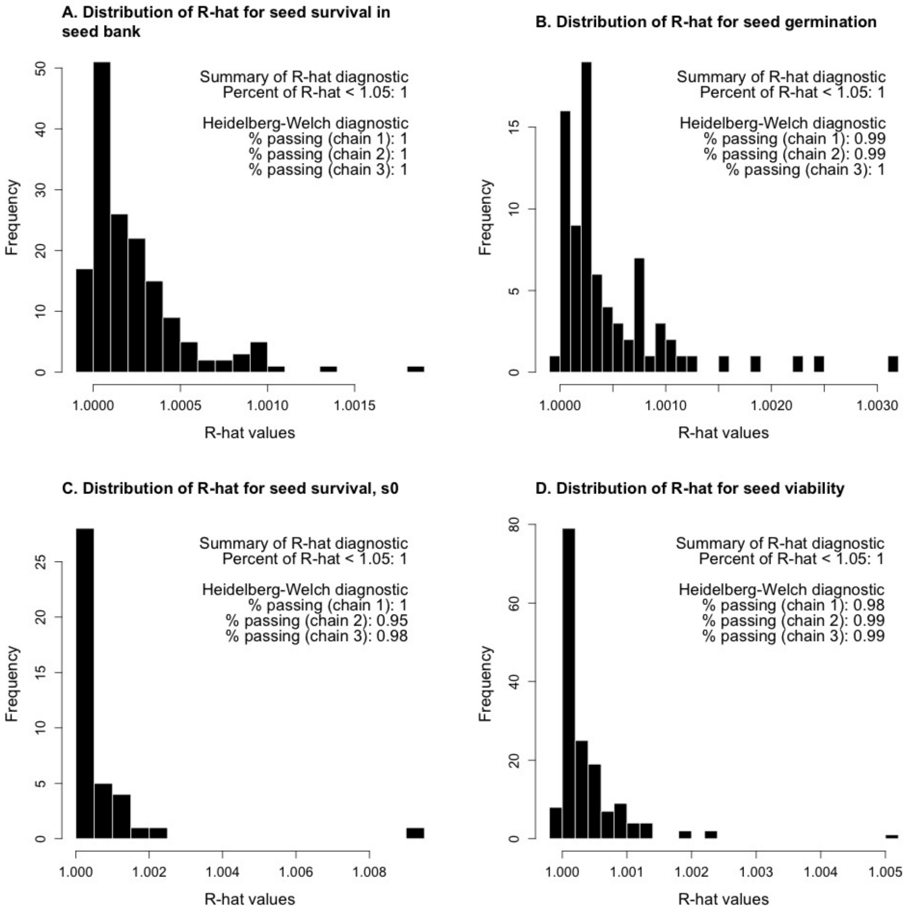

Figure S8: Histograms of $\hat{R}$ values for parameters associated with models for different datasets. In each panel, the histogram plots the frequency of $\hat{R}$ values. The inset text summarizes the percent of $\hat{R}$ values that are less than 1.05, and the percent of each chain that passes the Heidelberg-Welch diagnostic. See text for details. (A) Diagnostics for parameters from the model for seed survival in the seed bank. (B) Diagnostics for parameters associated with the model for germination. (C) Diagnostics for parameters associated with the model for seed survival, $s_0$. (D) Diagnostics for parameters associated with the model for data from the viability trials (including the viability and germination tests).

### S5 Computing vital rates

#### *S5.1 Per-capita reproductive success*

To make our analysis comparable to previous empirical studies of bet hedging, we calculated per capita reproductive success as the product of the probability of seedling survival to fruiting, fruits per plant, and seeds per fruit. We thus calculate per capita reproductive success as the number of seeds produced per seedling, on average (Gremer and Venable 2014; Venable 2007).

We used a consistent method to estimate seedling survival to fruiting throughout the experiment, and use the population- and year-level means ($\mu_{S,jk}$) in our calculation. Because we estimated fruit production in 2 different ways during the study, we chose to use total fruit equivalents (TFE) per plant as our common estimate of fruit production. From 2006–2012, we used $\mu_{TF,jk}$ as estimated in the statistical model. From 2013–2020, we used the ratio of seeds per damaged to undamaged fruit to calculate a proportion of damaged fruits to add to undamaged fruit counts, as in

$$\text{TFE} = \text{undamaged fruits} + \frac{\text{seeds per damaged fruit}}{\text{seeds per undamaged fruit}} \times \text{damaged fruits}. \tag{S14}$$

We used posterior distributions for population- and year-level parameters (e.g., $\mu_{US,jk}$) for these calculations and obtained $\mu_{TOT,jk}$ for 2013–2020. Finally, we used estimates of seeds per undamaged fruit ($\mu_{US,jk}$) as our estimate of seeds per fruit.

In terms of parameters from our statistical models, per capita reproductive success $F_{jk}$ in year $j$ at population $k$ is calculated as

$$F_{jk} = \phi_{jk} \times \lambda_{TFE,jk} \times \lambda_{US,jk} \tag{S15}$$

where

$$\begin{aligned} \phi_{jk} &= \text{logit}^{-1}(\mu_{\text{S},jk}), \\ \lambda_{\text{TFE},jk} &= \exp(\mu_{\text{TFE},jk}), \text{and} \\ \lambda_{\text{US},jk} &= \exp(\mu_{\text{US},jk}). \end{aligned} \tag{S16}$$

#### S5.2 Belowground vital rates

To calculate seed survival and germination for the population model, we combine parameter estimates from the statistical models for observations from the seed bag burial experiments and lab germination trials and viability assays. Importantly, the estimates of seed persistence and emergence from the seed bag burial experiment do not account for loss of seed viability because we cannot distinguish seeds that are intact and viable versus seeds that are intact but not viable in the field. We thus calculate seed viability by combining information the lab germination trials and viability assays (Appendix S5.2.1). Next, we obtain estimates of seed persistence and emergence that correspond to the times (October, January) at which we make observations in the seed bag burial experiment (Appendix S5.2.2). Then, we calculate *seed survival* by combining estimates for seed persistence (from the seed bags in the field) and seed viability (from the lab germination trials and viability assays) (Appendix S5.2.3). Seed survival accounts for (1) the loss of intact seeds from the soil seed bank and (2) the loss of seeds due to physiological death. Similarly, we calculate *germination* by combining estimates for emergence (from the seed bags in the field) and seed viability (from the lab germination trials and viability assays).

##### *S5.2.1 Calculating viability with lab germination trials and viability assays*

We use the viability trials to calculate the probability of viability ($\nu_1$ or $\nu_2$) for a seed that is intact in October one or two years, respectively, after the seed bags were buried. We use these overall estimates of viability, which combine information from both stages of the viability trial, to account for loss of seed viability in the seed burial experiments. The viability trails are a two-stage process in which seeds are subject to germination trials before a fraction of the remaining, ungerminated seeds are assayed for viability. The probability of viability is thus calculated using both pieces of the two-stage experiment.

We write the probability that a seed germinates in the germination trial as $\theta_{\mathrm{G}}$ and the probability that it is viable in the viability trial, conditional on not germinating in the germination trial, as $\theta_{\mathrm{V}}$. We obtain the posterior distributions of these parameters by marginalization. We transform the posteriors to $[0, 1]$ by taking the inverse logit; this transforms the parameters into the probability of success. We estimate the overall probability of viability, $\nu_1$ or $\nu_2$, as $\theta_{\mathrm{G}} + \theta_{\mathrm{V}}(1 - \theta_{\mathrm{G}})$. This weights the estimates relative to the probability of germination (eg. if no seeds germinate the estimate of viability will mostly come from the viability test).

In terms of parameters from our statistical models, overall probability of viability $\nu_{1,k}$ at population $k$ is calculated as

$$\nu_{1,k} = \theta_{\mathrm{G,k}} + \theta_{\mathrm{V,k}}(1 - \theta_{\mathrm{G,k}}), \tag{S17}$$

where

$$\begin{aligned} \theta_{\mathrm{G}} &= \mathrm{logit}^{-1}(\mu_{\mathrm{G},1k}) \\ \theta_{\mathrm{V}} &= \mathrm{logit}^{-1}(\mu_{\mathrm{V},1k}). \end{aligned} \tag{S18}$$

In terms of parameters from our statistical models, overall probability of viability $\nu_{2,k}$ at population $k$ is calculated as

$$\nu_{2,k} = \theta_{\mathrm{G,k}} + \theta_{\mathrm{V,k}}(1 - \theta_{\mathrm{G,k}}), \tag{S19}$$

where

$$\begin{aligned} \theta_{\mathrm{G}} &= \mathrm{logit}^{-1}(\mu_{\mathrm{G},2k}) \\ \theta_{\mathrm{V}} &= \mathrm{logit}^{-1}(\mu_{\mathrm{V},2k}). \end{aligned} \tag{S20}$$

We tested the viability of seeds in October, and were thus able to estimate the proportion of viable seeds at that time. We use the overall viability estimates ($\nu_1$) from October to calculate viability for the preceding January (9 months prior) by interpolation. We inferred the viability of intact seeds in January by assuming that seeds lost viability at a constant rate (exponential decay). In January one year after seed burial, the viability of seeds was thus estimated as $\nu_1^{1/3}$. Further, we interpolated between years 1 and 2 of the experiment by assuming that viability changed at a constant rate between years, and that all seeds were viable at the start of the experiment. In January two years after seed burial, the viability of seeds was thus estimated as $\nu_1(\nu_2/\nu_1)^{1/3}$.

##### S5.2.2 *Estimates for seed persistence and emergence*

We used the seed bag burial experiment to estimate seed persistence and emergence. For seed persistence, we estimated the probability that a seed remained intact between the times at which we unearthed the seed bags and counted intact seeds. We calculated the following values: (1) the probability that seeds remained intact from the start of the seed bag burial experiment in October to the first January, $\theta_{\cdot\cdot 1}$; (2) the probability that seeds remained intact from the first January to 8 months later in October, $\theta_{\cdot\cdot 2}$; and (3) the probability that seeds remained intact from the second October of the experiment to January, $\theta_{\cdot\cdot 3}$. Estimates of persistence over these intervals are the probability that a seed remains intact.

In terms of parameters from our statistical models, the probabilities of seed persistence in year $j$ at population $k$ are calculated as

$$\begin{aligned}\theta_{jk1} &= \text{logit}^{-1}(\mu_{\text{s},jk1})\\ \theta_{jk2} &= \text{logit}^{-1}(\mu_{\text{s},jk2})\\ \theta_{jk3} &= \text{logit}^{-1}(\mu_{\text{s},jk3}).\end{aligned} \tag{S21}$$

Similarly, we calculated the probability that a seed that is intact in the first January of the seed bag burial experiment emerges as a seedling, $\gamma_1$, and the probability that a seed that is intact in the second January of the seed bag burial experiment emerges as a seedling, $\gamma_2$. In terms of parameters from our statistical models, the probabilities of seed persistence year $j$ at population

$k$ are calculated as

$$\begin{aligned}\gamma_{jk1} &= \text{logit}^{-1}(\mu_{\text{g},1}), \text{and} \\ \gamma_{jk2} &= \text{logit}^{-1}(\mu_{\text{g},2}).\end{aligned} \tag{S22}$$

Finally, we calculated the $s_0$, the probability that seeds produced in July of year $t$ survived to October of year $t$. Because we assume that all seeds are viable at the start of the seed bag experiment, the estimate for $s_0$ is not subsequently adjusted for loss of seed viability. In terms of parameters from the statistical model,

$$s_0 = \text{logit}^{-1}(\mu_{s_0,jk}). \tag{S23}$$

##### *S5.2.3 Calculating seed survival and germination by combining information from seed bag burial experiments with viability estimates*

We calculated seed survival and germination with the equations given in Table S11. Each equation defines the survival and germination probabilities for the population model in terms of the values for persistence ($\theta$s), emergence ($\gamma$s), and viability ($\nu$s). We seek to transform estimates for persistence and emergence (which *do not* account for loss of viability) to estimates for survival and germination (which *do* account for loss of viability). To accomplish this, we adjust persistence and emergence to account for the loss of viability that seeds may experience. We do this by combining the posteriors for the persistence and emergence estimates with posteriors for the viability estimates according to the equations in Table S11.

For example, survival from the first October to the first January, $s_1$, is calculated as the product of $\theta_1$ with the sum of the probability of emergence and the probability of viability for a seed that does not emerge. To understand the motivation for these equations, consider the limiting cases in which all or no intact seeds are viable. If all intact seeds that do not germinate are viable, then survival is the product of persistence times one. On the other hand, if no intact seeds that do not germinate are viable, then survival is the product of persistence times emergence because only those seeds that emerged were alive. In general, we calculate seed survival by discounting the portion of persisting (intact) seeds that are not viable. Similarly, we calculate germination by discounting the portion of persisting (intact) seeds that are not viable and thus could not have become seedlings.

To calculate the parameters for the population model, we combine posterior distributions of parameters from the statistical models. Because we combine these estimates from multiple years of observations, the ratio of some posterior distributions is greater than 1. We restricted the posterior to be less than 1 by truncating the distribution and resampling to redistribute the probability mass. We take this step to retain parameter uncertainty about survival probability in cases where combining the estimates implies a high probability of survival.

Table S11: Belowground vital rate components of the population model.

| Description | Parameter | Probability |
|---|---|---|
| Probability that a seed produced in July of year $t$ is intact and viable in October of year $t$ | $s_0$ | $s_0$ |
| Probability that a seed survives from October of year $t$ to January of year $t+1$, for seeds produced in year $t$ | $s_1$ | $\theta_1 \times (\gamma_1 + (1-\gamma_1) \times \nu_1^{1/3})$ |
| Probability of germination for a seed that has survived to January of year $t+1$, for seeds produced in year $t$ | $g_1$ | $\frac{\gamma_1}{1-(1-\nu_1^{1/3})\times(1-\gamma_1)}$ |
| Probability that a seed survives from January of year $t+1$ to October of year $t+1$, for seeds produced in year $t$ | $s_2$ | $\theta_2 \nu_1^{2/3}$ |
| Probability that a seed survives from October of year $t+1$ to January of year $t+2$, for seeds produced in year $t$ | $s_3$ | $\frac{\theta_1(1-\gamma_1)\theta_2\theta_3\times(\gamma_2+(1-\gamma_2)\times\nu_1^{1/3}(\nu_2/\nu_1)^{1/3})}{s_1\times(1-g_1)\times s_2}$ |

#### *S5.3 Graphical summary of vital rate parameters*

We graphically summarize the vital rate parameters used in the analysis in this text, all of which are described in Table 1. Per-capita reproductive success is the product of seedling survival to fruiting, $\sigma$ (Fig. S9); fruits per plant, $F$ (Fig. S10)); and seeds per fruit, *phi* (Fig. S11)). For fruits per plant $F$, annual estimates from 2006-2012 (shown on gray background) are from models for counts of total fruit equivalents per plant. Annual estimates from 2013-2020 are calculated by combining estimates of undamaged and damaged fruits per plant with an estimate of the fraction of seeds lost to fruit damage.

Seed survival from seed production in the summer to the first January/February is described by the product of $s_0$ (Fig. S12) and $s_1$ (Fig. S13). Germination is given by $g_1$ (Fig. S14). Seed survival in the seed bank, from January/February in year $t$ to January/February in year $t$, is the product of $s_2$ (Fig. S15) and $s_3$ (Fig. S16).

For seed survival $s_0$, annual estimates for year $t$ are for the late summer/fall in year $t$, the year in which seeds were produced. For example, estimates from for $s_0$ in 2006 are for seed survival from seed production in late June 2006 to October 2006, for seeds produced in summer 2006. For $s_0$, the annual estimates are not centered on the population in several cases (e.g., LO in Fig. S12). The reason is that in order to model $s_0$, we aggregated observations to the level of transects, and used estimates of seed rain from 2006 and 2007 with seedling counts in 2008 (see Appendix S2.2.3). In comparison to the other vital rates in the study, we thus have less data to inform estimates of $s_0$. As a result, the weakly informative prior that we use (see Appendix S3), which is centered on a probability of survival of 0.5, has weak influence on the posterior in this model.

For seed survival $s_1$, annual estimates for year $t$ are for seed survival from October in year $t-1$ to January in year $t$, for seeds produced in summer of year $t-1$. For example, estimates from for $s_1$ in 2006 are for seed survival from October 2005 to January 2006, for seeds produced in summer 2005. For germination, $g_1$, annual estimates for year $t$ are for January in year $t$, for seeds produced in the summer of year $t-1$. For example, estimates for $g_1$ in 2006 are for seeds that germinated in January 2006, for seeds produced in summer 2006.

For seed survival $s_2$, annual estimates are for seed survival from January to October in year $t$, for seeds produced in summer of year $t-1$. For example, estimates for $s_2$ in 2006 are for seed survival from January 2006 to October 2006, for seeds produced in summer 2005. For seed survival $s_3$, annual estimates for year $t$ are for seed survival from October in year $t-1$ to January in year $t$, for seeds that produced in summer of year $t-2$. For example, estimates for $s_3$ in 2007 are for seed survival from October 2006 to January 2007, for seeds produced in summer 2005.

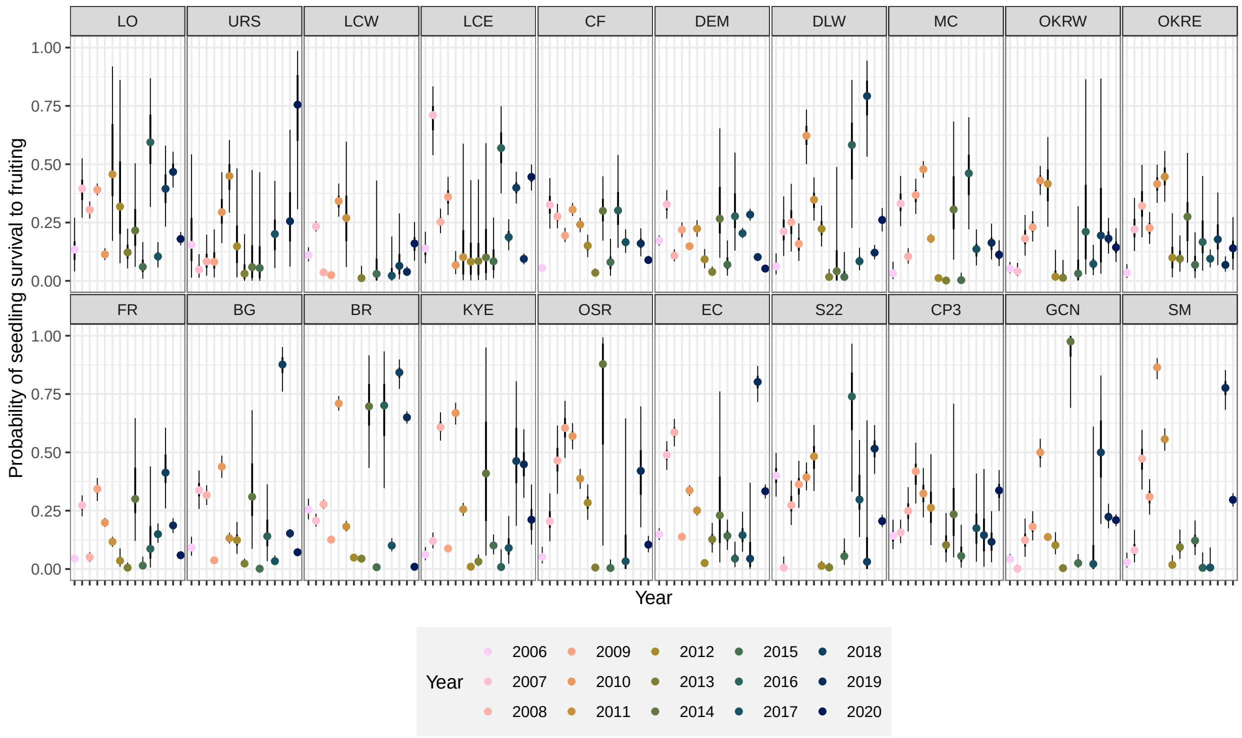

Figure S9: Probability of seedling survival to fruiting, $\sigma$. The points are the posterior modes for the annual estimates, and the thick and thin error bars are, respectively, the 50% and 95% highest posterior density intervals for the annual estimates. Populations are arrayed by easting from west to east, with the most western populations at the top left and the most eastern populations at the bottom right.

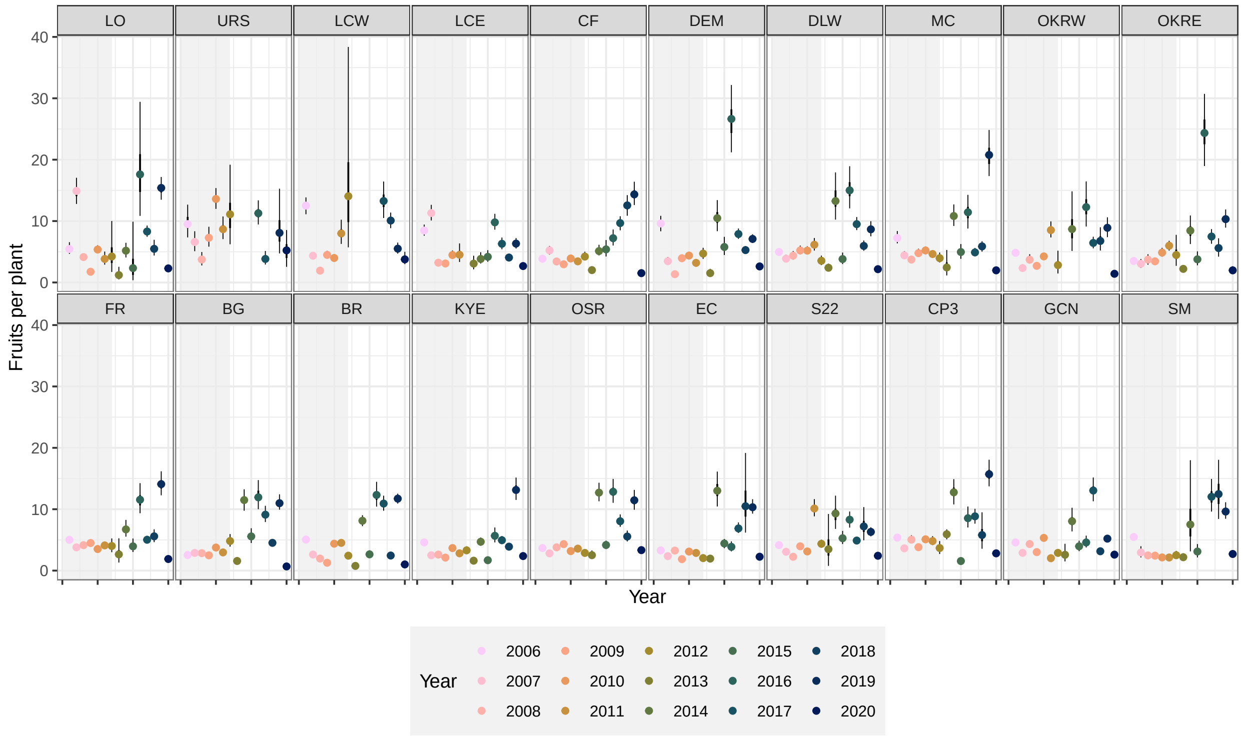

Figure S10: Fruits per plant, $F$. From 2006-2012, we estimated total fruit equivalents per plant directly from observations in the field. From 2013-2020, we calculated total fruit equivalents per plant by combining estimates of undamaged and damaged fruits per plant with an estimate of the fraction of seeds lost to fruit damage. The points are the posterior modes for the annual estimates, and the thick and thin error bars are, respectively, the 50% and 95% highest posterior density intervals for the annual estimates. Populations are arrayed by easting from west to east, with the most western populations at the top left and the most eastern populations at the bottom right.

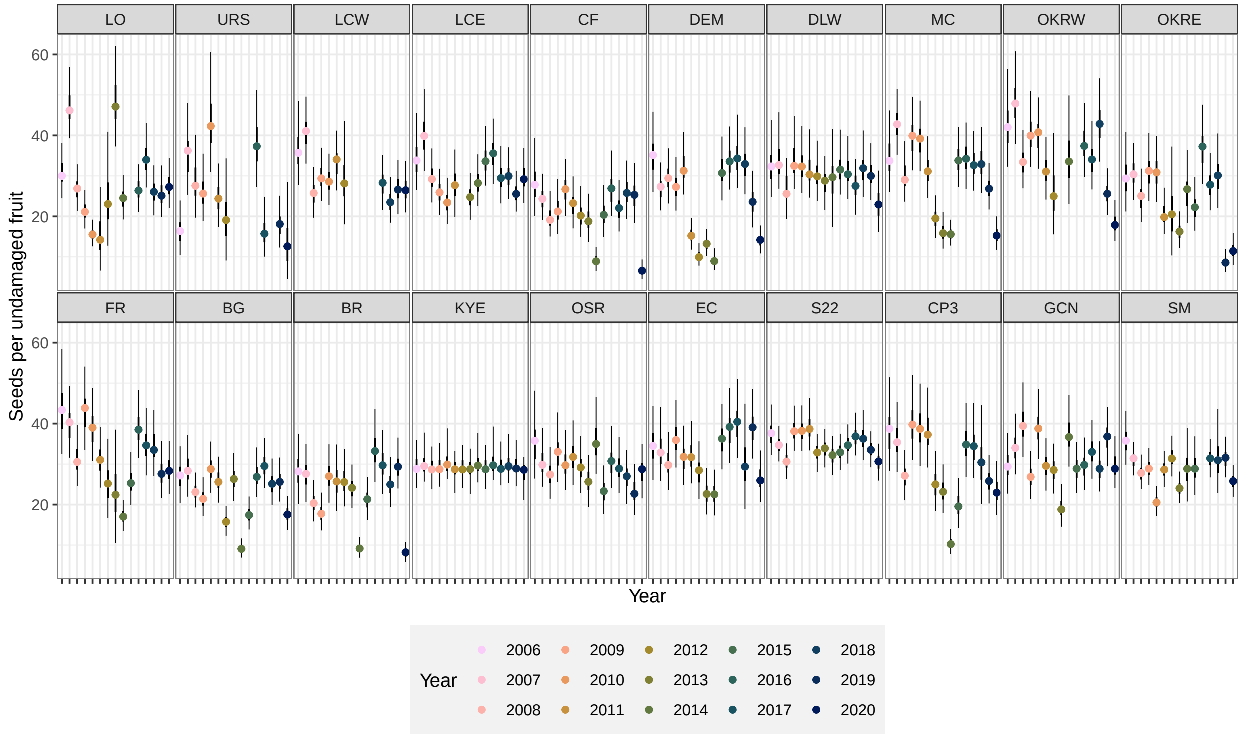

Figure S11: Seeds per undamaged fruit, $\phi$. The points are the posterior modes for the annual estimates, and the thick and thin error bars are, respectively, the 50% and 95% highest posterior density intervals for the annual estimates. Populations are arrayed by easting from west to east, with the most western populations at the top left and the most eastern populations at the bottom right.

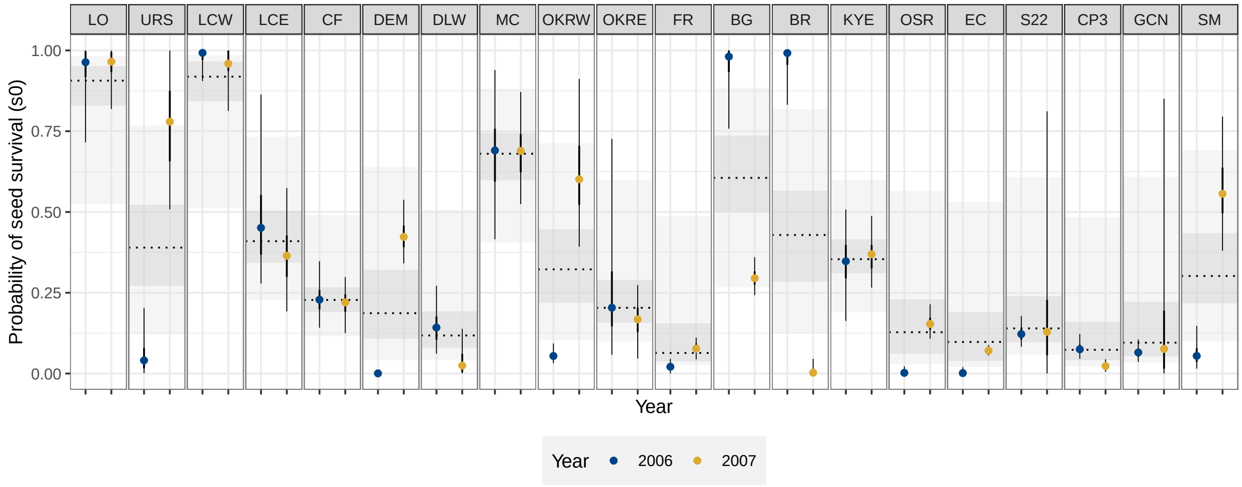

Figure S12: Probability of seed survival, $s_0$. In each panel, the dotted line shows the posterior mode for the population-level estimate, while the dark and light gray shading show the corresponding 50% and 95% highest posterior density intervals. The points are the posterior modes for the annual estimates, and the thick and thin error bars are, respectively, the 50% and 95% highest posterior density intervals for the annual estimates. Populations are arrayed by easting from west to east, with the most western populations at the far left and the most eastern populations at the far right.

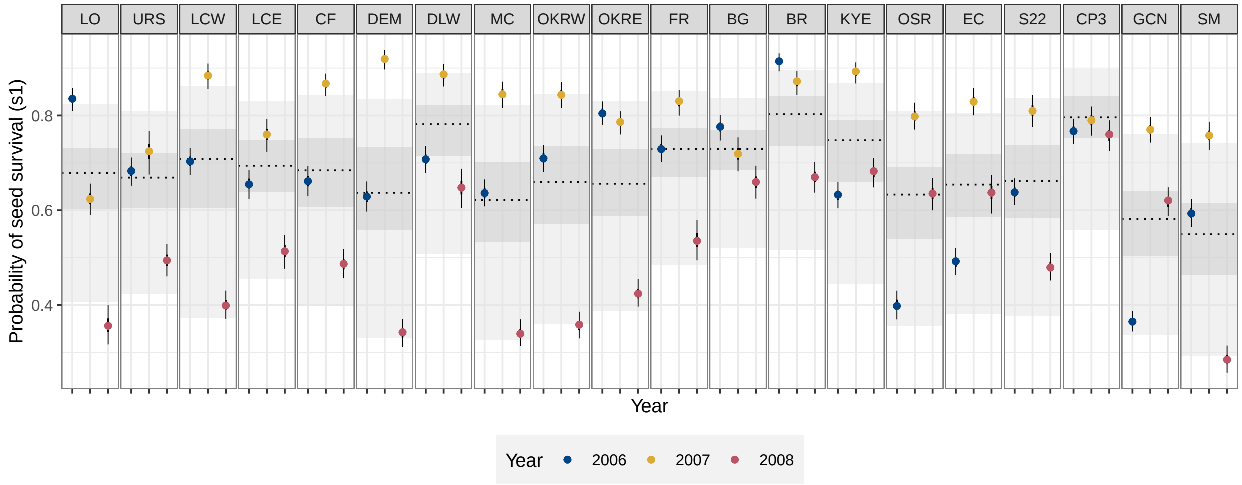

Figure S13: Probability of seed survival, $s_1$. In each panel, the dotted line shows the posterior mode for the population-level estimate, while the dark and light gray shading show the corresponding 50% and 95% highest posterior density intervals. The points are the posterior modes for the annual estimates, and the thick and thin error bars are, respectively, the 50% and 95% highest posterior density intervals for the annual estimates. Populations are arrayed by easting from west to east, with the most western populations at the far left and the most eastern populations at the far right.

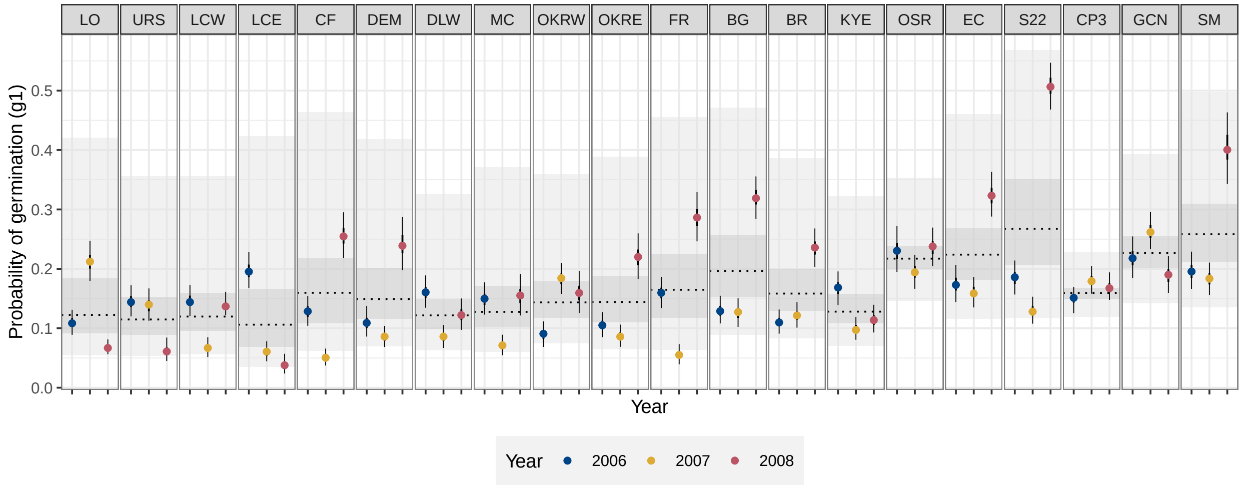

Figure S14: Probability of seed survival, $g_1$. In each panel, the dotted line shows the posterior mode for the population-level estimate, while the dark and light gray shading show the corresponding 50% and 95% highest posterior density intervals. The points are the posterior modes for the annual estimates, and the thick and thin error bars are, respectively, the 50% and 95% highest posterior density intervals for the annual estimates. Populations are arrayed by easting from west to east, with the most western populations at the far left and the most eastern populations at the far right.

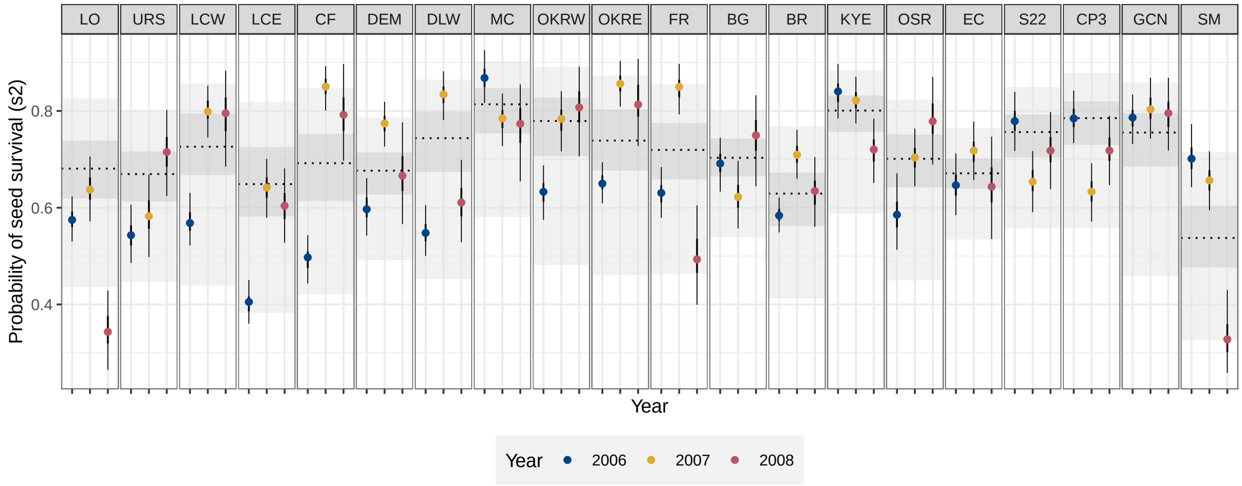

Figure S15: Probability of seed survival, $s_2$. In each panel, the dotted line shows the posterior mode for the population-level estimate, while the dark and light gray shading show the corresponding 50% and 95% highest posterior density intervals. The points are the posterior modes for the annual estimates, and the thick and thin error bars are, respectively, the 50% and 95% highest posterior density intervals for the annual estimates. Populations are arrayed by easting from west to east, with the most western populations at the far left and the most eastern populations at the far right.

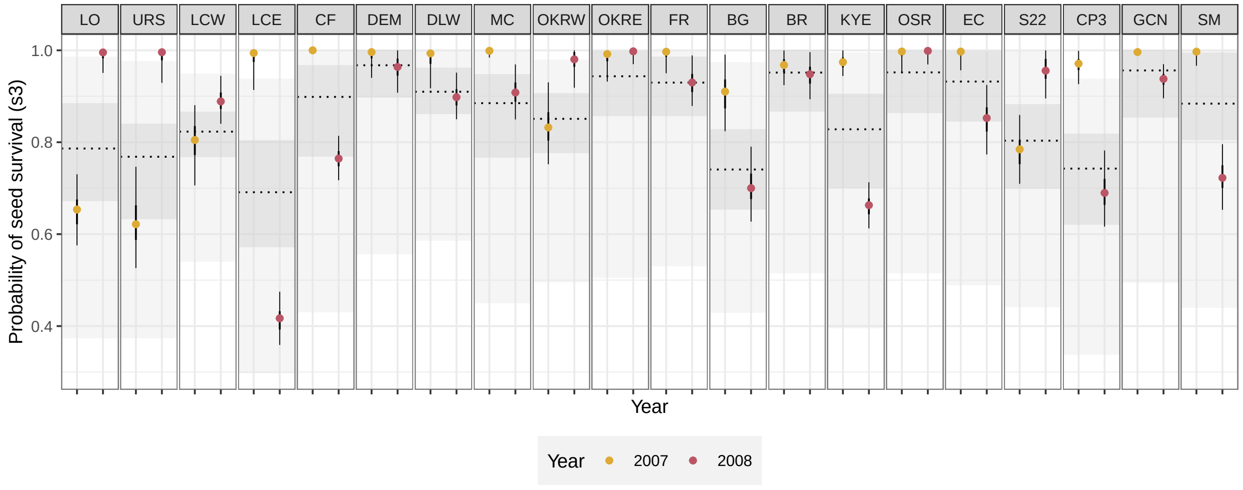

Figure S16: Probability of seed survival, $s_3$. In each panel, the dotted line shows the posterior mode for the population-level estimate, while the dark and light gray shading show the corresponding 50% and 95% highest posterior density intervals. The points are the posterior modes for the annual estimates, and the thick and thin error bars are, respectively, the 50% and 95% highest posterior density intervals for the annual estimates. Populations are arrayed by easting from west to east, with the most western populations at the far left and the most eastern populations at the far right.

### S6 Supplementary results and analysis

#### *S6.1 Magnifying panel B in results of demographic test of bet hedging*

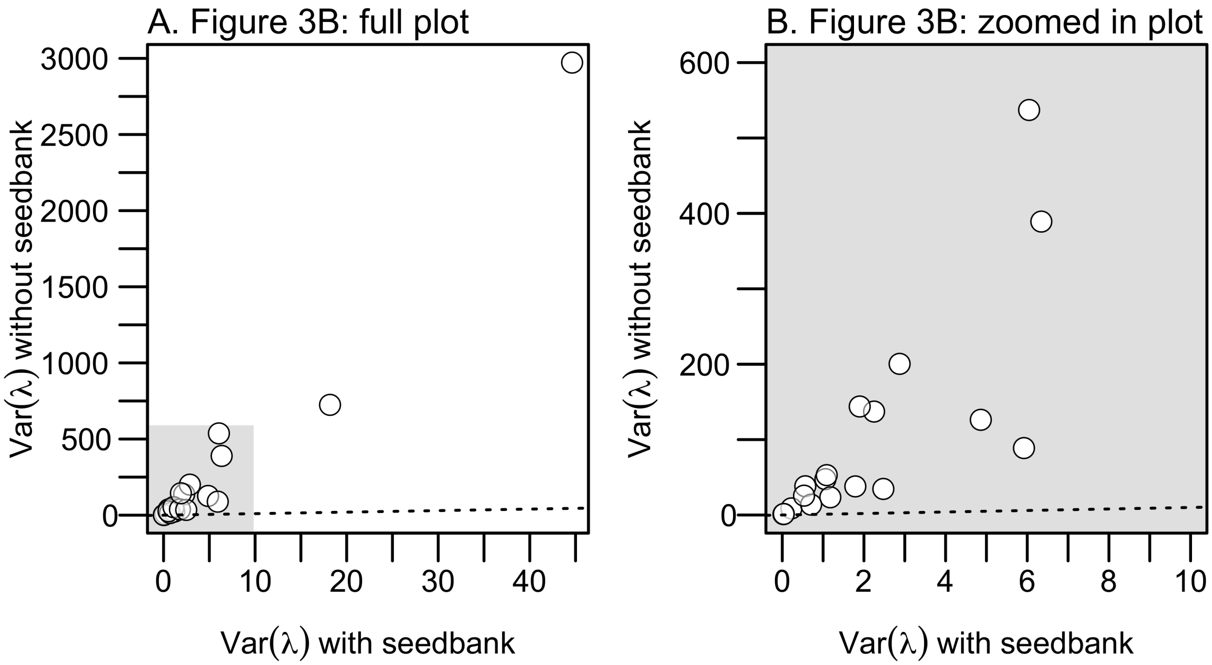

Figure S17: Plots of the variance in annual population growth rate without a seed bank against the variance in population growth rate with a seed bank. (A) The panel is identical to Figure 3B in the main text, with the addition of a gray area that highlights the region in the plot that is magnified in panel B. (B) The panel zooms in on the gray area in panel A to magnify the points that lie closest to the origin. In both plots, the dotted line is the 1:1 line.

#### *S6.2 Demographic test of bet hedging with parameter uncertainty*

To account for parameter uncertainty in the demographic test of bet hedging, we repeated the demographic test with samples from the posterior. In the main analysis, we used the posterior modes of per capita reproductive success to obtain 15 estimates for reproductive success, $Y(t)$, and the posterior modes of seed survival and germination to obtain population-level estimates of seed survival and germination. To incorporate uncertainty about our parameter estimates, we drew samples from the posterior distribution (e.g., Elderd and Miller 2016; Evans et al. 2010). For example, we collected 1 draw of 15 estimates of per capita reproductive success, and 1 draw of estimates for seed survival and germination. We used this draw to calculate arithmetic mean growth rate, and resample 1,000 years of per capita reproductive success and calculate long-term stochastic population growth rate and variability in population growth. We repeated this process for all 45,000 samples of the posterior distribution. This gave us 45,000 estimates for the arithmetic mean growth rate, variability in population growth, and stochastic population growth. We summarized these estimates with the posterior mode and highest posterior density interval. Results based on the posterior modes proved to be robust to parameter uncertainty (Fig. S18), such that the absence of evidence for a trade-off between arithmetic and geometric mean fitness is unlikely explained by uncertainty about parameter estimates.

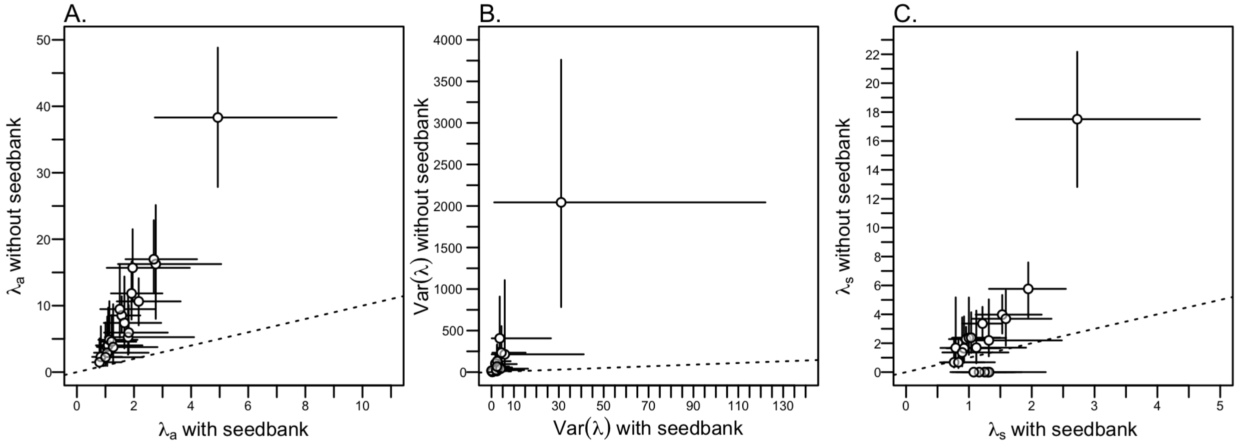

Figure S18: Test of three demographic patterns expected with bet hedging, accounting for parameter uncertainty. (A) Plots of the arithmetic population growth rate without a seed bank against arithmetic population growth with a seed bank. (B) Plots of the variance in annual population growth rate without a seed bank against the variance in population growth rate with a seed bank. (C) Plots of the long-term stochastic population growth rate without a seed bank against the long-term stochastic growth rate without a seed bank. In all plots, the dotted line is the 1:1 line. The points are the posterior mode. The error bars are the 68% highest posterior density intervals (under a normal distribution, 68% of the distribution is within 1 standard deviation).

#### *S6.3 Demographic test of bet hedging with quasi-complete germination*

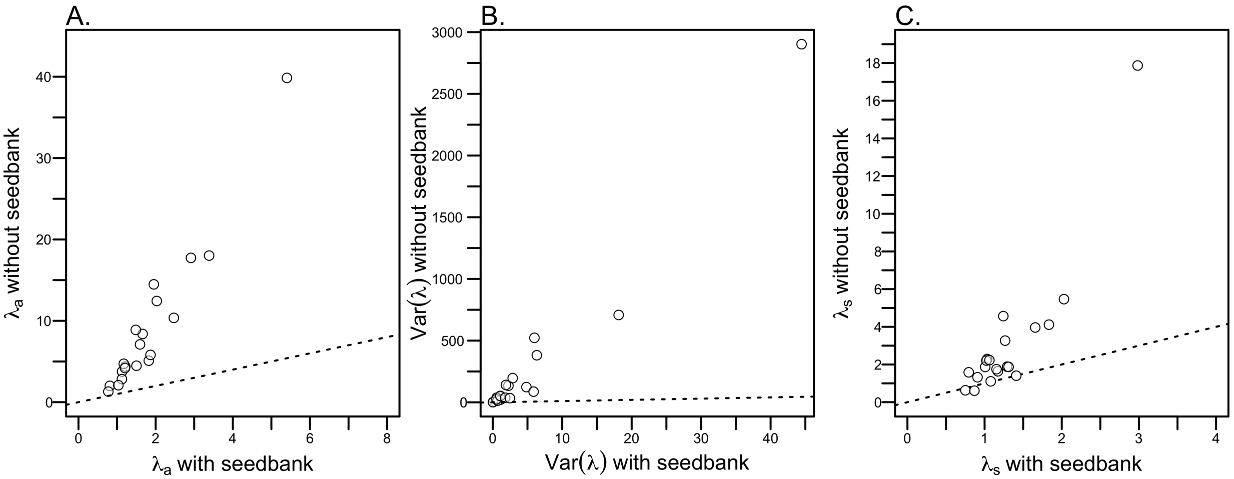

Figure S19: Test of the demographic patterns expected with bet hedging with a quasi-complete germination. In the analysis presented here, we assumed that almost all seeds germinate ($g_1 = 0.99$) and there is a minimal seed bank. In the analysis presented in the main text, we assumed that all seeds germinate ($g_1 = 1$) and there is no seed bank. (A) Plot of the arithmetic population growth rate with a minimal seed bank (quasi-complete germination) against arithmetic population growth with a seed bank. (B) Plots of the variance in annual population growth rate with a minimal seed bank (quasi-complete germination) against the variance in population growth rate with a seed bank. (C) Plot of the long-term stochastic population growth rate with a minimal seed bank (quasi-complete germination) against the long-term stochastic growth rate without a seed bank. In all plots, the dotted line is the 1:1 line.

#### S6.4 *Accounting for parameter uncertainty in optimal germination fractions*

To account for the effect of parameter uncertainty on the predicted, optimal germination fractions, we found optimal germination fractions for random samples from the posterior distribution. In the analysis in the main text, we used the posterior modes of per capita reproductive success and seed survival to calculate the optimal germination fraction by resampling the annual estimates of per capita reproductive success, $Y(t)$. Instead of conducting this analysis with the posterior modes, we did this analysis with draws from the posterior distribution.

We drew 1,000 samples from the posterior distributions for seed survival and per capita reproductive success. For each of these samples, we calculated the optimal germination fraction $G$ by bootstrapping 100,000 values for $Y(t)$. Using the same sequence of years, but allowing the values of $Y(t)$ to vary with each draw from the posterior, we repeated this for each of the draws from the posterior. We used the same sequence of years across samples, isolating the effect of parameter uncertainty from the effect of sampling particular sequences of years (Evans et al. 2010). For example, if one sequence of resampled years was 1, 6, 3, 4, ... we repeated the analysis for each draw $i$ from the posterior:

- Draw 1: $Y_1(1)$, $Y_1(6)$, $Y_1(3)$, $Y_1(4)$, ...
- Draw 2: $Y_2(1)$, $Y_2(6)$, $Y_2(3)$, $Y_2(4)$, ...
- ...
- Draw 1000: $Y_{1000}(1)$, $Y_{1000}(6)$, $Y_{1000}(3)$, $Y_{1000}(4)$, ...

To assess the influence of parameter uncertainty on optimal germination fractions, we summarized the optimal $G$ calculated for each draw from the posterior (Fig. S20). The optimal germination fractions were robust to uncertainty in parameter estimates. In most populations, sampling from the posterior distributions had a minor effect on optimal $G$, especially for populations with high optimal germination fractions (e.g., LO). However, parameter uncertainty led to a broad range of optimal germination fractions in several populations. When we accounted for parameter uncertainty, the distribution of optimal germination fractions for five populations

(MC, FR, OSR, S22, and GCN) was centered at a higher value than the optimal germination fraction calculated from the posterior modes (compare the Os and Xs in Fig. S20). The broad range of optimal germination fractions in these populations is the product of parameter uncertainty that favors higher optimal germination fractions. For example, the posterior distribution for $s_0$ is asymmetric with a positive skew for several populations (Fig. S12) and greater values of $s_0$ favor higher optimal germination fractions.

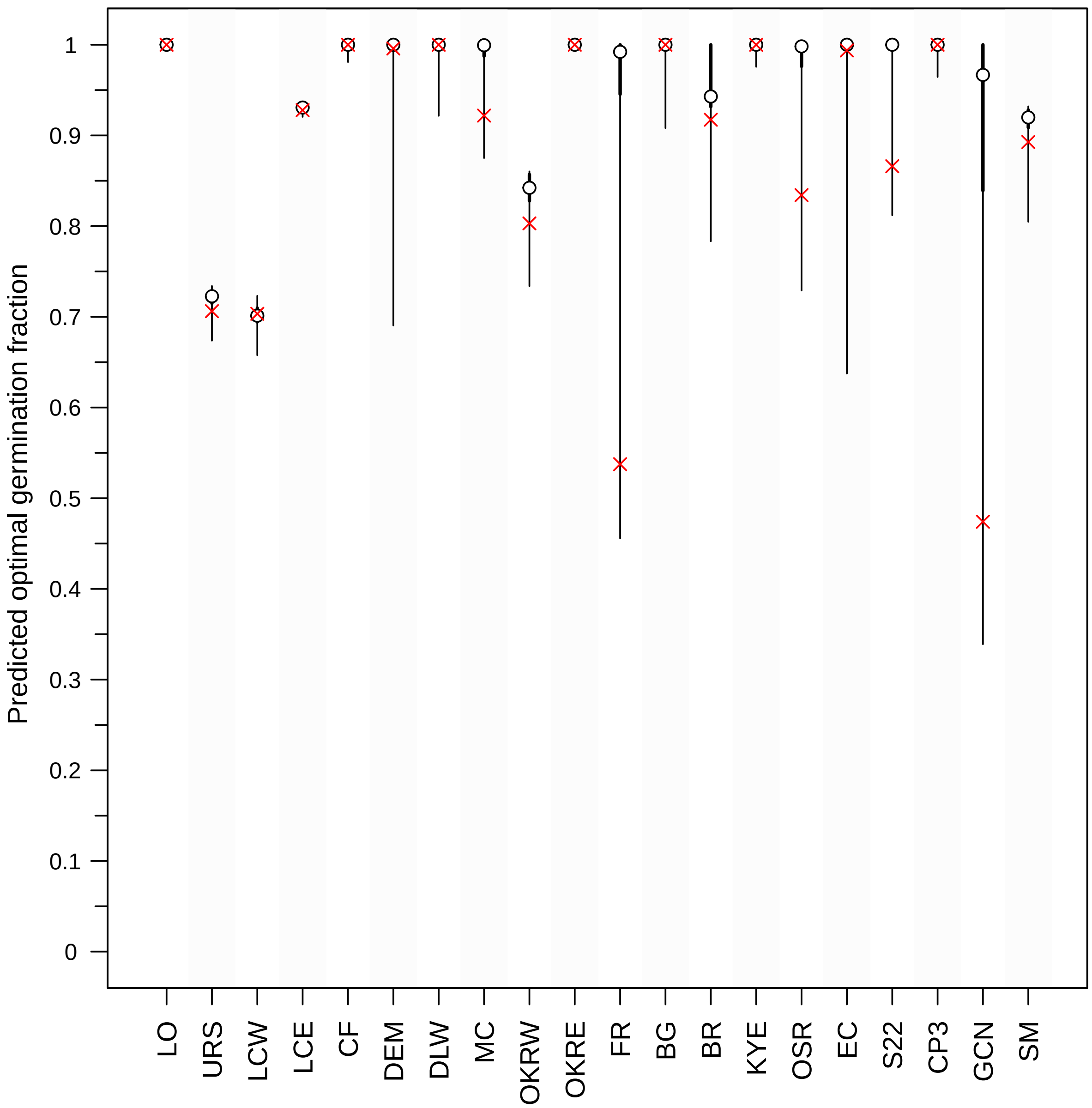

Figure S20: Influence of parameter uncertainty on predicted optimal germination fraction, G. The points are the posterior modes for optimal $G$ and the thick and thin error bars are, respectively, the 50% and 95% highest posterior density intervals. For each population, optimal $G$ was calculated for 1000 parameter sets that were randomly sampled from the full posterior. The red Xs show the optimal germination fraction in each population calculated using the posterior mode; these are the same values as those shown in Figure 4 in the main text. Populations are arrayed by easting from west to east, with the most western populations at the left and the most eastern populations at the right.

#### *S6.5 Exploratory analysis of environmental variability and per capita reproductive success*

##### *S6.5.1 Climate data*

A weather station network was established as part of the long-term study of *C. x.* demography (described in Eckhart et al. 2011). The network consists of 21 data loggers (Onset Computer Corporation) that recorded temperature and precipitation starting in October 2005; between 8 and 18 weather stations were actively recording throughout the study. Data from the network were used to spatially interpolate precipitation accumulation on a 1 hectare grid throughout the study area and estimate seasonal, cumulative precipitation (mm) at the study populations.

##### *S6.5.2 Relationship of reproductive success and growing season precipitation*

We did not observe a negative correlation between germination and the geometric standard deviation of per capita reproductive success (see **Results**). In our study, we do not make any explicit assumptions about the environmental variables, whether abiotic or biotic, responsible for the temporal variability in per capita reproductive success. However, precipitation is a major driver of temporal variability in plant demography (Compagnoni et al. 2021) and often assumed to be associated with reproductive success (but see Levine et al. 2008). We thus conducted an exploratory analysis of the relationship between growing season precipitation and reproductive success in the study populations. Our aim was not to conclusively identify the environmental drivers of demographic variability. Rather, we aimed to assess whether variability in precipitation during the growing season was sufficient to explain variability in reproductive success. Here, we define the growing season as February to June (i.e., after germination through to seed set).

We separately conducted a linear regression of the log of per capita reproductive success on the log of growing season precipitation (Venable 2007). As in Venable (2007), we added 0.5 to estimates for per capita reproductive success before log transformation (to include years with complete reproductive failure) and we added 1 to estimates of spring precipitation. For this exploratory analysis, we used the posterior mode as our point estimate of per capita reproductive success (as in the density-independent simulation). We applied a Bonferroni correction and assessed significance of our regressions at a confidence level of $p = 0.05/20 = 0.0025$.

We found that populations varied in how sensitive per capita reproductive success was to growing season precipitation (Fig. S21). The relationship between growing season precipitation and reproductive success was statistically significant in just one population (CF), after adjusting for multiple comparisons. The slope of the relationship also varied from close to zero (OKRE) to 3.4 (OKRW), indicating that sensitivity to spring precipitation varies among populations.

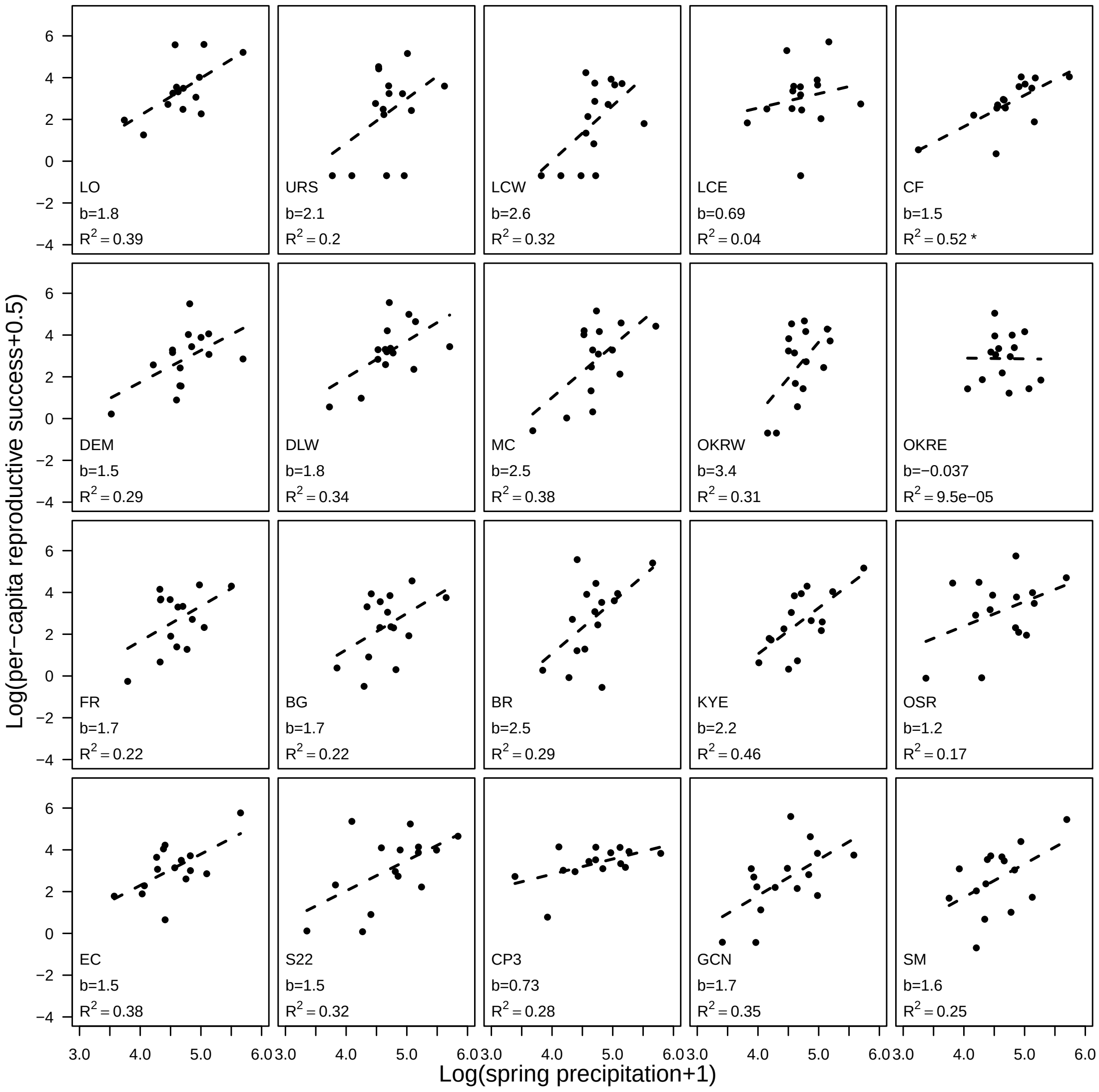

Figure S21: Log of per capita reproductive success (plus 0.5) plotted against the log of cumulative growing season (spring) precipitation (plus 1). Panels are arrayed by easting from west to east, with the most western populations at the top left and the most eastern populations at the bottom right.

#### S6.6 Seed mortality before and after germination have opposing effects on optimal germination

Pre- and post-germination seed mortality can have opposing effects on the evolution of delayed germination (Gremer and Venable 2014). Specifically, seed mortality after seed set, but before seeds have their first opportunity to germinate, discounts reproductive success and may thus favor delayed germination. Pre-germination seed mortality may include risks such as seed predation. On the other hand, for seeds in the seed bank, higher mortality makes it risky to remain in the seed bank and so selects against delayed germination.

We conducted an analysis to study the sensitivity of our results to our estimates of pre- and post-germination seed survival (the complement of seed mortality). First, we focused on the sensitivity of the optimal germination fractions to seed survival from seed production in July to October, $s_0$. Because our seed bag burial experiments started in October, we combined information from experiments and field surveys to estimate this parameter. We were thus interested in evaluating the impact of this parameter on optimal germination fractions. To compare the effect of seed survival before germination versus after germination, we also evaluated the sensitivity of optimal germination fractions to seed survival from after germination in January/February to the second October, $s_2$.

To evaluate the sensitivity of optimal germination fractions to seed survival, we calculated the optimal germination fraction across a range of potential seed survival values. For example, to calculate the sensitivity of optimal germination to $s_0$, we solved for the optimal germination fraction along an evenly spaced grid of possible values for $s_0$. For all parameters except $s_0$ and germination, we used the estimates from the field (in this case, $s_1$, $s_2$, $s_3$, and $Y(t)$ were all the posterior modes). For each population, this gave us a curve of potential optimal germination fractions associated with values for $s_0$ from 0 to 1. This curve shows the response of germination to the $s_0$. We then added the posterior mode and 68% highest posterior density interval of the estimate for $s_0$ to the curve, and highlighted the region of the curve for optimal germination fraction consistent with this uncertainty. We repeated this analysis with $s_0$ and $s_2$; in each case we varied the focal parameter and kept all other parameters at their posterior modes.

First, the results show that decreasing seed survival before germination favors lower optimal germination fractions (Fig. S22) while decreasing seed survival after germination favors higher optimal germination fractions (Fig. S23). Second, while this pattern is true in general, the sensitivity of optimal germination fractions to seed survival differs among populations. At one extreme, LO/LCE show little sensitivity to either $s_0$ or $s_2$; optimal germination fractions are 1 across a broad range for both parameters. At the other extreme, optimal germination fractions for MC/FR/S22/GCN are sensitive to both $s_0$ and $s_2$ across the whole range of 0 to 1. Third, combining the sensitivity analysis with parameter estimates shows that uncertainty about estimates for $s_2$ generally has a small effect on optimal germination fractions in most populations. However, FR and GCN both exhibit substantial sensitivity to estimates of $s_2$ (and $s_0$, which may help explain why optimal germination fractions for these populations are less robust to parameter uncertainty (Fig. S20). In contrast, the greater sensitivity to and larger uncertainty about $s_0$ translates to a larger range of optimal germination fractions that would be compatible with the data.

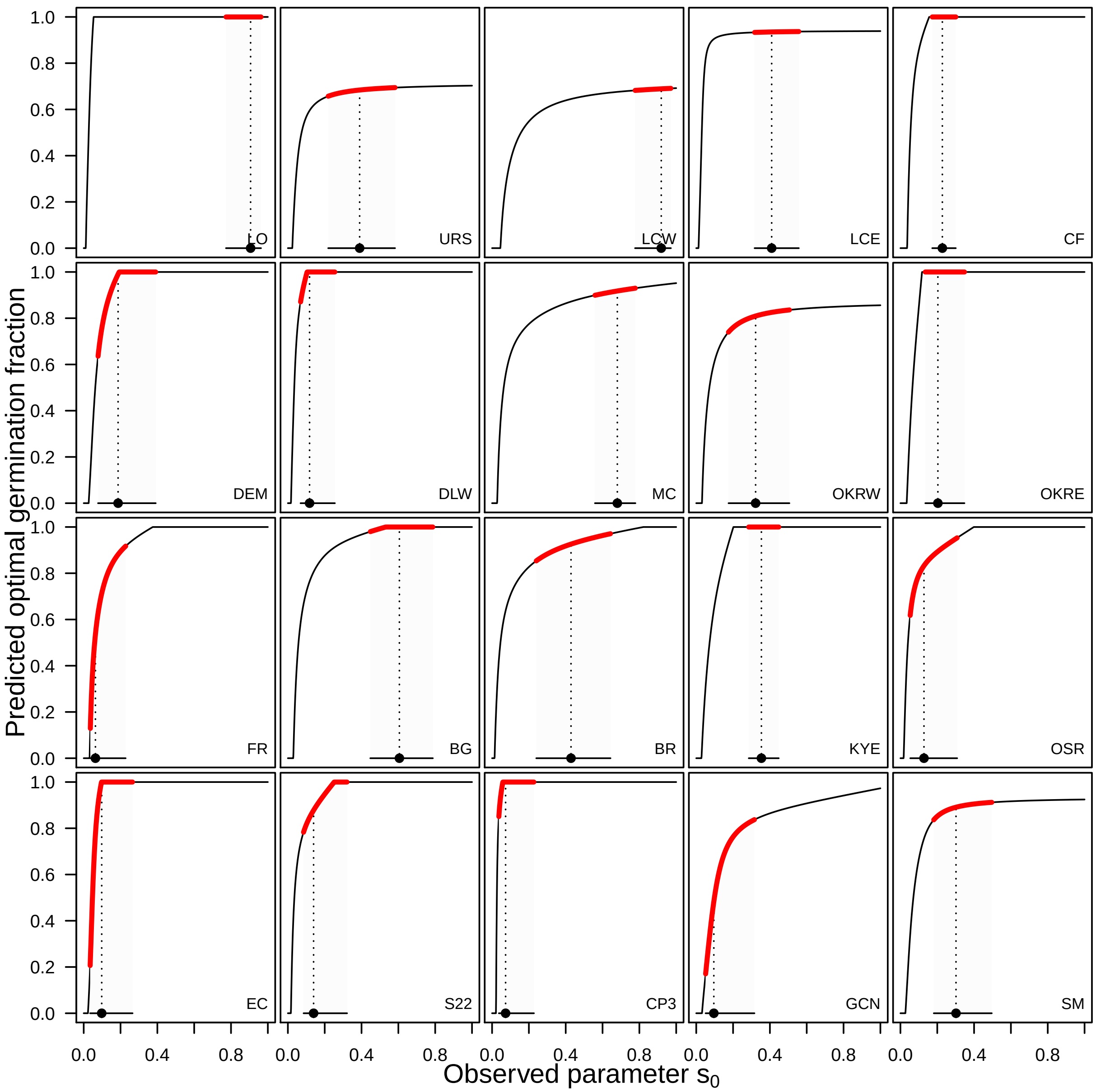

Figure S22: Sensitivity of the predicted optimal germination fraction to seed survival from seed production in July to October, $s_0$. For each population, the black line shows the optimal germination fraction for values of $s_0$ from 0 to 1. Estimates of $s_0$ are shown as the posterior mode and the 68% highest posterior density intervals (under a normal distribution, 68% of the distribution is within 1 standard deviation). The range of optimal germination fractions compatible with this uncertainty is displayed as a red line. Plots are arrayed by easting from west to east, with the most western populations at the top left and the most eastern populations at the bottom right.

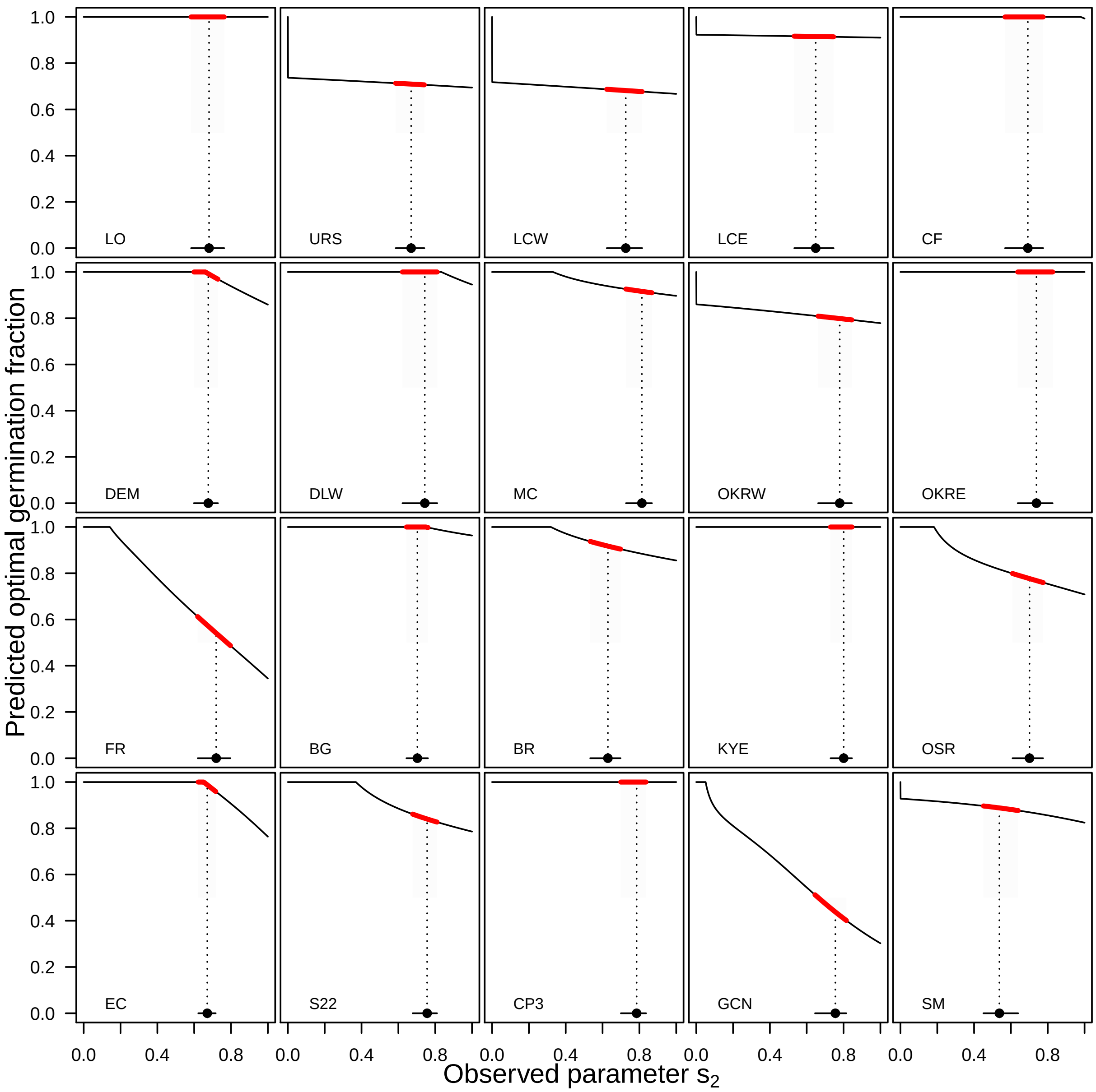

Figure S23: Sensitivity of the predicted optimal germination fraction to seed survival in the seed bank from January to October, $s_2$. For each population, the black line shows the optimal germination fraction for values of $s_2$ from 0 to 1. Estimates of $s_2$ are shown as the posterior mode and the 68% highest posterior density intervals (under a normal distribution, 68% of the distribution is within 1 standard deviation). The range of optimal germination fractions compatible with this uncertainty is displayed as a red line. Plots are arrayed by easting from west to east, with the most western populations at the top left and the most eastern populations at the bottom right.

#### *S6.7 Exploratory analysis of density-dependence in seedling establishment*

We used estimates for seed rain into permanent plots, estimates for seed mortality and germination from seed bag experiments, and observations of seedlings in permanent plots, to assess whether there is density-dependence at the early life stage of establishment. We consider establishment to be the transition between a seed in January and a seedling at the January/February census. Here, establishment is different than germination or emergence. Germination or emergence refer to transitions that we infer from the seed bags, while establishment refers to transitions from the seed bank to a seedling in permanent plots.

We combined observations and estimates of fruiting plants per plot, fruits per plant, and seeds per fruit to estimate seed production per plot from 2007-2020. For each population, we had 10 years of seedling counts that were each preceded by 3 years of seed rain estimates (e.g., seedling counts in 2010 were associated with seed rain estimates from 2007, 2008, and 2009). We used population-level (i.e., time-invariant) estimates for seed survival $s_0$, $s_1$, $s_2$, and $s_3$ to calculate the expected size of the seed bank, *s*seeds, in January of year $t$, just prior to the seedling census. For seeds produced just before the year $t$, in year $t-1$, we calculated the expected size of the seed bank as $(\text{seed rain}) \times (s_0 s_1)$. For seeds produced in year $t-2$, we calculated the expected size of the seed bank two years later as $(\text{seed rain}) \times (s_0 s_1 (1-g_1) s_2 s_3)$. For seeds produced in year $t-3$, we calculated the expected size of the seed bank two years later as $(\text{seed rain}) \times (s_0 s_1 (1-g_1) s_2 s_3 (1-g_1) * s_2 * s_3)$. These calculations make the following assumptions: germination probability does not vary through time; survival after the first germination opportunity is seasonal ($s_2$ or $s_3$) but does not exhibit interannual variability. To calculate the expected size of the seed bank for this exploratory analysis, we also used posterior modes of parameter estimates. This means that our estimate for the size of the seed bank does not include any parameter uncertainty; a full analysis would likely want to account for this.

We used these data to explore whether seedling establishment exhibits density-dependence. We used the analysis for quantifying intraspecific density-dependence in the seed to seedling transition described in the section *Results: Seedling-seed relationships* in Detto et al. (2019). Briefly, for each population, we fit an 'offset-power model':

$$\log(n_i^{\text{seedling}} + 1) = \log(a) + b\log(s_i^{\text{seeds}} + 1). \quad \text{(S24)}$$

The model is a log-transformed form of the power-law model relating transitions from one stage or size to the next ($N_0$ to $N_1$): $N_1 = aN_0^b$. For each population, we fit the model to pooled data on expected seed bank size ($s_i^{\text{seeds}}$) and observed seedling counts ($n_i^{\text{seedling}}$) in each plot $i$ from all 10 years. We used the estimates from the model to test the null hypothesis that the transition from seed to seedling does not exhibit density-dependence. Under this null hypothesis, the parameter $b$ is expected to be 1. If the transition exhibits density-dependence, the parameter $b$ is expected to be less than 1. To assess evidence for density-dependence, we subtract the null hypothesis, $b_{\text{Null}} = 1$, from the parameter estimate, $\hat{b}$. Negative values would be consistent with density-dependence in seedling establishment.

The offset-power model can be biased due to log transforming and adding 1 (Detto et al. 2019; Hille Ris Lambers et al. 2002) and due to spatial correlations between the location where seedlings were counted and seed rain was estimated (Detto et al. 2019). Both biases can cause $\hat{b}$ to be underestimated, which means that the strength of density-dependence is overestimated. We used the correction recommended by Detto et al. (2019) to check for the first form of bias but do not have the information to correct for the latter. Briefly, the correction compares $\hat{b}$ against a null hypothesis in which $b_{\text{Null}}$ is not 1, but is instead a distribution of parameter estimates obtained by re-fitting the power-offset model to simulated data. We use the average probability of seedling establishment simulate data in which emergence is not density-dependent. We re-fit the offset-power model to these simulations and obtained a distribution of slope estimates which then forms our null hypothesis, $b_{\text{Null}}$. To assess evidence for density-dependence, we subtract the null hypothesis, $b_{\text{Null}}$, from the parameter estimate, $\hat{b}$. A difference less than zero would be consistent with density-dependence in seedling establishment.

Our estimates were consistent with densvity-dependence during the transition from seed to seedling: all differences between $\hat{b}$ and $b_{\text{Null}}$ fall below 0, and confidence intervals do not overlap 0 (Fig. S24). We accounted for at least one source of possible error in our estimate of density-dependence; doing so reduced the estimated strength of density-dependence (red points are generally closer to 0 than black points). We interpret the results of this exploratory analysis as evidence that seedling establishment is possibly subject to density-dependence. For example, establishment may be limited by the availability of microsites (Eriksson and Ehrlén 1992). We were unable to account for other variables that might explain patterns of establishment or account for all known biases in estimating density-dependence. However, this exploratory analysis is, at the least, consistent with the hypothesis that density-dependence acts during seedling establishment.

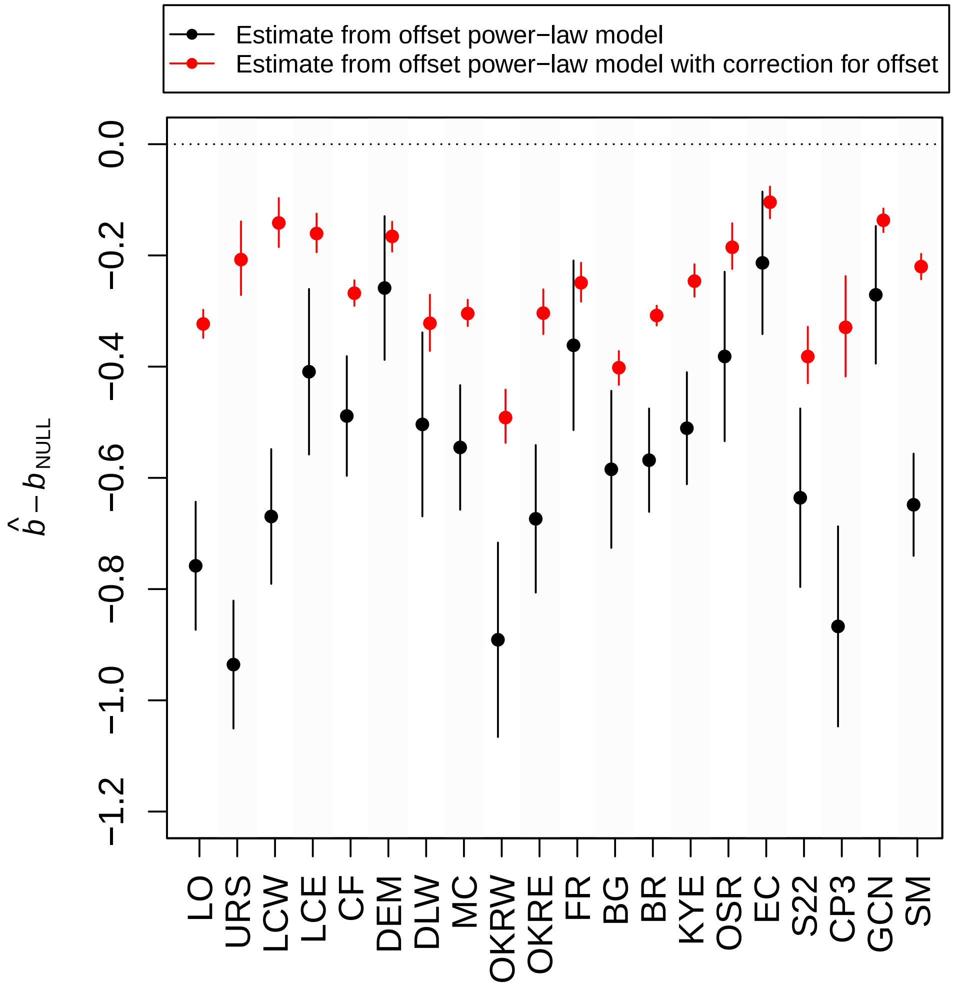

Figure S24: Evidence for intraspecific density-dependence in the seed to seedling transition, without (black points) and with (red points) a correction for bias introduced by using an offset of one in the power model. Estimates for $\hat{b}$ come from the slope parameter in the offset-power model, fit to 10 years of data on estimated seed bank sizes and seedling counts. The plot shows the difference between $\hat{b}$ and a null hypothesis, $b_{\mathrm{Null}}$; the null hypothesis is that there is no-density dependence and values less than zero are consistent with density-dependence. Error bars show the 95% confidence intervals. Without a correction for bias from the offset, $b_{\mathrm{Null}} = 1$. With a correction for bias from the offset, $b_{\mathrm{Null}}$ is calculated by refitting the offset-power model to simulated data generated by assuming an average transition probability.

#### S6.8 *Exploratory analysis of density-dependence in seedling survival to fruiting*

We conducted an exploratory analysis of density-dependence in seedling survival to fruiting using the observations of seedlings and fruiting plants. We used observations of seedlings (number of trials) and fruiting plants (number of successes) to estimate the probability of seedling survival to fruiting for each population $k$ with a logistic regression (Detto et al. 2019). First, we built a model (M0) that only included interannual variation in seedling survival:

$$\begin{aligned} y_{ij}^{\text{fruiting}} &\sim \text{binomial}(n_{ij}^{\text{seedling}}, p_{ij}) \\ \log(\frac{p_{ij}}{1-p_{ij}}) &= b_o + \epsilon_j. \end{aligned} \tag{S25}$$

We fit this model to observations from each population separately. In this model, $b_0$ is the population-level seedling survival, and $\epsilon_j$ is a random effect for year $j$.

Second, we built a model (M1) that included interannual variation and the effect of intraspecific density-dependence on seedling survival:

$$\begin{aligned} y_{ij}^{\text{fruiting}} &\sim \text{binomial}(n_{ij}^{\text{seedling}}, p_{ij}) \\ \log(\frac{p_{ij}}{1-p_{ij}}) &= b_o + b_1 n_{ij}^c + \epsilon_j. \end{aligned} \tag{S26}$$

In this model, $n_i j$ is the number of seedlings in plot $i$ in year $j$. The parameter $c$ controls the shape of the relationship between seedling density and survival. The parameter $c = 1$ defines a linear increase in the effect of seedling density on survival; values of $c$ less than 1 correspond to a model where the additional effect of increases in seedling density decline as the number of seedling grows (Detto et al. 2019). Finally, $\epsilon_j$ is again a random effect for year $j$.

We fit models M0 and M1 to observations from each population using maximum likelihood in the R package lme4 (Bates et al. 2015). For model M1, we first fit the models for values of $c$ between 0.01 and 1. We then calculated the value of $c$ that maximized the sum of the log-likelihood (equivalent to minimizing the negative log-likelihood) over the models fit to all populations (Detto et al. 2019). This allowed us to use the data to inform the shape of the functional relationship between seedling density and seedling survival. We subsequently used the value of $c$ that maximized the log-likelihood ($c = 0.3$; Fig. S25) to fit the model M1. We used likelihood ratio tests to test whether a model with (M1) or without (M0) density-dependence provided a better fit to the data from each population. Note that model M0 is a subset of model M1 with $b_1 = 0$ (i.e., no density-dependence). In assessing statistical significance, we applied a Bonferroni correction to account for the multiple comparisons that we made ($0.05/20 = 0.0025$).

In the majority of populations (16/20), the model with a term for density-dependence (M1) provided a better fit to the data (Table S12). In the remaining four populations (LCE, MC, SM, URS) the likelihood ratio tests did not suggest that the model with or without a term for density-dependence provided a better fit to the data.

We interpret the results of this exploratory analysis as suggesting that intraspecific density-dependence likely influences seedling survival to fruiting in many of the populations we studied. However, we note that we have counts of *C. x.* ssp. *xantiana* seedlings in our study plots, but that we do not have data on the density or biomass of other forbs or grasses, which may complicate interpretation of the patterns we report here. While this analysis suggests that density-dependence affects seedling survival to fruiting, there may be additional effects of density-dependence on other aspects of the life cycle, such as fruit number.

Explicitly incorporating density-dependence into our population models is beyond the scope of the present study. To test predictions of bet hedging with density-dependence, we will need to use methods from adaptive dynamics and calculate evolutionary stable strategies (ESSs) for germination rather than the optimal strategies we currently find in the manuscript. We expect that the results we present in this manuscript form a solid foundation for pursuing this as a promising future direction.

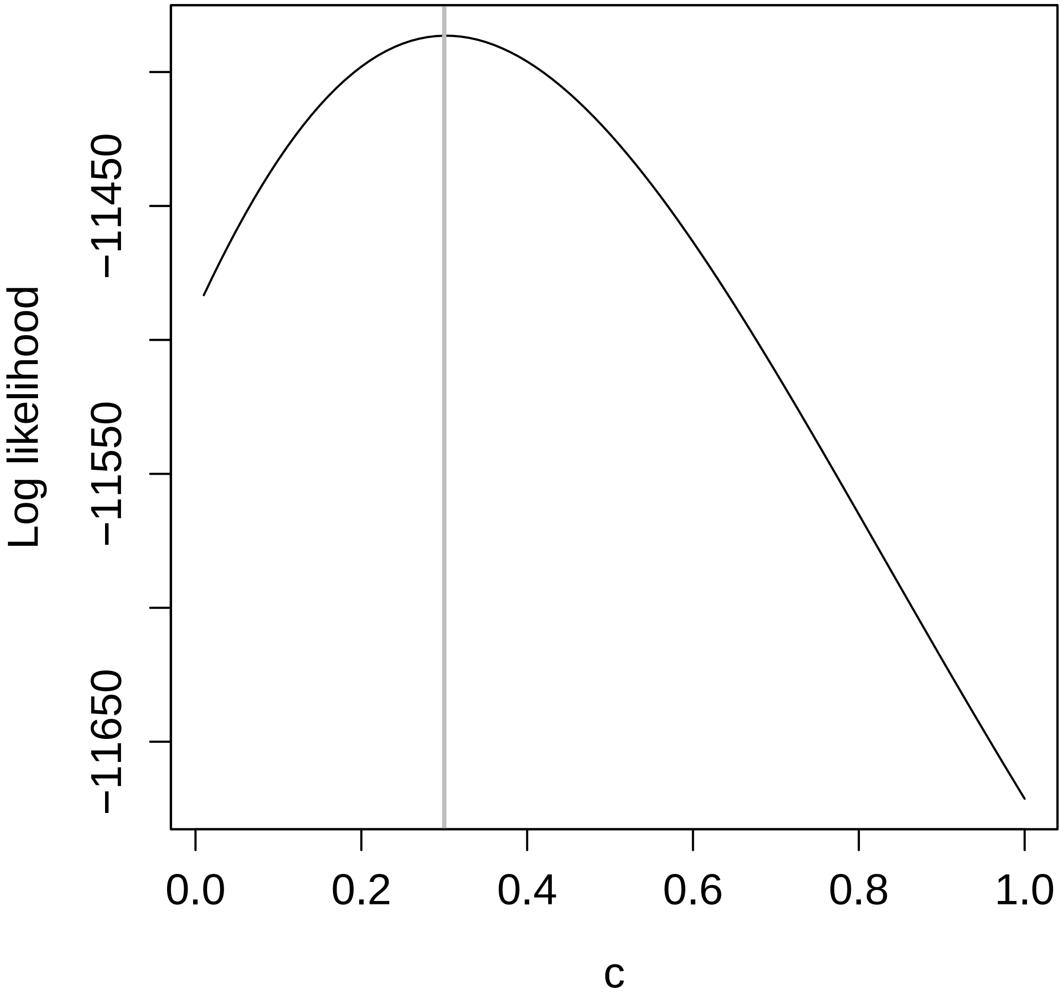

Figure S25: Log-likelihood for a model of intraspecific density-dependence in seedling survival to fruiting against the parameter $c$. The parameter $c$ is an exponent that modifies the effect of seedlings as seedling number increases. Values of $c = 1$ indicate a linear relationship, while values of $c < 1$ indicate a non-linear relationship. In other words, $c$ influences the shape of the relationship between seedling number and the effect of competition. We modeled density-dependence as a logistic regression, with seedling counts as trials and fruiting plant counts as successes. For each population, we refit the model over an even grid of $c$ values between zero and one. For each $c$, we then summed the log-likelihoods for all models. The value $c = 0.30$ maximized the log-likelihood (equivalent to minimizing the negative log-likelihood) and is shown as a vertical gray line. We subsequently used this value of $c$ to make inferences about density-dependence in seedling survival to fruiting.

Table S12: Likelihood ratio tests for logistic regressions of seedling survival to fruiting without (M0) and with (M1) density-dependence.

| Population | Log-likelihood (M0) | Log-likelihood (M1) | $\chi^2$ | p-value |
|---|---|---|---|---|
| LO | -653.45 | -572.06 | 162.79 | $< 10^{-4}$ |
| URS | -85.31 | -82.31 | 6.00 | 0.0143 |
| LCW | -723.53 | -541.29 | 364.48 | $< 10^{-4}$ |
| LCE | -348.36 | -344.60 | 7.53 | 0.0061 |
| CF | -748.98 | -671.97 | 154.02 | $< 10^{-4}$ |
| DEM | -1335.30 | -1268.10 | 134.37 | $< 10^{-4}$ |
| DLW | -307.55 | -277.50 | 60.09 | $< 10^{-4}$ |
| MC | -498.96 | -497.32 | 3.27 | 0.0704 |
| OKRW | -251.38 | -230.51 | 41.73 | $< 10^{-4}$ |
| OKRE | -261.28 | -235.87 | 50.82 | $< 10^{-4}$ |
| FR | -859.90 | -709.12 | 301.56 | $< 10^{-4}$ |
| BG | -808.21 | -746.46 | 123.50 | $< 10^{-4}$ |
| BR | -1622.20 | -1383.80 | 476.96 | $< 10^{-4}$ |
| KYE | -679.31 | -651.00 | 56.63 | $< 10^{-4}$ |
| OSR | -401.37 | -391.91 | 18.93 | $< 10^{-4}$ |
| EC | -1090.60 | -1038.60 | 104.08 | $< 10^{-4}$ |
| S22 | -380.86 | -346.14 | 69.44 | $< 10^{-4}$ |
| CP3 | -245.21 | -228.98 | 32.47 | $< 10^{-4}$ |
| GCN | -722.90 | -650.57 | 144.66 | $< 10^{-4}$ |
| SM | -521.44 | -518.41 | 6.06 | 0.0138 |

Likelihood ratio tests for logistic regressions for seedling survival to fruiting with density-dependence (M1) and a nested model without density-dependence (M0). For each population, the table presents the log-likelihoods of models M0 and M1. The critical $\chi^2$ value, is calculated as $-2$ times the difference between log-likelihood of M0 and M1. The p-value gives the probability that a $\chi^2$ distribution with 1 degree of freedom (the difference in number of parameters between M0 and M1) is greater than the critical $\chi^2$ value. A small difference in log-likelihoods between models M0 and M1 suggests including a term for density-dependence does not improve the fit of the models to the data. A larger difference in log-likelihoods between models M0 and M1 suggests including a term for density-dependence does improve the fit of the models to the data.

### S7 Glossary

**Experiments and data sources**: description of which experiments produce which data.

**Field experiments**: We use ‘field experiments’ to refer to the seed bag burial experiments conducted to study seed bank dynamics in the field.

**Lab trials**: We use ‘lab trials’ to refer to the lab germination trial and lab viability assays conducted to assess the viability of intact seeds.

**Lab germination trial**: In lab germination trial, we tested all or a subset of the intact seeds remaining in the seed bags when bags were recovered to the lab. Intact seeds were subject to a germination trial in the lab, in which intact seeds are induced to germinate. The germination trial produces observations of germinants.

**Lab viability assays**: In lab viability assays, we tested all or a subset of the seeds that did not germinate in the lab germination trial. Intact seeds that did not germinate were subject to a tetrazolium assay, which determined whether a seed was viable. The viability assay produces observations of viable seeds.

**Seed bag burial experiment**: In seed bag burial experiments, we added seeds and soil to mesh bags before burying them in the field. Bags are recovered from the field at different intervals and used for counts of intact seeds, seedlings, and/or lab trials. In the 2005-2008 seed bag burial experiments, we buried bags in Octobers and then counted seedlings and intact seeds in January 1, 2, or 3 years after bags were buried. Bags were then returned to the soil until October when intact seeds were again counted. Intact seeds were then used in a series of lab trials to assess the viability of intact seeds.

**Observations**: definitions of the observations.

**Intact seeds**: Seeds that remain intact in a seed bag. It is not possible to tell apart seeds that are intact and viable from seeds that are intact but dead in the field. Observations of intact seeds are produced by seed bag burial experiments.

**Seedlings**: Seedlings are young plants of indeterminate age that likely exhibit only cotelydons or first true leaves. Observations of seedlings are produced by seed bag burial experiments (when bags are surveyed in the winter).

**Seeds germinating**: When discussing the lab germination trials, we describe the number of seeds germinating. This refers to the seeds germinating in the lab trials, and distinguishes the counts of seeds germinating in the lab trials from the field experiment.

**Seeds staining**: When discussing the lab viability trials, we describe the number of seeds staining in the context of the lab viability assay. This refers to the observation of seeds staining red in the tetrazolium assay.

**Seed fates**: terms used to describe the processes that produce the observations.

**Emergence**: Emergence describes intact seeds becoming seedlings from the seed bank. Because intact seeds may not be viable, emergence describes the transition from intact seed to seedling but does not account for the proportion of intact seeds that are not viable.

**Germination**: Germination describes viable, intact seeds becoming seedlings from the seed bank. Germination thus accounts for the proportion of intact seeds that are not viable.

**Germination in the germination trial**: Germination in the germination trial describes germination of seeds that are tested in the lab germination test. To avoid confusion with seed fates in the field, we always refer to this by the full phrase.

**Persistence**: Persistence describes seeds remaining intact (but possibly not viable) in the seed bank. Persistence does not account for the proportion of intact seeds that are not viable.

**Emergence**: Emergence describes intact seeds becoming seedlings from the seed bank. Because intact seeds may not be viable, emergence describes the transition from intact seed to seedling but does not account for the proportion of intact seeds that are not viable.

**Survival**: Survival describes the viable, intact seeds remaining viable, intact seeds in the seed bank. Survival thus accounts for the proportion of intact seeds that are not viable.

**Viability in the viability assay**: Viability in the viability trial describes seeds being viable in the lab viability assay, conditional on not having germinated in the lab germination test. To avoid confusion with seed fates in the field, we always refer to this by the full phrase.

**Viability**: Viability describes intact seeds in the field being both intact and viable. Viability is calculated by combining estimates for germination in the germination trial and viability in the viability assay.
